# Single Cell Transcriptomics Reveals the Hidden Microbiomes of Human Tissues

**DOI:** 10.1101/2022.10.11.511790

**Authors:** Gita Mahmoudabadi, Sheela Crasta, Tabula Sapiens Consortium, Stephen R. Quake

## Abstract

The human microbiome has been studied extensively across those sites in the body that are readily accessible to sampling. Internal organs and tissues, however, have remained largely unexplored and, in the absence of infectious disease, are widely assumed to be free of microorganisms. Using single-cell transcriptomic data from the Tabula Sapiens spanning 11 human organ donors, 19 tissue types, 400,000+ annotated cells, 100+ cell types, and ∼70 billion sequences, we created an atlas of the human tissue microbiome with cell type resolution across three domains of life which we refer to as the Tabula Sapiens Microbiome. Together with data from 8 additional donors derived from a different set of experimental techniques for validation, we identified sequences from numerous bacterial, viral and fungal species in human tissues. We mapped the likely microbial flow routes in the body from external-facing microbiomes and natural environments to internal tissues and tumors, demonstrating the existence of several unexpected routes. In the case of tumor microbiomes, we find a considerable number of bacterial genera found in tumors are detectable across tumor-free tissues, suggesting that tumor microbiomes are in part sourced from adjacent normal tissues, even those from distant sites. Finally, by increasing the resolution of sampling from tissues to cell types, we quantified the microbial load and diversity across different human cell types to reveal a network of host cell type and microbe associations. For example, we identified traces of both latent and active Epstein Barr Virus infections in various cell types such as splenic plasma cells. Broad exploration of the human tissue microbiomes may provide insights which ultimately are of clinical importance.

## Introduction

Microbial life contains a vast repertoire of biochemical pathways that enable it to colonize even the most extreme environments on the planet. It is no wonder that the human body, an energy-rich environment for trillions of our own cells, would be host to many highly diverse microbes^1, 2^. Historically, understanding of microbes in connection to human health has emerged from waves of encounters with devastating pathogens such as *Mycobacterium tuberculosis*, *Vibrio cholerae*, and most recently, SARS-CoV2. In the last two decades our view of human-microbe interactions has been transformed through increasingly powerful sequencing technologies which have enabled broad surveys of the microbial world^3–8^. As such, previously held dogma has been upended; one such example of this is the assertion that internal organs such as the healthy human lung are sterile environments^9^.

The study of the human microbiome has seen an explosion of data connecting communities of microbes or individual species to a variety of metabolic^10, 11^, neurological^12, 13^, cardiovascular^14, 15^, and autoimmune^16, 17^ diseases, among others. Most microbiome studies, however, have delineated microbial communities at sites easily accessible to sampling such as the human intestinal tract via feces, the mouth, and the skin. Less accessible internal sites, on the other hand, are largely assumed to be free of microorganisms, as they are protected by layers of epithelial and endothelial tissue. However, a few recent studies have challenged this assumption. For example, human tumors were shown to contain tumor-specific microbial sequences, providing a possible diagnostic approach for cancer detection^18^. Other studies similarly identified the presence of tumor-specific mycobiomes and microbiomes including intracellular bacteria^19–24^. Microbiomes in both tumor-free and tumor-derived breast, esophageal, oropharyngeal, gastrointestinal, and colorectal tissues have also been shown^25–27^.

We assembled the first atlas of human tissue and cell type microbiomes by mining the raw sequences from one of the largest single-cell transcriptomic datasets of human tissues called the Tabula Sapiens^28^ which encompasses 400,000+ annotated human cells, spanning 100+ cell types and 19 tissues, originating from 11 human organ donors. We identified high-confidence viral, bacterial, and fungal sequences originating from human single-cell transcriptomes, together forming the Tabula Sapiens Microbiome (TSM) dataset. Using a series of experimental and bioinformatic filters including a machine learning model for decontamination and a Large Language Model for habitat classification, we rigorously decontaminated and quality controlled this dataset.

Moreover, we combined our data with 16 external datasets (**SI Figure 1.a-b**), 10 of which were used for creating and benchmarking a pipeline for the identification of microbial sequences. Some examples of these databases include the Human Gastrointestinal Bacteria Culture Collection (HBC)^29^, the expanded Human Oral Microbiome Dataset (HOMD)^30^, Human Microbiome Project (HMP)^4^, and others. The other external databases, including Narunsky-Haziza *et al*.^20^ (tumor mycobiomes), Nejman *et al.* (tumor microbiomes)^21^, the Unified Human Gastrointestinal Genome^5^ (UHGG) collection, Genomes from Earth’s Microbiomes (GEM)^7^, Pathosystems Resource Integration Center (PATRIC)^31^, and others were used for decontamination and analysis. For example, we explored the intersection of species in TSM tissues and those found at tumor sites and at sites accessible to sampling, or what we will refer to as “external-facing microbiomes” (e.g. the gastrointestinal, oral and skin microbiomes), as well as natural environments, to chart out possible microbial flow routes throughout the body. We report the inter-and intra-donor variability of tissue microbiomes and characterize microbial load and diversity across human cell types. Moreover, using an additional cohort of 8 donors, we devised an assay to extract virus-like and bacterial-like particles from human tissues and explored them using imaging and DNA sequencing. We will refer to this dataset as the Extracted Human Tissue Microbiome (EHTM) dataset. We further show that the human tissue and cellular microbiomes, beyond cell-free or capsid-free nucleic acids, may also include intact organisms.

## Results

### Computational pipeline for identification of microbial sequences from multiple biomes in single-cell datasets

We created a computational pipeline written in Snakemake^32^ to identify viral, bacterial and fungal sequences in single-cell transcriptomic data. Briefly, we use UMI-tools^33^ to filter and extract cell barcodes and UMIs from fastqs, and then use STAR^34^ to align resulting sequences against a reference library containing the human genome and ERCCs. After the removal of low-complexity sequences from the unmapped reads using a custom script, we use BLASTn^35^ to align reads against several custom databases followed by the nt database, which is one of largest publicly available databases. We refer to this pipeline as SIMBA. For a more detailed description of this pipeline see **SI: SIMBA, a computational pipeline for the identification of bacteria, viruses and fungi in single cells,** along with **SI Figure 1.a.** We also benchmarked this pipeline against several truth datasets to show that it is highly precise (see **SI: Benchmarking SIMBA’s precision and recall** as well as **SI Tables 1-8).** We provide a full explanation for why we chose not to rely on faster aligners or smaller reference databases given their higher false positive rates in samples that have low ratios of non-host to host sequences (See **SI: SIMBA, a computational pipeline for the identification of bacteria, viruses and fungi in single cells,** as well as **SI: Benchmarking Kraken2 precision and recall,** as we all as **SI Tables 9-11** and **SI Figure 2)**. A common theme in our study is that in the tug-of-war between precision and recall, we choose precision as we decided it was more important to us to identify microbial reads precisely and with high confidence than it was to capture more of them.

### Decontamination and quality filtering of the Tabula Sapiens Microbiome dataset

In the previous section and its supplementary material, we address a major source of false positives in microbial datasets, which are false positives generated during alignments. Another important source of false positives is physical contamination. We discuss in depth the measures taken during tissue procurement and transport to prevent potential contaminants (see **SI: Measures against contamination during tissue recovery and transport)**. We acquired samples of reagents and instruments that were used to process tissues and sequence single cells. Even when the same reagent was used across different labs, we obtained each lab’s specific sample of that reagent to account for lab-specific contaminants. Following the same protocols as the original experiments, we deeply sequenced more than 200 reagent and instrument controls using five 10X lanes (**SI Table 12**), and eliminated contaminant species (filter 1). We then eliminated putative bacterial (filter 2) and fungal (filter 3) contaminants previously identified by Salter *et al.*^36^ and Narunsky-Haziza *et al*^20^. For a detailed description of these three filters and their impact on the dataset, see **SI: Identification of contaminants based on negative controls and external datasets** and **SI Tables 12-13**.

To further account for any potentially missed contaminants, we built 6 types of machine learning models to learn the shared features of the known and putative contaminants acquired from the first three filters. As our fourth filter, we applied the best performing model type, which was the gradient boosting classifier (test AUPR and AUROC 95% confidence intervals were 0.75-0.78 and 0.82-.84, respectively) to identify putative contaminants in the portion of the dataset that was withheld from both training and testing (see **SI: Machine learning models of contamination, SI Tables 14-15** and **SI Figure 1.c-d**). Additionally, we observed a bimodal distribution of alignments and removed species that are represented primarily by the lower-quality alignment peak, which served as the 5th filter (see **SI: Gaussian Mixture Model of alignments,** and **SI Figure 3).** To further verify our findings, we imposed a 6th filter which selects taxa that have been previously identified in the human microbiome by three large studies, namely **H**MP, **U**HGG, or G**E**M datasets, or what we abbreviate as HUE studies (see **SI: Identifying putative habitats of origin for 500+ bacterial genera in the Tabula Sapiens Microbial dataset using the Genomes from Earth’s Microbiomes, the Human Microbiome Project, and the Unified Human Gastrointestinal Genome collection** and **SI Figure 4)**. For the remaining taxa that had not appeared in HUE studies (7th filter), we benchmarked (precision of 98% and a recall of 75%) and programmatically employed openAI’s Davinci Large Language Model to further select human-associated taxa (see **SI: Benchmarking the Davinci large language model for classification of microbial species based on their habitat, SI Tables 16-18** and **SI Figure 1.e**). Habitat-based filters were placed to select commonly known human-resident or human-associated microbes from the rest.

The output of SIMBA that has been filtered to contain only sequences from the three biomes of interest is referred to as the Tabula Sapiens Microbiome (TSM) dataset F0, to signify that it has not been filtered based on contaminants. The dataset filtered only by the first filter is referred to as TSM F1. The dataset filtered by the first two filters is referred to as TSM F2 and so on, such that TSM F7 is the dataset filtered by all seven filters. The impact of each filter on the total number of microbial hits, species, genera, cell-containing droplets and empty droplets with significant hits is shown in **SI Table 12** and **SI Figure 5**. With the imposition of all filters, we arrive at ∼0.4% of the unique hits found in the TSM F0 dataset, ∼3% of the species and ∼5% of the genera. Like the percentage of unique hits remaining, the percentage of cell-containing and empty droplets is ∼0.7% and ∼0.4%, respectively. Because the first five filters represent decontamination and quality control filters, we will generally report our findings on TSM F5 and TSM F7, which is the most conservative version of the TSM dataset.

### The Tabula Sapiens Microbiome atlas reveals numerous species of microbes spanning viruses, fungi and bacteria

Binning sequences by donor and tissue type, we summarize their breakdown into high-level taxonomic categories **(Figure 1)**. Numerous species spanning three domains of life are present across 11 donors and 19 tissue types, with bacteria having the highest prevalence of the three domains. We show the fraction of microbial hits in TSM F5 and F7 datasets that have resolved taxonomic identity at each classification level and demonstrate a high percentage, typically >95% (**SI Table 19**). Unlike bacterial sequences, which map largely to commensal species, viral sequences mapped mostly to the Epstein Barr Virus (EBV) - a ubiquitous and persistent human pathogen with very limited transcriptional footprint during its latency compared to its reactivation^37, 38^. As we will discuss in more detail, this dichotomy in the transcriptional load of EBV during its latency and reactivation likely gives rise to detectable transcripts in some donors while appearing to be missing from others. Finally, fungal sequences are generally present with lower relative abundances compared to bacteria. This is consistent with previous estimates of fungi at external-facing microbiomes, depicting them significantly outnumbered by bacteria^39–41^.

**Figure 1.**
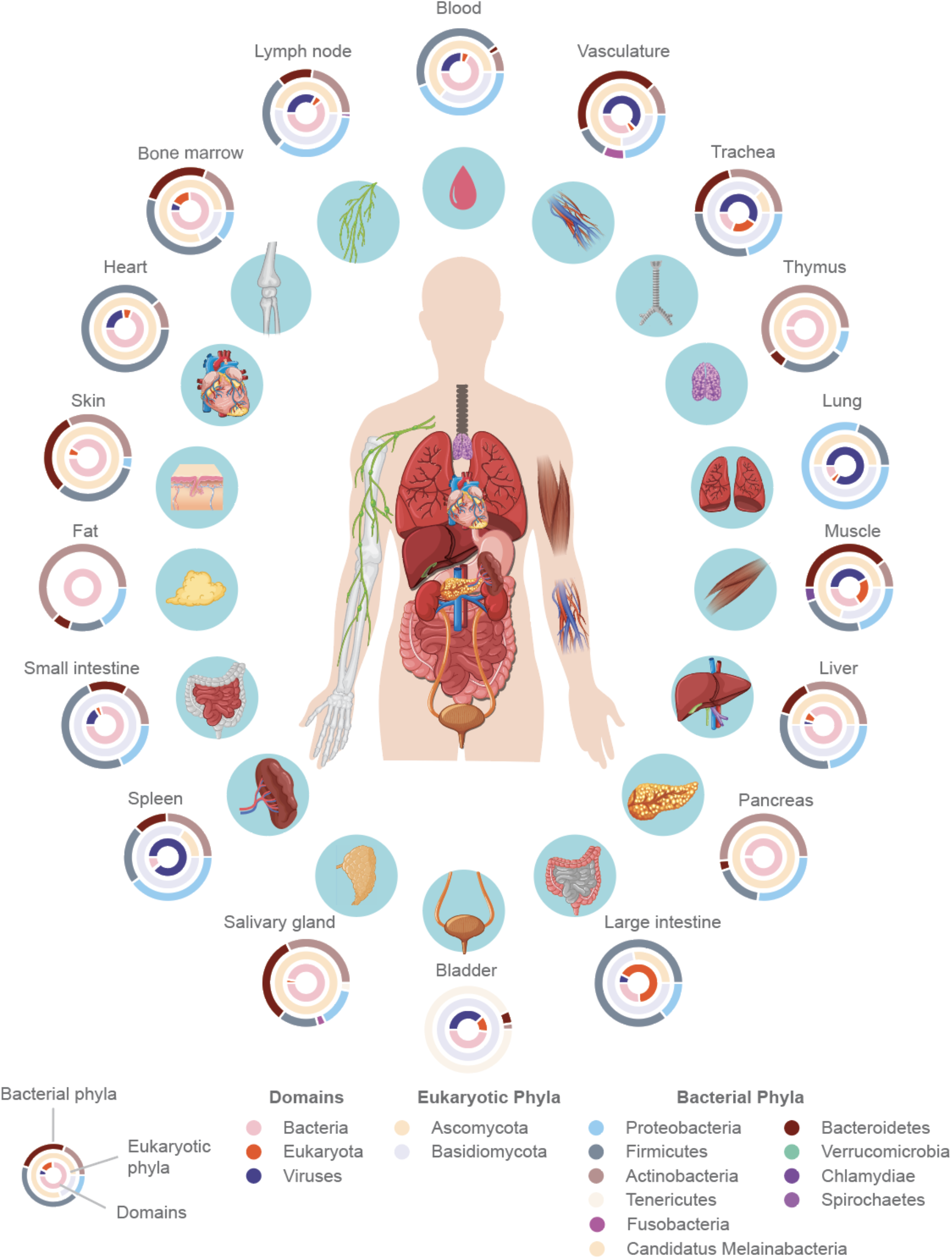
Tabula Sapiens Microbiome (TSM) atlas derived from the Tabula Sapiens single-cell transcriptomic dataset consisting of 19 tissue types from 11 donors (identified by TSP#), 100+ cell types and 400,000+ annotated cells. Tissue microbiomes are stratified based on taxonomy: domains (inner ring) followed by eukaryotic phyla, and bacterial phyla, respectively. Sequences are grouped based on donor (one donor is shown per tissue). Most tissues shown are from TSP2, except for Thymus, Liver, Salivary gland which are from TSP14, Pancreas from TSP9, Heart from TSP12, and Skin and Fat from TSP10. Tissues shown have at least 10 hits and are from the TSM F7 dataset.

Bacterial sequences are derived from a range of phyla, most notably Actinobacteria, Firmicutes, and Proteobacteria (**Figure 1**). We find that more than a quarter of all transcripts map bacterial ribosomal RNA (rRNA) (**SI Figure 6**). This relatively large percentage of ribosomal sequences is expected as rRNA is the most abundant form of RNA in bacterial cells, and its abundance is tightly coupled to the bacterial growth rate^42, 43^, accounting for up to 85% of the RNA pool in rapidly growing bacterial cells in culture^44, 45^. Furthermore, our finding of bacterial rRNA from diverse bacterial taxa across numerous human tissue types (**SI Figure 7)** is in agreement with a recent study of several tumor types, wherein 16S rRNA was detected using both sequencing and imaging^21^.

The single cell RNA sequencing protocols such as the one used in this study rely on a polyA selection. Hence, at first glance, the discovery of any microbial transcripts is surprising. However, many families of viruses have polyadenylated transcripts^46, 47^. Additionally, polyadenylation plays a ubiquitous yet understudied role in prokaryotic transcription^48, 49^. Moreover, it is well known that oligo-dT primers often bind with similar efficiencies to A-rich regions as they do to polyadenylated tails, and thus mispriming events inadvertently provide a window into non-polyadenylated transcripts^50, 51^.

### Mapping the possible microbial flow routes from external-facing microbiomes and natural environments to internal tissues and tumors

To explore the human tissue microbiome at a more granular level, we created a Sankey plot (**Figure 2.a**), wherein each one of the microbial hits is represented by a line, the path of which can be traced to obtain the tissue type of origin, taxonomic categorization, as well as additional information gathered from external databases. We provide this as an interactive plot (**SI File 1**) along with the table underlying these flow paths (**SI Table 20**). The HMP dataset was used to identify species that had previously been observed at external-facing microbiomes from the gastrointestinal tract, oral cavity, ear, nose, airways, urogenital tract, and the skin. Approximately 50% of bacterial genera that make up tissue microbiomes appear in external-facing microbiomes (**SI Figure 8**).

**Figure 2.**
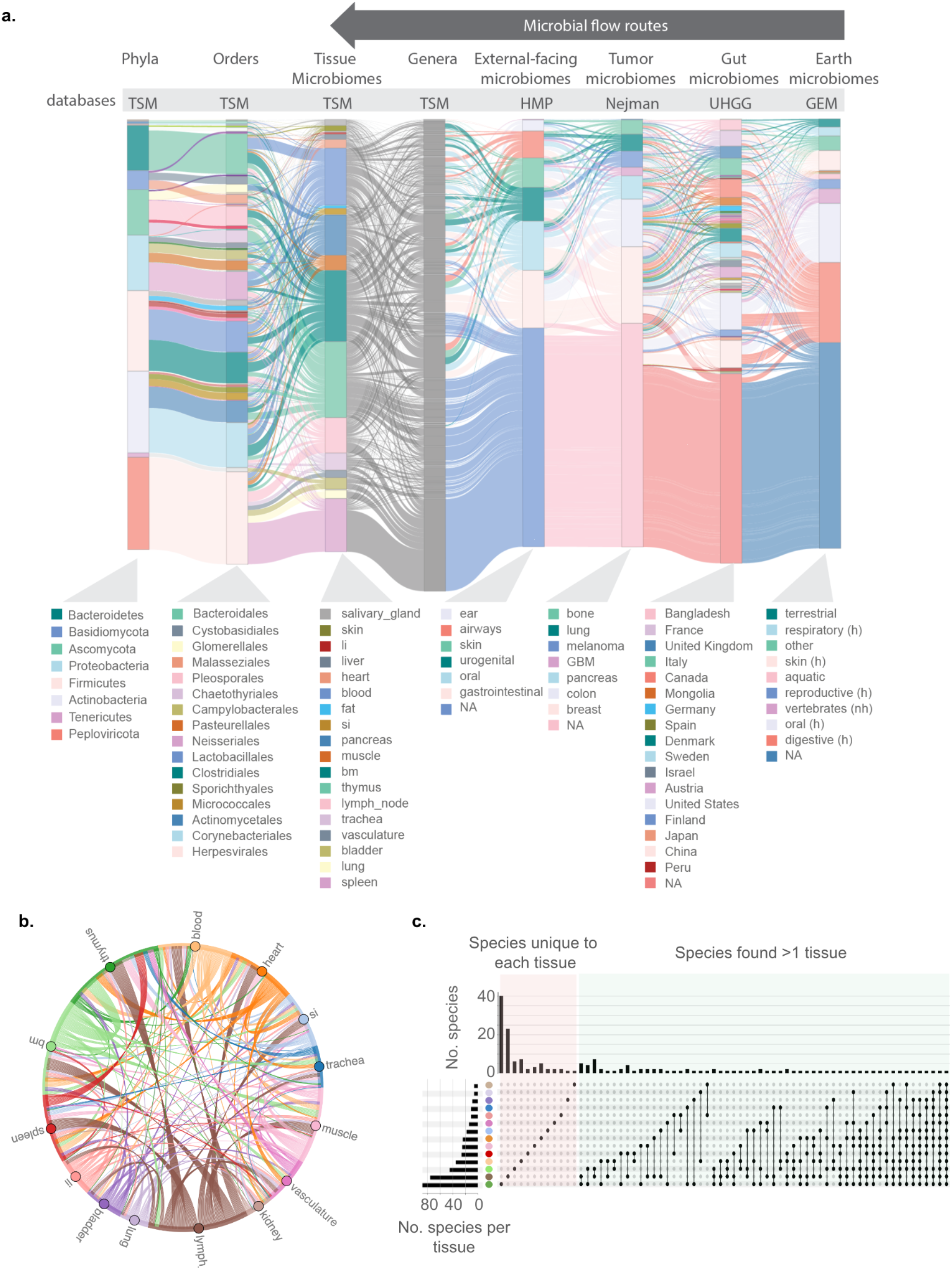
Microbial flow within the body from external and internal environments (TSM F7 dataset). a) Sankey plot revealing the taxonomic classification and tissue of origin for each sequence (each line in the plot represents an individual sequence) along with additional data from the Human Microbiome Project (HMP) database, tumor microbiome dataset by Nejman *et al.*, the Unified Human Gastrointestinal Genome (UHGG) collection and the Genomes from Earth’s Microbiomes (GEM). Sequences belonging to species that are not found in these additional databases, including all viral and fungal species, are labeled as “NA”. Under the GEM column, “h” denotes human and “nh” denotes non-human. Only some of the largest groups in each column are denoted in the legend. The rest can be found as part of the interactive plots provided in the **SI File 1**. b) Circos plot of shared species within TSP2 (donor 2). Each link represents a unique species. Species that are not shared between tissues are not shown. c) UpSet plot for TSP2 demonstrating the number of shared species between tissues, as well as the number of species unique to each tissue. Unique sets of species are shown as filled circles. The intersection of two or more sets are shown via connected filled circles. Bars denote the size of each set or the intersection of sets.Tissues are color-coded according to panel b. bm: bone marrow, li: large intestine, si: small intestine.

With the assumption that microbial flow occurs from external to internal sites, we depict possible flow routes for numerous bacterial genera and show that various external and internal sites are connected through their shared taxa (**Figure 2.a**, **SI Figure 9**). Surprisingly, not all species found at internal sites originate from the gastrointestinal tract. Other external-facing microbiomes appear to contribute many species to internal organs and tissues. To illustrate a few microbial flow paths, we detected sequences from *Treponema denticola,* an oral bacterium commonly associated with periodontal disease, in the heart. This observation is in line with studies that have linked *Treponema denticola* to atherosclerosis through its invasion of aortic tissue^52–54^. Sequences mapping to *Aggregatibacter aphrophilus*, another oral bacterium whose infection can lead to endocarditis, was also detected in the heart (**SI Figure 9**). While various members of the oral microbiome have been associated with cardiovascular diseases^55^, we find that the heart is certainly not the only destination for sequences from species normally found in the oral cavity. For example, we found sequences mapping to *Porphyromonas gingivalis* not only in the salivary gland tissue as might be expected, but also in tissues from the thymus, liver, vasculature, fat, bone marrow and others. Sequences mapping to inhabitants of other external-facing microbiomes were similarly found in various tissues (**Figure 2.a**, **SI Figure 9**). It remains to be discovered to what extent species from external-facing microbiomes can colonize these distant sites.

In addition to the HMP dataset, we compared the number of shared genera between TSM datasets and UHGG, which represents a much larger number of gut metagenomic samples than found in HMP (**Figure 2.a, SI Figure 10**) and includes samples collected from various countries, thus providing a global perspective on the shared genera. We further explored the bacterial taxa shared between TSM and tumors (Nejman *et al.*^21^) of the pancreas, breast, colon, lung, skin, ovaries, bone, and the brain (**Figure 2.a, SI Figures 11-14**). Nearly half of TSM F7 bacterial genera have previously appeared in human tumors (**SI Figure 11**). We created an abundance heatmap of these genera across TSM tissues (**SI Figure 11**). Sequences from species that were previously identified as highly enriched in tumors including *Paracoccus marcusii* and *Roseomonas mucosa* are also present across various TSM F5 tissues, primarily from lymphatic organs (**SI Figure 12**). We do not identify any of the tumor-enriched species in the TSM F7 dataset. The relatively large overlap between TSM and tumor microbiomes suggests that tumor microbiomes may be sourced from adjacent tumor-free tissue microbiomes, even possibly from tissues at distant sites from the tumor (**SI Figure 12-14**). We provide a list of bacterial genera found in tumor microbiomes that are missing from the TSM F5 or F7 datasets (**SI Table 21**). Some of these genera may perhaps prove to be tumor-microbiome specific, however future studies applying the same experimental and bioinformatic protocols to tumor-free and tumor-derived tissues are needed.

We mapped the possible microbial flow routes from natural environments and external-facing microbiomes to tissues and tumors by focusing on sequences from species found across TSM, GEM, HMP, UHGG, and the Nejman *et al.* datasets (**SI File 2, SI Table 22**). Many interesting flow routes appear to exist with varying degrees of complexity. As an example of a flow route, sequences from members of the *Aggregatibacter* genus appear to originate from the oral microbiome and the gastrointestinal tract (HMP, GEM, UHGG). They are detected in salivary gland tissue as well as thymus, blood, liver, heart, lymph nodes and other tissues (TSM F7). They are also found in breast tumors (Nejman *et al.*). We provide these flow routes both as an interactive plot (**SI File 2**) and as a table (**SI Table 22)**.

Additionally, we used the PATRIC database^31^ to obtain further information about the pathogenicity potential of bacterial species. We found the following potential bacterial pathogens in the TSM F7 dataset: *Porphyromonas gingivalis, Campylobacter showae, Aggregatibacter actinomycetemcomitans, Capnocytophaga gingivalis, Campylobacter curvus,* and *Treponema denticola*. From this list, the HMP dataset contains all but one pathogen, namely *Campylobacter curvus*. *Campylobacter curvus* and *Porphyromonas gingivalis* appear in human gut samples from the UHGG dataset. Finally, *Capnocytophaga gingivalis* appears in the Nejman et al. dataset.

To explore the inter-and-intra donor tissue microbiomes at finer taxonomic resolution, we computed the shared species within and across individuals. Whether considering the tissues of one donor or the collective tissue microbiomes of many donors, tissue microbiomes appear to have many overlapping species **(Figure 2.b, SI Figure 15)**. However, when accounting for all species, species unique to each sample outnumbered those that are shared (**Figure 2.c, SI Figures 16-17**). Furthermore, we compared species-level bacterial microbiomes in a pairwise fashion using the Spearman correlation for both TSM F7 and F5 datasets (**SI Figures 18-19**), revealing tissue microbiomes across and within each individual to be uncorrelated at the species level, but increasingly more similar at higher levels of classification. For a more detailed discussion, see **SI: Exploring the inter-and-intra tissue microbiomes.**

### Increasing the resolution of tissue microbiomes to delineate cell type microbiomes

Many of the microbial sequences can be traced back to cells that have been annotated by the Tabula Sapiens Consortium. Merging cell type annotation data from Tabula Sapiens objects with cellular barcodes provides an unprecedented view of each tissue’s microbiome at the level of cell types and even individual cells. We wondered how different cell types compare in terms of their microbial load. Thus, we quantified single-cell microbial load as log_2_(h_c_/g_c_), where h_c_ represents the number of microbial sequences detected in cell *c* and is normalized by the number of host genes detected in that cell, g_c_, to account for cell to cell variation in sequencing coverage. It is important to note that this data does not indicate whether these microbes or their sequences have been internalized by these cells; they could be binding to the external cell membrane.

Ranked distributions of single-cell microbial loads (**Figure 3.a**), reveal that cell types with the largest microbial load include enterocytes, T cells, and plasma B, among other cell types, especially those from the immune compartment. The over-representation of immune cell types is expected as these cells, with the ability to survey tissues, are the first to encounter microbial organisms. Enterocytes are also the first line of barrier against the trillions of microbes housed in the human intestinal tract. At first glance, the finding of microbial sequences in pancreatic acinar cells and in the pancreas more generally, may be the most surprising observation, however the pancreas is an organ with a physical connection to the small intestine. This connection has been previously shown to enable the passage of intestinal fungi to the pancreas where they can promote pancreatic adenocarcinoma^24^. Additionally, pancreatic adenocarcinoma tumors have been shown to harbor microbiomes that are predictive of long-term survival^22^.

**Figure 3.**
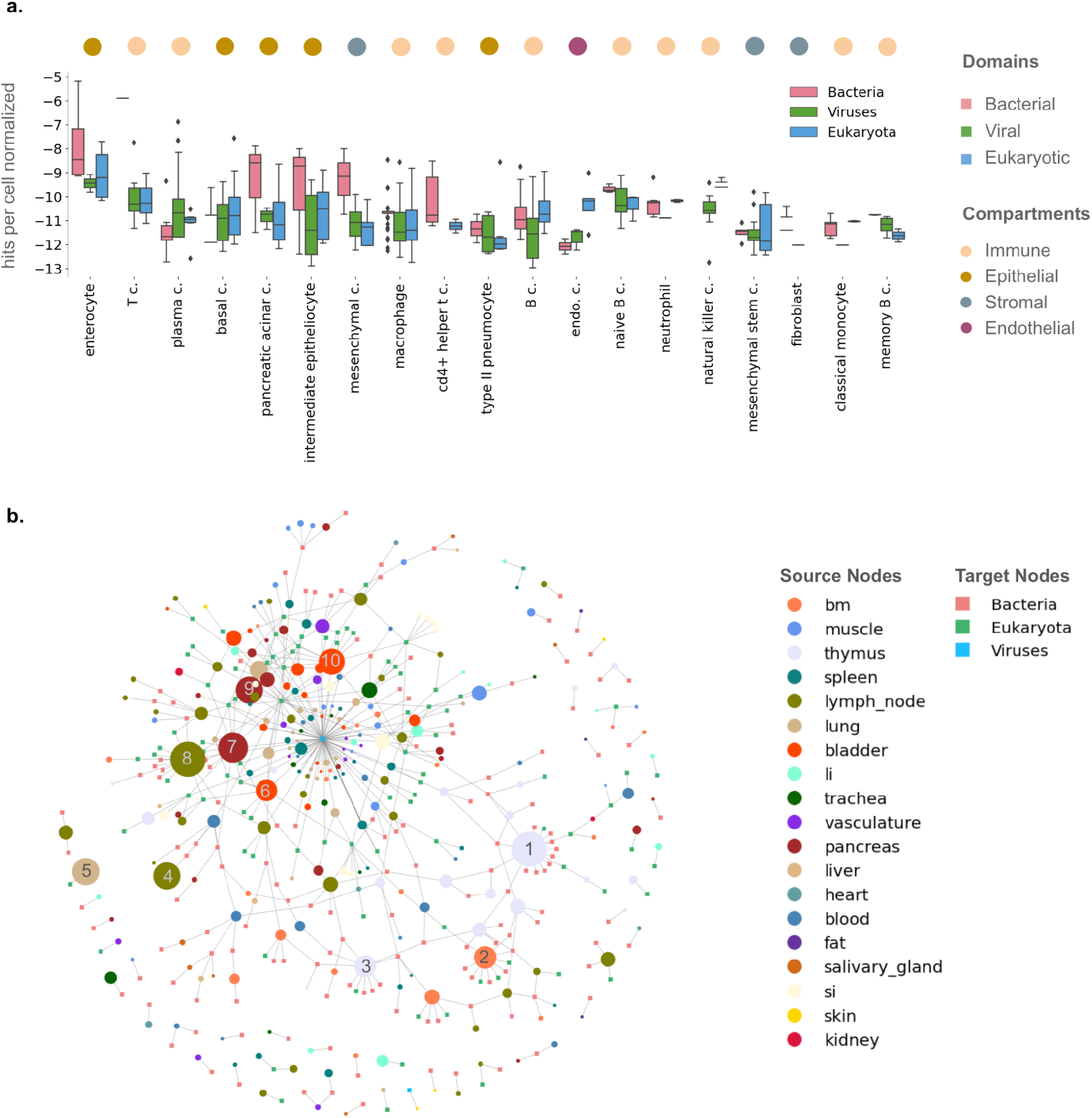
Cellular microbiomes across all donors. a) Distributions of hits per cell against three domains of life, grouped by cell type from TSM F5. Cell types are ordered based on the maximum value found in any of the three distributions. Y-axis corresponds to log_2_(h_c_/g_c_), where h_c_ is the number of hits per cell, normalized by the number of g_c_, number of detected genes per cell. Only cells with detected hits are included in the analysis and only cell types with at least 10 cells. b) Network of human cell types (grouped by tissue of origin) in association with microbes. Cell types are source nodes (circular), and microbial species are the target nodes (square). Source nodes are weighted by S_i_/log_2_C_i_, where S_i_ represents the number of unique species associated with source node *i*, and is normalized by the total number of cells, C, sequenced for source node *i*. The normalization ensures that cell types with greater representation in the dataset do not artificially appear to have more diverse microbial associations. Moreover, edges in this network are weighted by N_j,i_, the counts of sequences mapping to microbial species *j* associated with source node *i.* For visual clarity, only larger source nodes are labeled: 1) Thymus CD4+ T cell, 2) bone marrow CD4+ T cell, 3) thymus fibroblast, 4) lymph node effector memory T cell, 5) lung regulatory T cell, 6) bladder intermediate epitheliocyte, 7) pancreatic acinar cell, 8) lymph node B cell, 9) pancreas fast muscle cell, 10) bladder macrophage. For all other nodes, please see the accompanying interactive plot (SI File 3). bm: bone marrow, li: large intestine, si: small intestine.

### Network of human cell types in association with microbes reveals cell types with greatest microbial diversity

To explore the diversity of cellular microbiomes, we built a bipartite network of cell types stratified by tissue of origin (source nodes) and their associated microbial species (target nodes) for both the TSM F5 and TSM F7 datasets (**Figure 3.b, SI Files 3**). Cell types in each tissue are weighted by the number of unique microbial species, normalized by the number of annotated cells sequenced. Hence, in this network of host-microbe associations, hubs represent cell types with the greatest microbial diversity. Examples of these hubs include Thymus and bone marrow CD4+ T cells, lymph node B cells and CD8+ T cells. As with microbial load measurements, immune cell types are over-represented in their diversity of microbial associations. Cellular microbiome network depictions such as this open a window into host cell and microbe interactions in their native context which would not be captured in bulk tissue sequencing experiments. An example of an important association is between EBV and splenic plasma cells. We report statistics on droplets with hits, and identify outlier cells based on number of hits per cell (see **SI: Single-droplet statistics, outlier cells and outlier cell types,** and **SI Tables 23-26**). Interestingly, the most prevalent species found in outlier cells is EBV, and most outlier cells are from immune cell types.

### Detecting the transcriptional footprints of a persistent viral pathogen and an ancient viral relic in human cells

EBV was the first oncovirus to be discovered and has since been shown to cause Burkitt’s, Hodgkin’s, NK and T cell lymphomas as well as nasopharyngeal and gastric carcinomas, thus accounting for ∼200,000 (or 2% of) cancer deaths annually worldwide^56^. In absence of an approved EBV vaccine, currently most primary infections occur in children and adolescents, such that by adulthood a significant fraction of the population is latently infected^57–59^. Additionally, a recent longitudinal study of 10 million young adults has shown a ∼30 fold increase in the risk of multiple sclerosis after EBV infection, placing EBV as one of the leading potential causes of multiple sclerosis^60^.

During the latent phase of its lifecycle, EBV suppresses the transcription of most of its genome^38^ and replicates infrequently and in synchrony with the host cell while utilizing the host polymerase^61^. In our cohort, we detected EBV sequences in five seropositive donors, and across various tissues including the lung and the bladder. The majority of hits originate from donors 1 and 2 (TSP1 and TSP2) and from lymphoid organs such as the spleen, where many of the susceptible host cell types reside (**Figure 4.a, SI File 4**). Consistent with the known aspects of EBV tropism^62^, we found that the majority of sequences originate from cell types belonging to immune and epithelial compartments.

**Figure 4.**
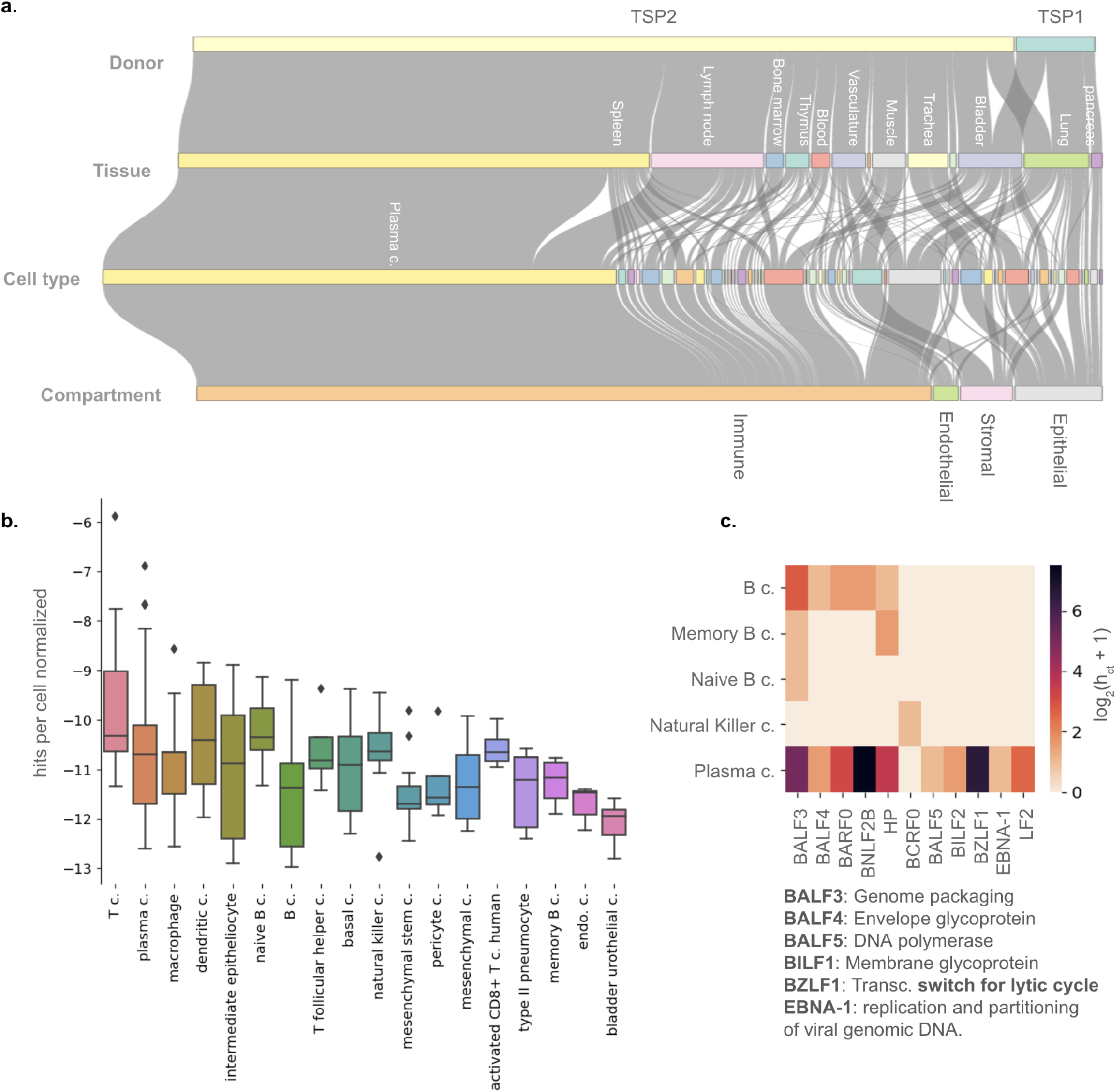
Detection of EBV across tissues and cell types (TSM F7). a) Sankey plot of EBV sequences, demonstrating their donor, tissue, cell type, and compartment of origin. For visual clarity, only hits from annotated cells are included (TSP1 and TSP2 hits remain) and only large groups have been labeled for tissue and cell type rows. Please see the accompanying interactive plot (SI File 4). b) EBV hit per cell distributions grouped by cell type from any tissue that is EBV hit positive. Y axis denotes log_2_(h_c_/g_c_), where h_c_ is the number of hits per cell, normalized by the number of g_c_, number of detected genes per cell. Only cell types with at least 5 cells are shown. c) heatmap of protein-coding sequences identified in TSP2 mapping to EBV, grouped by cell type. The scale bar denotes log_2_(h_ct_ + 1), where h_ct_ represents the number of hits per cell type. The functional role of some of the EBV protein coding sequences are denoted. Note, “HP” corresponds to “hypothetical protein”.

Across distributions of EBV single-cell load in different cell types, plasma cells outrank other cell types (**Figure 4.b**). Applying BLASTx to identify protein coding sequences, we primarily found those expressed during the lytic phase (**Figure 4.c**): membrane glycoprotein (BILF1), DNA synthesis and genome packaging protein (BALF3)^63^, and the viral DNA polymerase (BALF5), which is used to replicate the EBV genome exclusively during the lytic phase^61^. Notably, we observe BZLF1 expression, an important transcriptional switch that initiates the expression of various lytic genes^64^. Although EBV sequences are present across a variety of cell types, only plasma cells, B cells and NK cells are positive for lytic transcripts, consistent with the fact that these cell types are known susceptible host cell types for EBV. Considering that the latent phase of EBV life cycle is characterized by the expression of non-coding RNAs, many of which suppress lytic genes^56^, absence of coding lytic transcripts suggests that other cell types are experiencing latent infections. Alternatively, they may have uptaken transcripts passively by virtue of physical proximity to sites of lytic infections, or actively through their involvement in the clearance of lytically infected cells.

In addition to EBV, we detect the presence of human endogenous retroviruses (HERVs). Introduced numerous times during the evolutionary history of primates, HERV genomes, which are mostly defunct, occupy 8% of the human genome^65^. The HERV K (HML-2) family, representing the latest insertions into the human genome, is the only family of HERVs that is transcriptionally active. Differential expression of HERV K genes has been detected across several cancers and is thought to contribute to tumor progression^66, 67^. Additionally, HERV K expression and even virus-like particles have been detected during human blastocyst formation^68^. Although this virus was tagged as a putative contaminant by our ML decontamination model, we believe this may represent one instance of the model’s false positives. This is because we did not detect HERV K in the ∼200 negative controls obtained for the TSM dataset. We also identified it again separately through another set of experiments described in the next section.

Contrary to the claim that HERV K expression is silenced in all but a few special contexts^69^, we identified HERV K (HML-2) transcripts across five donors, spanning a diverse range of tissues (e.g. skin, fat, spleen and the salivary glands) and cell types (**SI Table 27**). These sequences map to the three canonical retroviral genes in the HERV K genome, namely pol, gag and env (**SI Figure 20**). HERV K expression in human donors beyond tissues and cell types previously investigated, opens the possibility that this enigmatic ancient relic may be more widely active and physiologically relevant than previously appreciated.

### Tissue microbiomes through the lenses of DNA sequencing, fluorescence, and electron microscopy

To further examine our findings through other techniques beyond single-cell transcriptomics, we developed an assay that extracts virus-like and bacterial-like particles (VLPs and BLPs) from human tissue (see **Methods: The Extracted Human Tissue Microbiome dataset)**. We refer to the resulting dataset as the Extracted Human Tissue Microbiome (EHTM). We used DNA sequencing, fluorescence and electron microscopy of VLPs and BLPs extracted from 4 tissue types (pancreas, blood, lymph node, and lung) and 8 donors (**Figure 5.a**). In this assay, we gently dissociate tissues, eliminate cell- or capsid-free nucleic acids, and subsequently size select for VLPs and BLPs. We used negative controls for all experimental steps to eliminate contaminants from bulk DNA sequencing of these extracted particles. Using nucleic acid staining followed by fluorescence microscopy of these particles from the lung and lymph nodes (**SI Figure 21**), we observed high concentrations of VLPs and BLPs (∼10^7^ BLPs and ∼10^8^ VLPs per cm^3^ of tissue). Electron microscopy of VLPs further showed familiar virus morphologies such as a hexagonal capsid (similar to many bacteriophages) and rod-shaped capsids (reminiscent of the tobacco mosaic virus or the rudivirus) in the pancreas (**Figure 5.b**). It also revealed several novel morphologies, for example one with interior nodules and fibrous protrusions residing in a lymph node. Moreover, DNA sequencing of these extracted particles further captured numerous sequences mapping to viruses, bacteria, and even some fungi as some have similar sizes to bacteria (**Figure 5.c**). These results support and are consistent with the single cell transcriptomic data, thus providing independent evidence for several of our conclusions.

**Figure 5.**
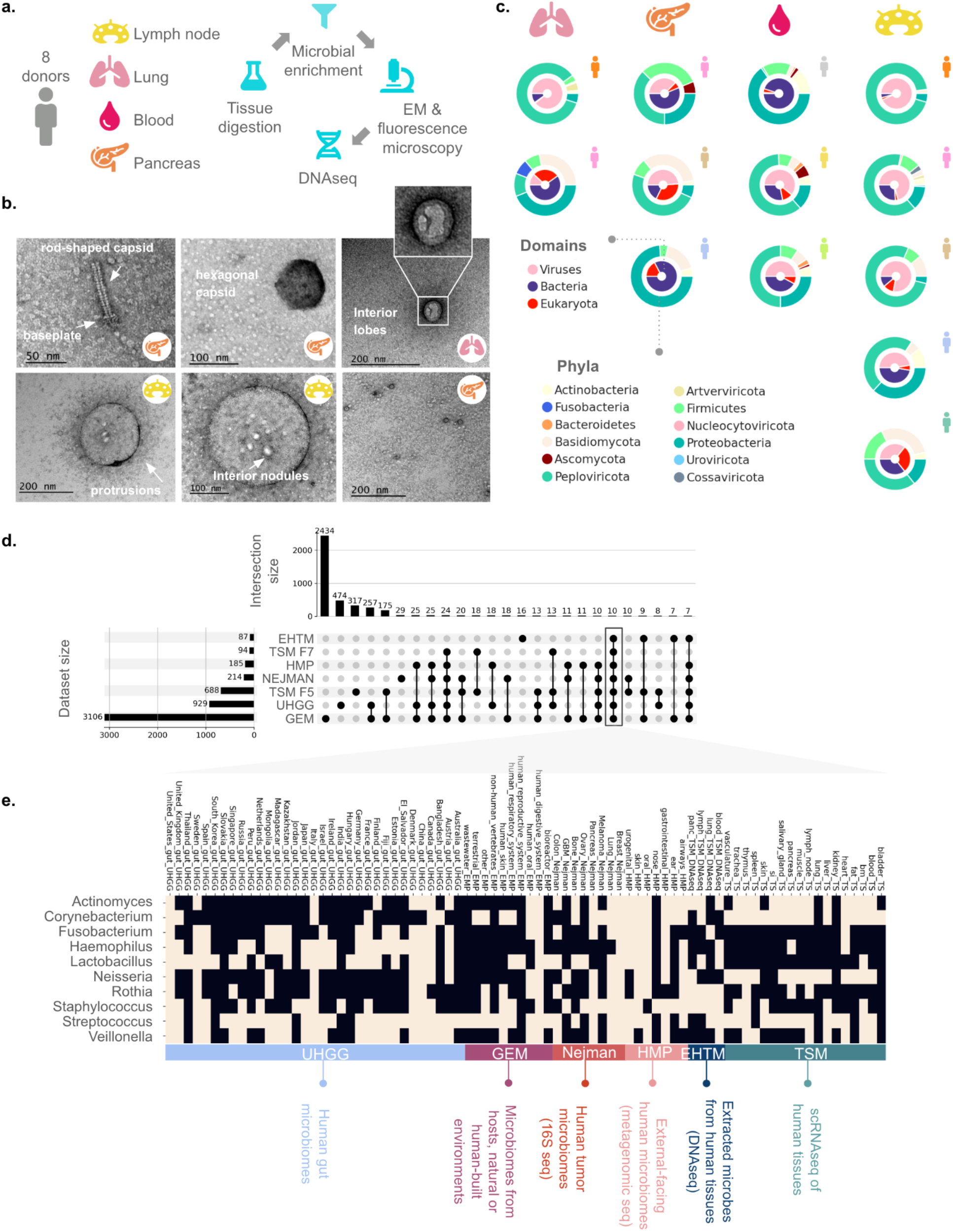
Tissue microbiomes from 8 donors using DNA sequencing, fluorescence and EM imaging from the Extracted Human Tissue Microbiome dataset along with a comparison to several other datasets. **a)** study design and experimental workflow. **b)** EM images of viral fractions retrieved from the pancreas, lung and lymph nodes showing both known and novel virus-like morphologies. **c)** hits per tissue and donor stratified based on taxonomy: domains (inner ring) and phyla. **d**) Exploring the intersection of bacterial genera across various datasets using an UpSet plot: Tabula Sapiens Microbiome (TSM) F5 and F7, Extracted Human Tissue Microbiome (EHTM), Human Microbiome Project (HMP), Nejman *et al.* tumor microbiome, Unified Human Gastrointestinal Genomes (UHGG) collection, and Genomes from Earth’s Microbiomes (GEM). Only sets with a minimum size of 7 are shown for visual clarity. **e**) The genera that are at the intersection of all these datasets are shown. Presence (light boxes) and absence (dark boxes) of each genus is shown. bm: bone marrow, li: large intestine, si: small intestine, panc: pancreas, GBM: glioblastoma multiforme.

Interestingly, EBV and HERV K were both identified in the EHTM cohort. EBV was found in all 8 donors, which is a higher prevalence than obtained via RNA sequencing. Considering that single-cell RNA sequencing preferentially captures transcriptionally active organisms, one would expect to see higher EBV donor positivity rate using DNA sequencing of VLPs. The combination of these two methodologies could perhaps be used to decipher active versus inactive organisms to further make a distinction between residents and passengers of the human tissue microbiome.

Moreover, we identified the genera that overlap between all six datasets: TSM, HMP, Nejman *et al.*, UHGG, GEM, and the EHTM datasets (**Figure 5.d-e**). For example, we found sequences mapping to *Rothia mucilaginosa* in the pancreas and several other organs using single-cell transcriptomics (TSM dataset). This species, a member of the airway microbiome (HMP dataset), appears in pancreatic, lung and breast tumors (Nejman *et al.*). Consistent with its traces in these three datasets, we identify this species through DNA sequencing of the extracted BLPs in the pancreas, lung, blood, and lymph nodes. Given these additional experiments, it is perhaps fair to assume that sequences found in single-cell transcriptomic data are not all cell-or capsid-free. Some perhaps originate from intact organisms that escape cellular barriers to reach internal tissues and tumors.

## Discussion

The transcriptional footprints of microbial life in human organs and cells raise several questions. For example, how could microbes escape cellular barriers established by epithelial and endothelial cells? Are there any long-term residents of human tissues in a commensal relationship with our own cells? And can these microbial signatures be used as diagnostic markers for organ health? One likely route of microbial migration is from the gut to other organs, as leaky gut epithelia together with the gut-liver axis have been proposed as a route of dissemination for various microbial metabolites from the gut into the systemic circulation^70, 71^. The same route is likely to harbor microbial genomes, transcripts, and perhaps whole cells and capsids. In fact, viral migration across epithelial layers has been demonstrated *in vitro*^72^. However, as we have shown there are likely other possible flow paths from other external-facing microbiomes such as the oral cavity and the urogenital tract to various internal organs.

Overall, there is growing evidence for the existence of microbial interactions with human cells at sites in the body that are typically inaccessible to sampling. While microbial sequences have already been shown as diagnostic markers for cancer detection, our understanding of their associations with human cells is largely lacking. It is possible that causal associations exist, and that the list of oncogenic microbes is much larger than previously appreciated. Our finding of significant overlaps between tumor-free tissue microbiomes and tumor microbiomes suggests that tumor microbiomes are at least partially sourced from normal tissue microbiomes. As such, mechanism-level understanding of microbe-host interactions in human tissues could further shed light on the highly variable nature of individual responses to cancer immunotherapies^73, 74^ and provide novel and targeted avenues for microbiome-based immune modulation, as well as lay foundational insights into the interplay between microbes, tumor cells and the host immune cells. Moreover, given that infectious agents account for ∼15% of all cancers^75^, and that many pathogens can manipulate the host cell cycle^76, 77^ and predispose host cells to cancer^78^, it is highly plausible that we have only scratched the surface of causal links between microbes and various pathologies.

With the availability of high-throughput single cell sequencing technologies today, we stand at a unique time in the history of microbiology, where the study of microbe-host interactions offers an exciting new frontier. These interactions can now be studied at the level of individual cells^79–81^ not just *in vitro,* but as we have shown, in the very real context of organs - in the presence of the immune system, neighboring cells, nutrient fluxes and many other important factors that shape microbial life cycles.

## Conclusion

In this study we provide the first atlas of the human tissue microbiome with cell type resolution, for which we have developed new experimental and computational pipelines such as a machine learning-based decontamination model. We discovered sequences mapping to numerous species from three domains of life. We map out the possible flow routes of microbes from external-facing microbiomes to internal tissues and tumors, and show that tumor microbiomes share many common taxa with normal tissue microbiomes. We further constructed a network of associations between human cell types and microbes, shedding light on the types of organisms that cells from different tissues encounter either by way of immune clearance or physical proximity to microbial niches and sites of infections. This type of analysis can reveal the interactions between various cell types and microbial organisms. In cases of intra-cellular infectious events such as EBV, host and pathogen transcriptomic landscapes may reveal novel pathways that underlie cellular susceptibility and resistance, which would have great clinical application.

## Materials and Methods

### The Tabula Sapiens dataset

The Tabula Sapiens raw fastqs containing single-cell RNA sequencing data from Smart-seq2 and 10X droplet-based experimental pipelines were acquired from the Tabula Sapiens consortium^28^ (https://github.com/czbiohub/tabula-sapiens). This included raw fastqs for 12 donors 1,2,4,6,7,8,9,10,11,12,14, and 16 as well as objects which contained cell type annotations for ∼0.5 million cells (donor 16 cell type annotations were not included). All analysis was performed on the 10X data, which represents the vast majority of this dataset. Donor 4 tissues (mammary, myometrium, uterus) were eliminated in the TSM F7 version of the dataset due to not meeting the criteria imposed. Additionally, because we could not acquire negative control samples for tongue tissue, we do not report on this tissue. Thus we have a total of 11 donors and 19 tissues in the TSM F7 dataset.

### The Extracted Human Tissue Microbiome dataset

#### Tissue Digestion and DNA extraction

Tissues were acquired from Donor Network West (delivered on ice) and cut into 1 cm blocks using UV and ethanol sterilized scalpels and petri dishes. Tissues were incubated for 90 minutes with collagenase IV (1 mg/mL, Worthington) at 37°C and further disrupted by pipetting. Disrupted tissue samples were then incubated for 30 minutes at 37°C with 19U/mL DNase I and 1mg/mL RNase A (ThermoFisher) - added to eliminate as much of the cell- or capsid-free DNA and RNA that is released during tissue digestion. The RNAase and DNAase enzymes were deactivated using RNAsecure at 1X and EDTA at 5mM for 10 minutes at 65℃. Each sample was then filtered through a sterile 40µm cell strainer, to eliminate the cellular debris. The filtrate was passed through a 5µm pore size syringe filter to enrich particles in the size range of most microbes and to eliminate host cells. To separate out as many virus-like particles from this mixture of filtrate, we imposed another filter, this time using a 0.8µm pore size syringe filter. The filter was inverted and objects larger than 0.8µm were collected. Although this was a protocol developed in-house, we were able to get key insight from several sources: Qiagen TissueLyser Handbook, Worthington Tissue Dissociation Guide and viral metagenomic studies^82^. DNA extraction was then performed using QIAmp DNA mini kit (Qiagen) according to kit instructions. Moreover, we used multiple negative controls for each step of the experimental workflow, including tissue digestion, DNA extraction, and library preparation.

#### Sequencing and data analysis

We used SIMBA on this dataset, excluding steps used for UMI extraction. We used the negative control samples to create a blacklist of contaminants, which were subsequently removed. Despite significant methodological differences between the EHTM dataset, and the TSM single-cell transcriptomic dataset, we found members from the Actinomyces, Neisseria, Rothia, Corynebacterium, Nesterenkonia, Streptococcus, Veillonella genera shared between the two datasets. However, it is important to note that the level of overlap between these two datasets and any others depends on parameters such as sampling depth and method of detection.

#### TEM

We plasma treated EM grids (300 mesh Carbon/Formvar coated Copper grids) using Leica EM ACE600 to allow each sample to bind to the grid. We applied 5µL of 1:1000 diluted sample onto each EM grid and allowed the sample to bind for 3 minutes. The grid was then exposed to 1% uranyl acetate for one minute, and subsequently dried using a filter paper. Samples were imaged using JEOL JEM-1400.

#### Fluorescence imaging

Following tissue digestion and microbial extraction, filterates were stained with SYBR™ Gold Nucleic Acid Gel Stain (ThermoFisher) at 10X for 10 minutes at room temperature and imaged using Leica CTR6000. Bright dots were counted manually across different sections across a hemocytometer grid and average counts were projected to estimate VLP and BLP concentrations per cubic centimeter of tissue.

## Acknowledgements

We thank Rob Phillips for insightful comments on the manuscript. We also thank Leila Sidow, Robert Jones, Roja Kolluru, Ed Marti, Elizabeth Jerrison, Mark Kowarsky, Alina Isakova, Stephanie Tzouanas Schmidt, Yuan Xue, Mira Moufarrej, Michael Swift, Ivana Cvijovic and all the other members of the Quake lab, along with the Tabula Sapiens and the Boundaries of Life consortia for their help and suggestions towards experimental and computational pipelines. Additionally, we thank the following Tabula Sapiens consortium members for their help towards tissue digestion controls: Mallory Ann Laboulange, William Kong, Siva Vijayakumar, Hua Dong, Rebeca Arroyo, Hayley Raquer, Michael Swift, Zhen Qi, Will Goodyer, Gabriel Loeb, Simon Bucher, Helen Zue Zhang, Soochi Kim, Yan Hang, Juan Irwin, Emily Trimm and Irene Fan. We are grateful to Donor Network West personnel including Ahmad Salehi and Ravi Ponnusamy, as well as the Cell Sciences Imaging Facility at Stanford University. We are especially indebted to donors and their families for their incredible contribution to science. This work was supported by the Chan Zuckerberg Biohub, the Gordon and Betty Moore Foundation and the John Templeton Foundation.

## Funding

This work was supported by the Chan Zuckerberg Biohub, the Gordon and Betty Moore Foundation and the John Templeton Foundation.

## Author contributions

Conceptualization: GM, SRQ

Methodology: GM, SC, SRQ

Investigation: GM, SRQ

Data Curation: GM

Formal Analysis: GM

Visualization: GM

Software: GM

Funding acquisition: GM, Tabula Sapiens Consortium, SRQ

Resources: GM, Tabula Sapiens Consortium, SRQ

Project administration: GM, Tabula Sapiens Consortium, SRQ

Writing – original draft: GM

Writing – review & editing: GM, SC, SRQ

## Competing interests

Authors declare that they have no competing interests.

## Data and materials availability

All data and code is available through our GitHub repository: https://github.com/gitamahm/human_tissue_microbiome_atlas. This repository includes SIMBA, which is written in Snakemake^32^, Post-SIMBA data analysis and visualization organized into separate Jupyter Notebooks (written in python v3.6.10). The Jupyter notebooks are listed below.

- p01_databaseCreationSC_microbeDB_fungalDB.ipynb
- p02_creating_truth_datasets_for_testing_simba.ipynb
- p03_testing_truth_datasets.ipynb
- p04_experimental_negative_controls.ipynb
- p05_post_SIMBA_processing_alldonors_and_EHTM_gh.ipynb
- p06_cell_ranger_counts_gh.ipynb
- p07_machine_learning_modeling_contamination_and_other_filters.ipynb
- p08_habitat_assignment_using_GEM_HMP_UHGG.ipynb
- p09_davinci_language_model_habitat_of_origin_gh.ipynb
- p10_impact_of_filters.ipynb
- p11_single_cell_microbiome_stats.ipynb
- p12_tabula_sapiens_multibiome_analysis.ipynb
- p13_ts_intersection_with_external_datasets.ipynb

## SI Text

### SIMBA, a computational pipeline for the identification of bacteria, viruses and fungi in single cells

With the rapidly rising number of single-cell transcriptomic datasets, there is a growing interest in detecting microbial transcripts within these datasets. Although there are many existing tools for microbial profiling in standard metagenomic datasets, tools that take single-cell transcriptomic data as input are currently more limited^83^. There are however a growing number of pipelines being developed for this purpose, including those developed by Robinson et al. (CSI-Microbes)^84^, Ghaddar et al. (SAHMI)^23^, and Llorens-Rico et al.^85^. CSI-Microbes supports three different existing tools for detection of microbial sequences, namely GATK PathSeq^86^, CAMMiQ^87^ and SRPRISM^88^. GATK PathSeq relies on BWA-MEM alignments, replacing its original use of BLAST^89^ in favor of faster alignment speed. Llorens-Rico’s pipeline uses Kallisto pseudo-alignments against a reference database of ∼12K microbial reference genomes. SAHMI implements kmer-based aligners against a microbial and human reference dataset and implements additional filters to detect false positives.

In our pipeline, called **Si**ngle-cell **M**ultibiome **B**LAST **A**nalysis (SIMBA), we first use UMI-tools^33^ to filter and extract cell barcodes and UMIs from raw fastqs using the *extract* and *whitelist* commands. We used STAR^34^ (v2.7) to align resulting sequences against a reference library containing the human genome (GRCh38.p13) and ERCCs. We then took unmapped sequences for further processing. We wrote *poly.py* to eliminate low-complexity sequences with high fractions of consecutive bases, often representing primer dimers and other sequences with long stretches of polyA. We used BLASTn^35^ (v2.10) to align against several custom databases followed by the nt database, which is one of largest publicly available databases, consisting of ∼70 Million sequences (11/02/2020) pooled from GenBank, EMBL, DDBJ, PDB, and RefSeq databases. Taxonomic lineages for each top hit against the nt database were added using TaxonKit^90^. Top hits were filtered based on their taxonomic assignment to include only viral, bacterial and fungal species.

The premise behind creating a pipeline that relies on BLAST given its high computational cost, is based on our early observations that some of the faster tools for microbial profiling can result in an unacceptably high false positive rate. As an example of a kmer-based profiling method, which are among the most popular and widely used methods for microbial profiling, we explore instances of false positives from Kraken2 against the nt database using several truth datasets. We demonstrate why in the context of our dataset, where the vast majority of sequences are human-derived, such tools lack the necessary precision (see **SI: Benchmarking Kraken2 precision and recall**). Robinson et al. similarly employ GATK PathSeq rather than faster kmer-based profiling methods^84, 87^. We further show that while Kraken2’s recall is high when the input sequences are sampled from the Bacterial Refseq dataset, it is highly variable and can be much lower when the input sequences are sampled from other datasets. This is one example illustrating why alignments using BLAST against large, comprehensive reference databases such as nt are still widely considered the gold standard in precision against which new tools are often benchmarked^91–94^. Because the study of microbiomes in low-biomass settings presents a high burden of proof with many potential sources of false positives, in our approach we favor precision over computational time and recall. In these settings, we and others argue that the problem of bioinformatic false positives is critical^89, 92, 95, 96^ and on par with the problem of experimental false positives, or contamination.

However, aligning large datasets against large databases using BLASTn is computationally infeasible even with access to large compute clusters. To lower this computational cost, we created a lean human-microbiome centric database for bacteria and fungi which we call microbDB (see **SI: Creating custom databases to limit the computational burden of BLASTn against the nt database**). We use microbeDB as a first-pass filter to select sequences with the highest likelihood of being bacterial or fungal in origin. Additionally, through subsampling, we created leaner versions of the publicly available bacterial and fungal RefSeq databases^97^ to use as second-pass filters. Viral sequences were aligned against the Viral RefSeq database^97^ (release 98) which included 9,000+ viral genomes. Surviving sequences that produced significant alignments or “hits” (E-value < 10-5) against these intermediate databases were then aligned against the nt database. This particular E-value threshold is commonly used^98–101^. For each sequence the top significant hit (lowest E-value) was selected for downstream analysis. We wrote *dedup.py* to create input fasta files with sequences that produced a top hit in this step, and use the resulting file to BLASTn against the nt database. By taking this type of screening strategy, we lower the computational burden of alignments by ∼100 fold. Without the initial screening, we would require ∼3 × 10^8^ CPU hours, equivalent to ∼$9 million using Amazon’s compute instances (estimate made in 2020).

The resulting top hits that emerged from the viral, bacterial and fungal workflows against nt were combined into one dataset consisting of ∼19 million sequences for further processing and quality control. Despite first filtering human reads using the STAR alignment step, out of ∼19 million sequences that generated significant hits against nt, only 12% mapped to the viral, bacterial, and fungal biomes of interest, while half of them mapped back to the human genome. This indicates the importance of implementing multiple checkpoints to eliminate non-microbial reads, and cautions against relying solely on smaller databases without validation against a comprehensive database such as nt. This is especially relevant to human tissue samples where the ratio of non-microbial to microbial sequences is high^86, 89, 95^.

We applied our pipeline not only to the raw data underlying Tabula Sapiens annotated single cell dataset but all the raw data, including those from empty droplets, which we refer to as the Tabula Sapiens Microbiome F00 (TSM F00) dataset. The rationale for including unannotated or empty droplets is that we suspect that microbial reads from extracellular environments could appear in those droplets, and more generally, different species may have different patterns of association with these two categories of droplets. Later we incorporate this information for each species as a parameter in our machine learning models for decontamination. Using TSM F00 as input, we analyzed reads from 150+ Million droplets, ∼1.3 Million of which had more than 200 human genes and that we treat as potentially cell-containing. By matching the droplet barcodes (along with sample name) to those found in the Tabula Sapiens objects, we were able to acquire the cell type annotations for a portion of the cell-containing droplets. For all other droplets, we provide tissue and donor information. Please see **Data and Materials Availability** for the Jupyter notebook where details of cell-containing and empty droplet counts using cell ranger can be found.

The output of SIMBA once filtered to contain only bacterial, fungal or viral hits was 2.4 Million significant hits. There were ∼724K significant unique hits (counted based on Unique Molecular Identifiers or UMIs) arising from ∼0.5 Million droplets - 79K cell-containing droplets (∼6% of TSM F00 cell-containing droplets) and ∼380K empty droplets (∼0.3% of TSM F00 empty droplets). This dataset is what we will refer to as the TSM F0 dataset.

### Creating custom databases to limit the computational burden of BLASTn against the nt database

RefSeq databases^97^ for bacterial and fungal genomes are available through NCBI, however they are typically large databases that would create significant computational bottlenecks. We created several custom databases to serve as lean human-microbiome centric databases for the initial iteration of BLASTn search for bacterial and fungal sequences. This was not done for viruses as they have much smaller genomes relative to bacteria and fungi and thus the viral RefSeq could be used directly.

MicrobeDB was constructed by combining two other custom databases, bacteriaDB and mycoDB. BacteriaDB was designed by taking the intersection of the HumGut database^102^, a comprehensive database of ∼30,000 prokaryotic genomes mined from ∼5,700 healthy human metagenomes, and the bacterial RefSeq database (downloaded on 01/2021). The HumGut dataset was obtained from https://arken.nmbu.no/~larssn/humgut/. We therefore identified the unique set of species in HumGut and eliminated entries with unidentified species-level taxonomy. Similarly, we identified the unique set of bacterial RefSeq entries, and took the intersection of these two sets to arrive at several hundred bacterial species. We used a similar strategy to bias the initial search for fungi towards species previously identified in human mycobiome studies. We took five representative species from each genus found in both the fungal RefSeq (downloaded on 01/2021) and a recent survey of the healthy human mycobiome^103^ to create mycoDB. While microbeDB was used for the first search iteration, larger databases were used for the second iteration. These larger databases, slimBacterial_RefSeq and slimFungal_RefSeq are the reduced versions of the bacterial and fungal RefSeq databases which we subsampled to include representative species and strains. Please see **Data and Materials Availability** for the Jupyter notebook where details of how intermediate databases were constructed is described.

### Benchmarking SIMBA’s precision and recall

We evaluated SIMBA’s precision and recall against several truth datasets. For all truth datasets, we randomly selected sequences that were 100 bases long to match the length of sequences in the TS dataset. We will refer to each sequence from truth datasets as “input seq” and its corresponding output hit from SIMBA as “output seq”. If an input seq goes undetected by way of not producing any corresponding output seq, we count that as a false negative. We use taxonomic IDs on input and output seqs to measure precision and recall at each level of taxonomic classification. For example, if an output seq does not match an input seq in taxonomic ID or species name, we consider that as a false positive at the species level of classification.

First, we created a dataset of 2400 sequences derived from human chromosomes (GCF_000001405.39_GRCh38 assembly) including X and Y, with 100 randomly selected sequences from each chromosome. We then employed SIMBA’s bacterial and fungal branch of the pipeline without using STAR to eliminate human-derived sequences as we knew most of these input seqs would already be eliminated at this step. Even without this step, the pipeline did not produce any false positives. Only one input seq produced a hit when aligned against microbeDB and that sequence was eliminated in the subsequent alignment step against the next larger intermediate database, thus no false positives were detected.

We also used this truth dataset against the viral branch of the pipeline, again forgoing the STAR alignment step. Out of 2400 sequences, 65 produced hits against the viral Refseq database, and were subsequently aligned against the nt database. At this stage, all these sequences were correctly identified as human in origin, including one human retroviral sequence. Thus, SIMBA produced no false positives when encountering human-derived sequences, which is an important aspect of this pipeline considering the high ratio of human to non-human sequences in the TS dataset and in human tissues more broadly.

Next we tested SIMBA’s performance when encountering viral sequences. To create a truth dataset, we randomly selected 100 viral genomes and selected 10 random sequences from each genome using the viral RefSeq database. This resulted in sequences from 62 genera of viruses, thus representing a diverse viral truth dataset. We found that all input seqs produced an output seq. Interestingly we noticed that some mismatches that were encountered between the taxonomic IDs of input and output seqs were the result of finding a prophage output seq when the input seq originated from a phage. Because the prophage is embedded in a bacterial genome, the resulting output seq had the taxonomic ID of the host bacterium. We excluded these special cases from calculations of precision and recall. At the species level, we see the following precision and recall, respectively: 0.8, 1 **(SI Table 1)**. False positives detected at the species level correspond to sequences from highly similar viral species that typically share the same host **(SI Table 2)**. This is expected as short sequences (e.g. ∼100 bp) can be shared identically or with very high similarity among related species. Indeed, we found that all false positives had 100% sequence similarity across the entire length of the input seq **(SI Table 3)**. Hence, genus level classification provides a more meaningful depiction of precision as it is less prone to this type of ambiguity in the mapping of short reads. Indeed, the precision jumps to almost 1 when examined at the genus level.

Moreover, the high recall associated with our viral search is expected given that we did not use intermediate, subsampled databases to lower the computational cost for the viral search since it does not pose a computational bottleneck. The lower computational cost of searching for viruses is a result of viruses having 1) some of the smallest genomes across all domains of life, and 2) fewer representatives than bacteria in many databases, including the nt database.

Next, we characterized false positives as well as other parameters in the face of bacterial input seqs. Given that environmental bacteria can be a source of contamination in kits and reagents, we aimed to bias SIMBA to finding those bacteria that are more likely to reside in the human microbiome to 1) mask potential contaminants and 2) limit the initial search space to reduce the pipeline’s computational time. To introduce this bias, we created microbeDB which serves as a first pass filter on the input data. This database is composed of bacteria and fungi commonly found in the human gut microbiome, based on the assumption that the gut likely contributes the most to human tissues due to its greater microbial biomass compared to any other site in the human body. Therefore, to evaluate the pipeline’s performance, our first truth dataset contained bacteria cultured from the human gut microbiome.

We obtained taxonomic IDs from the Human Gastrointestinal Bacteria Culture Collection (HBC) - to select representative genomes from 50 random bacterial species cultured from the human gastrointestinal tract, for which we downloaded a representative genome from NCBI. We obtained the Human Gastrointestinal Bacteria Culture Collection (HBC) dataset from https://www.ebi.ac.uk/ena/browser/view/PRJEB23845?show=reads (filereport_read_run_PRJEB23845_tsv.txt). We then selected 100 random contigs, for a total of 5000 sequences. Using this truth dataset, we report the corresponding precision and recall **(SI Table 4)**. At the species level, we arrive at the following precision and recall, respectively: 0.69, 0.22. As with the viral truth dataset, when we analyzed the false positives, we found that the vast majority represent sequences from highly related bacterial species **(SI Table 5)** with nearly identical or 100% identical output seqs to input seqs. We again see a jump in precision to 0.93 going from the species level of classification to the genus level. We have a recall of 0.28 at the genus level, and thus we likely underestimate the true number of bacterial species using this pipeline. However, we wanted to limit instances of false positives even at the expense of false negatives. In future iterations of this pipeline, increasing the size and diversity of microbeDB would likely enhance the pipeline’s recall.

To test how well SIMBA identifies members of another microbiome, we created a truth dataset based on the expanded Human Oral Microbiome Database (HOMD), consisting of 50 randomly selected genomes and 100 randomly selected segments from each. The expanded Human Oral Microbiome Dataset (HOMD) was obtained from https://www.homd.org/download~genome, specifically from HOMD_taxon_table2022-09-15_1663245201.txt and HOMD_genome_table2022-09-15_1663245202.txt files. Unexpectedly we found similar values for precision and recall across different classification categories **(SI Table 6)**. This stands in contrast to the metrics we obtained when we used a truth dataset that was not focused on bacteria from the human microbiome.

We randomly selected 50 bacterial genomes from the bacterial Refseq database, and randomly selected 100 sequences from each genome for a total of 5000 sequences. Comparing the performance results of this truth dataset at the genus level to the gut or oral bacteria truth datasets, we see that while the precision remains high, recall has dropped down to only 0.05 **(SI Table 7)**. Thus, the pipeline preferentially recalls bacteria from the human microbiome compared to any randomly selected bacterial sequence. It can mask potential contaminant bacterial sequences. Even those input seqs that produce output seqs have a high chance of being correctly identified due to the pipeline’s high precision afforded by the last BLASTn step against the nt database.

We created a fungal truth dataset by taking ∼5000 sequences randomly sampled from a study of the gut mycobiome^104^. At the genus level, we found similar values of recall and precision to those we found for the bacterial truth datasets created from HOMD and HBC (**SI Table 8**). Please see **Data and Materials Availability** for the Jupyter notebook where our integration of datasets for benchmarking SIMBA is described in detail.

### Benchmarking Kraken2 precision and recall

We tested the precision and recall of a popular tool in the field of microbial metagenomics, Kraken2, both against the microbial truth datasets that we created to test SIMBA as well as a sample of the Tabula Sapiens dataset. We chose Kraken2^105^ as it is one of the fastest and most accurate metagenomic taxonomic profiling tools available, and in general Kmer-based approaches to fast taxonomic profiling represents a good fraction of the tools that are available^91^.

This was done primarily to see whether Kraken2’s speed performance would justify its use for our dataset. We assembled the nt database to use Kraken2 against. We found that, in agreement with a previous benchmarking study^91^ of Kraken2 and related tools, Kraken2 has very high precision and recall against the bacterial RefSeq truth dataset **(SI Table 9)**. However, its recall is quite lower for the other bacterial truth two datasets, starting from 0.53 at the genus level against the truth dataset created from HBC **(SI Table 9)**.

Upon removing human-derived sequences using STAR for a bladder sample from the Tabula Sapiens dataset, we used Kraken2 against the nt dataset. Kraken2 created outputs marked classified and unclassified. More than 60% of reads remained unclassified at the species level, and nearly half of the reads still remained unclassified at the domain level **(SI Figure 2a)**. Using the provided taxonomic IDs, we obtained taxonomic lineage information for reads that were classified.

Among the classified reads, ∼285K were identified as eukaryotic, ∼2.7M as bacterial and ∼1.1M as viral **(SI Figure 2b)**. To test the precision and recall of these classification calls, we subsampled each domain (except Archaea) and randomly selected 100 sequences from those reads that had acquired full classification up to the species level. We then used BLASTn against the nt database to measure the extent of the agreement between Kraken2 and BLASTn taxonomic classifications. Surprisingly, we found very little agreement and a high rate of false positives associated with Kraken2 classifications. For bacteria and viruses, the mean precisions at the genus level are 0.06 and 0.04, respectively **(SI Table 10)**.

Taking a closer look at instances of false positives for reads that Kraken2 classifies as various species of bacteria, we see that many are classified as human (and other primate species) **(SI Table 11)**. Human reads that are not sufficiently eliminated by STAR present a major challenge for Kraken2. This issue of high false positive rate when analyzing clinical samples has been discussed in the past^89, 91^. Given a large enough number of human reads, as is the case with Tabula Sapiens datasets, many sequences will be misclassified as microbial or viral using this tool. On the contrary, because our pipeline relies on BLASTn against nt, it has significantly better precision than kmer-based approaches. Hence, we could not justify the faster speeds associated with some of the existing metagenomic classification tools over their lack of extreme precision for this specific application.

### Measures against contamination during tissue recovery and transport

As detailed by the Tabula Sapiens consortirum^28^, the tissues were procured at various hospitals through collaboration with Donor Network West (DNW, CA) - a not-for-profit organization that is a federally mandated organ procurement organization. Tissues were procured consistently across all donors across different operating rooms (OR). Contaminants from air, hair, skin, and instruments were carefully mitigated using strict aseptic techniques as outlined by the American Association of Tissue Banks (AATB) some of which is briefly discussed in the following paragraphs.

In each OR, multiple sterile fields are created. A sterile field is created for each time a pair of sterile gloves or gowns is opened. An additional sterile field is set using sterile surgical drapes around a surgical site. Several sterile fields are created on tables where sterile surgical instruments are placed. Another sterile field is created on a table where storage and shipping supplies are placed.

To mitigate the risk of contamination from air, the OR rooms used to recover organs contain elaborate air filtration systems, and OR personnel are required to wear sterile PPE such as a face mask at all times. They also wear hair bonnets prior to putting on surgical scrubs and maintain them at all times. The OR personnel scrub their hands and arms before a procedure; using sterile surgical attire as barriers between their skin and recovery environment, they then prepare the donor for tissue recovery.

The donor is prepped for tissue recovery using rigorous aseptic protocols as outlined by AATB (cite). First, clothing is removed, and hair is removed from any recovery areas using surgical scrub soap to prevent contamination from the donor’s skin. These areas are then wiped with 70% Isopropyl Alcohol to disinfect the skin while also removing any loose hair or debris not removed by the surgical soap. FDA-tested and approved surgical soaps are used. Donor draping is performed using aseptic standards of practice to isolate, form a barrier and create a sterile field. Recovered organs are protected by a triple sterile barrier and stored in enough cold, sterile preservation solution (UW solution) to sufficiently cover the organ or vessel. To capture any contamination possibly introduced through the UW solution, we performed sequencing of this solution.

It is important to note that given our study relies on post-mortem samples, thus there is a possibility of tissue barrier breakdown that would impact microbiomes. To partly address this concern, we set up a private courier service to keep the time between organ procurement and initial tissue preparation to less than one hour. This is a relatively short period considering that heart and lung procured for transplantation are viable for 4-6 hours, liver for 8-12 hours, and kidneys for 24-36 hours. As tissues were recovered in the same manner as those used for transplantation, and were rapidly prepared for single cell sequencing, the possibility of gross tissue breakdown is low. Additionally, histological staining of tissues showed viability along with certain markers used to identify rapidly dividing cells (e.g. transient amplifying cells of the intestinal epithelium)^28^. Using single-cell transcriptomes, we could differentiate between hundreds of cell types, identify differentiation trajectories, and tease apart different phases of the cell cycle^28^. Together these findings point to cellular and tissue viability at large.

### Identification of contaminants based on negative controls and external datasets (Filters 1-3)

The introduction of contaminants was mitigated during tissue procurement in the OR setting by creating sterile fields and various other strict measures as these tissues were collected in the same manner as those recovered for organ transplantation. These tissues were placed in UW solution and then within sterile bags on ice and quickly shipped to different tissue expert labs to preserve tissue and cellular viability. Within different labs, organs were further dissected into smaller tissue pieces and digested to create single-cell suspensions in preparation for single-cell sequencing. A detailed description of tissue procurement and transport is provided in **SI: Measures against contamination during tissue recovery and transport.**

There are multiple potential sources of contaminants, including contamination introduced during tissue procurement, tissue transport, tissue digestion and sequencing. However, we suspect that different sources of contamination do not have equal contribution to the total list of contaminants in each sample. We posit that for dense tissue types, contaminants from instruments, reagents, or air are exposed primarily to the surface of each tissue sample, and thus surface area to volume ratio, SA:V, is an important physical constraint on each contamination source. We will illustrate the importance of SA:V using simple back-of-the envelope estimates, assuming samples are cubes for simplicity. For a cube, SA:V is (6 × s^2)/(s^3) or 6/s, where s is the length of a side. Let us assume the organ recovered in the OR is 10 cm on each side, and that it is further dissected into 1 cm cubes in a lab and digested into 10 um cubes (or cells). In this process, the SA/V ratio starts at 0.6 cm^-1 in the OR setting. Then in the lab setting, it goes up to 6 cm^-1 at the dissection step and then to 6000 cm^-1 during the tissue digestion and single-cell suspension step. Because of the increase in SA:V, we estimate that contaminants are orders of magnitude more likely to come into contact at the tissue digestion and sequencing steps than they are at tissue procurement, transport or dissection steps.

All labs performed tissue digestion and sequencing using aseptic techniques, however because these samples were largely intended for studying the human transcriptome, no blank instrument or reagent controls were generated at the time of the original experiments. In absence of these controls, we partnered with the labs that had processed these tissues and obtained reagents and consumables from the following tissue experts for the Tabula Sapiens project: bladder, blood, bone marrow, fat, heart, intestines, kidney, liver, lung, lymph node, muscle, pancreas, salivary gland, skin, spleen, thymus, trachea, uterus, and vasculature. The reagents and consumables are listed by tissue in **SI Table 12**. The tissue dissociation protocols were performed exactly as detailed by the Tabula Sapiens consortium^28^ in small scale (∼250 µl final eluate) but in the absence of tissue. These protocols included incubation steps, centrifugation, sample filtration using cell strainers, enrichment using MACS columns, among other steps. Even when the same reagent was used across different labs, we obtained each lab’s specific batch of that reagent to account for lab-specific contaminants.

Equal volumes of supernatants obtained at the end of tissue dissociation protocols were pooled and mRNA seq libraries were generated using the Chromium Next GEM Single Cell 3’ v3.1 chemistry (10X Genomics, Pleasanton, CA). Each pool was loaded onto a Chromium G Chip channel for a total of 5 channels. Libraries were prepared using manufacturers recommendations. All PCR reactions were performed using the Biorad C1000 Touch Thermal cycler with 96-Deep Well Reaction Module. As per 10X Genomics protocol recommendation, amplified cDNA and libraries were evaluated on a 2100 Bioanalyzer system (Agilent Technologies, Santa Clara, CA). Libraries were sequenced on the NextSeq 500 (Illumina, San Diego, CA) to obtain a total of 86 million reads.

With ∼200 negative control samples, we arrived at a list of contaminant species. When we excluded all species found in negative control samples, the number of unique hits was reduced to ∼63K, or roughly 9% of the TSM F0 unique hits. Interestingly, the number of species dropped from ∼8200 to only ∼7100 and the number of genera went down from ∼2500 to ∼2400. Filter 1 reduced the numbers of cell-containing and empty droplets to ∼13K (∼17% of TSM F0 cell-containing droplets) and ∼44K (∼12% of TSM F0 empty droplets), respectively.

We then imposed a second filter based on genera commonly found in negative controls from previous studies^36^, which we will refer to as the Salter genera. The removal of these genera reduced the number of unique hits, species and genera to ∼52K, ∼6200, and ∼2300, respectively. It reduced the numbers of cell-containing and empty droplets to ∼12K and ∼35K.

Next we measured 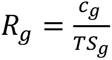 where *TS_g_* represents the number of species from genus *g* found in the TSM dataset, and represents the number of species from that genus in the negative control dataset. This simple parameter can provide insight into how confidently species from a given genus can be flagged as true signal or contaminant. For example, if a considerable fraction of species in a genus found in the TSM dataset are also identified in the control dataset (high *R_g_*), our confidence in other species from that genus would be lower than in the reverse scenario. This is because as we showed through benchmarking of both BLASTn and Kraken2 against several truth datasets, species-level taxonomic assignments based on short reads (∼100 bases) are less precise than genus-level taxonomic assignments. It is also possible to experimentally undersample contaminants and miss certain species. However, blanket removal of genera may also result in a significant loss of true signal. Moreover, comparing the distributions of *R_g_* for Salter genera versus all others in the TS dataset, we observe significantly larger values (one-sided Mann-Whitney U test, P < 10^-5^), providing support that this parameter can be used to help distinguish between genera that are known contaminants versus true signal. Thus, rather than defining an arbitrary cutoff for *R_g_*, we incorporated it in several machine learning models of contamination (described in the next section) and allowed these models to learn this parameter along with others. Indeed, the inclusion of *R_g_* boosted the area under the precision recall curve (AUPR) as well as the area under the receiver operator curve (AUROC) for the machine learning models discussed in the next section.

We imposed a third filter to eliminate putative fungal species based on a recent study of fungi in human tissue samples^20^. This filter reduced the number of species to ∼6100. Other numbers remained nearly the same (rounded numbers do not reflect a change, see exact numbers in **SI Table 13**).

### Machine learning models of contamination (Filter 4)

Despite having collected many negative control samples representing reagents and instruments used by the Tabula Sapiens consortium, we built 6 different types of machine learning (ML) models to learn the shared features of the known and putative contaminants in the TSM dataset to identify potentially missed contaminants. We used the best performing model to identify putative contaminants in the part of the dataset that is withheld from training and testing. As input to each model was a table where the rows were species and the features were: 1) number of tissues and 2) number of donors positive for that species, 3) abundance of that species, 4) median percent identity and 5) median length of alignments corresponding to that species, 6) *R_g_* (described in the previous section), 7) domain-level taxonomic assignment, and 8) median number of genes per droplet in which the species was identified. The last parameter is included to incorporate information about whether the sequence corresponding to a given species is coming from an empty droplet or from a cell-containing droplet. The premise behind including the last parameter is that we suspected contaminants and true signals may have different patterns of association with those two classes of droplets. Counts of empty droplets were performed using Cell Ranger (v6.0.1) in addition to a custom script for downstream analysis (see **Data and Materials availability**).

If a species was flagged as a known or putative contaminant by any of the first three filters (see previous section), it was considered as a contaminant (labeled 1). This dataset was split into parts A (25% of total) and B (75%). Part A dataset was split into training (80%) and test (20%) subsets (**SI Figure 1.c**). We built k-nearest neighbor, multi-layer perceptron, gradient boosting, random forest, logistic regression, and support vector RBF (radial basis function) models. Because the total dataset (part A + B) contains a small fraction of contaminating species, it represents an imbalanced dataset where the detection of the minority positive class, or contaminant species, is of interest. Thus, to balance the training dataset (and only the training dataset), we employed SMOTE (Synthetic Minority Oversampling Technique)^106^ to generate synthetic minority class instances from existing ones. Next, we tested the models on the unseen test dataset and repeated this process 10 times using different train/test splits to generate 95% confidence intervals for test performance. We report the performance of the model on test data. The test AUPR and AUROC performance for the models are provided in **SI Figure 1.d**, ordered by the median value. The 95% confidence intervals for AUPR and AUROC values are provided in **SI Table 14** and **SI Table 15**. Further details on model creation and hyperparameters is provided in p07_machine_learning_modeling_contamination_and_other_filters.ipynb (see **Data and Materials availability**).

Although we provide AUROC test scores, we rely primarily on AUPR as the performance metric of interest since AUROC may be artificially high when the dataset consists of a minority positive class^107^, though in our case the dataset is only moderately imbalanced. The null AUPR, which corresponds to the prevalence of the positive class, is ∼0.3 and denoted by the dashed line (**SI Figure 1**). The gradient boosting classifier (GBC) had the highest median AUPR test scores (95% confidence interval: 0.75-0.78) and highest median AUROC (95% confidence interval: 0.82-.84).

We then fit the GBC model trained and tested in Part A to Part B. We eliminate the known and putative contaminants in part B (based on filters 1-3) as well as the putative contaminants identified by the GBC model (filter 4). While some of the species will be falsely labeled as contaminants, we take a conservative approach and eliminate them from further analysis. The application of the GBC decontamination model reduced the number of unique hits, species and genera down to ∼41K, ∼4700, and ∼2200. It lowered the numbers of cell-containing and empty droplets with hits to ∼10K and ∼28K.

### Gaussian Mixture Model of alignments (Filter 5)

BLASTn outputs with E-values < 10^-5^ against the nt database are conventionally considered significant^98, 99^. In addition to E-value, BLASTn outputs two parameters that characterize the quality of each alignment, namely percent identity and the alignment length. By multiplying these two parameters and dividing by 100, we arrive at the total number of bases with a perfect match, which we refer to as n_match. Although all significant hits had E-values < 10^-5^, the distribution of n_match was a bimodal distribution (**SI Figure 3**). We use a Gaussian Mixture Model (GMM) to separate these two distributions. Because a species can have hits from both distributions, we systematically eliminated species if they were represented by more hits from the second distribution (μ_2_ = 91 bases) than the first (μ_1_ = 62 bases). We posit the first distribution represents lower-quality alignments that may be biologically interesting, as in the case of sequences from uncharacterized microbes with only a distant relative in the current nt database. They may also be technical artifacts. We take a conservative approach and favor precision over recall and impose this filter. A similar combination of filters has been used by Joint Genome Institute’s microbial genome annotation pipeline (E-value ≤ 1e-5, ≥ 90 percent identity, alignment ≥ 80 %)^102^ and by the Kraken team when benchmarking their tool against BLAST (E-value ≤ 10−5 and percent identity ≥90%)^99^. With the application of the GMM on the alignments, we arrived at ∼13K unique hits, ∼2300 species, and ∼1100 genera from ∼1K cell-containing droplets, and ∼11K empty droplets.

### Identifying putative habitats of origin for 500+ bacterial genera in the Tabula Sapiens Microbial dataset using the Genomes from Earth’s Microbiomes, the Human Microbiome Project, and the Unified Human Gastrointestinal Genome collection (Filter 6)

In addition to experimental (Filter 1) and bioinformatic decontamination (Filters 2-4) and alignment QC (Filter 5) steps, we wanted to capture information about the likely habitats of origin for TSM F5 taxa. We posit that some fraction of these species resides in the human microbiome’s external facing sites and have been observed in prior human microbiome studies, while the rest may be species that are transiently introduced via inhalation and consumption, among other possible means of transport, or may represent less frequently observed taxa in the human microbiome. To identify and separate members of these two groups, we leveraged the Genomes from Earth’s Microbiomes^7^ (GEM) dataset to gain insight into the likely habitats from which bacterial hits in the TSM F5 dataset originate. The GEM dataset is derived from 10,000+ metagenomes sampled from various habitats across the earth including terrestrial, aquatic, human and animal hosts, engineered environments, among others. This extensive dataset includes more than 12000 species-level operational taxonomic units.

We obtained the metadata for the GEM dataset from https://portal.nersc.gov/GEM/genomes/ (genome_metadata.tsv). While most bacterial genera were either strictly host-associated or free-living, some contained species that belonged to both habitat categories. Thus, to identify the dominant habitat of for each genus, we labeled species based on three categories: 1) “Human-associated” species include organisms found in the human microbiome; 2) “other_hosts” are species isolated from non-human vertebrates, invertebrates, fungi, and plants; 3) “free-living” are those species isolated from human-built or natural environments. With a table where each row represented a genus, and each column represented the fraction of its species belonging to one of these three categories based on the GEM dataset, we performed K-nearest neighbor clustering (number of clusters=3) and assigned each genus to a cluster (**SI Figure 4)**. We created a filter here called GEM_1 which is set to 0 (0=keep) for genera found in the human-associated cluster, and set to 1 (1=remove) if found in the other two clusters. GEM_1 is set to NA (NA=not applicable) if a genus is not found in the GEM dataset. We also created another filter called GEM_2 that functions in the same way as GEM_1 but for species. The combination of the two filters (GEM_1_2) allows for GEM_2 to override GEM_1. In other words, unless there is habitat information for a given TSM species, we assume the filter applied on to its genus.

In addition to the GEM dataset, we obtained habitat data for bacterial species from the Unified Human Gastrointestinal Genome (UHGG) collection^5^ and the Human Microbiome Project (HMP). The UHGG dataset is a large-scale and recent dataset of gut prokaryotes, comprising ∼200K non-redundant genomes from more than 4000 gut prokaryotes. These genomes are collected from humans living across 31 countries and 6 continents. This comprehensive catalog includes genomes from the Gastrointestinal Bacteria Culture Collection^29^, the Culturable Genome Reference^108^, as well as cultured human gut genomes from NCBI, PATRIC^31^ and IMG^109^ datasets, as well as metagenome-assembled genomes from Pasolli et al.^110^, and Nyfach et al.^7^ studies. We gathered the Unified Human Gastrointestinal Genome (UHGG) collection dataset from http://ftp.ebi.ac.uk/pub/databases/metagenomics/mgnify_genomes/human-gut/v2.0.1/ (genomes-all_metadata.tsv). We created a filter called UHGG_species, which is set to 0 (0=keep) if a TSM species has been identified in the UHGG dataset. It is otherwise set to NA.

The Human Microbiome Project (HMP) dataset acquired from https://www.hmpdacc.org/hmp/catalog/grid.php?dataset=genomic consisted of ∼3000 entries primarily from bacterial strains from various isolation sites. We chose species found in external-facing microbiomes (oral cavity, gastrointestinal tract, urogenital tract, airways, and the skin). When comparing the TSM dataset to these two datasets, entries with unknown taxonomic assignments were excluded. We created a filter called HMP_species, which is set to 0 (0=keep) if a TSM species has been identified in the HMP dataset. It is otherwise set to NA.

Filter 6 combines filters based on **H**MP, **U**HGG and G**E**M datasets, or what we refer to as the HUE datasets. Explicitly, the HUE filter (filter 6) is set to 1 (1=Remove) if any of the GEM_1_2, UHGG_species or HMP_species filters is equal to 1. The HUE filter will be set to 0 (0= Keep) if any of the three filters is equal to 0, effectively overriding the previous condition. Hence, if in one dataset a given species is found to be human-associated, and in another it has been found as not human-associated, we consider the species to be human-associated. The HUE filter is set to NA if all three filters it comprises are set to NA.

Based on the HUE filter,159 species were determined to be human-associated and the rest originated from other hosts or environments, or were not found in these datasets. To obtain putative habits of origin for those species in the NA category, we used OpenAI’s Davinci large language model described in the next section. Removing bacterial species and genera based on their habitat of origin in the HUE datasets further reduced the number of unique hits, species, and genera down to ∼10K, ∼1700, and ∼900, respectively, originating from 1K cell-containing droplets and 8K empty droplets. Please see **Data and Materials Availability** for the Jupyter notebook where further details for incorporating the GEM, UHGG, and HMP datasets into TSM dataset can be found.

### Benchmarking and employing Davinci large language model for identification of putative habitats for 1500+ microbial species (Filter 7)

The HUE datasets provide putative habitat information for many taxa, however they do not include all taxa found in the TSM dataset. Specifically, there were 1500+ species for which filter 6 (the HUE filter) does not provide any information. To systematically retrieve information on where these species have previously been isolated from, we employed openAI’s Davinci Large Language Model (text-davinici-003 accessed Jan 2023). We simply asked the following question for each species: “provide a short summary of where species_i has been isolated from”, where species_i corresponds to the species of interest. To make queries programmatically, we employed openAI’s API and set max_tokens=200. We used the following default parameters: temperature=0.5, top_p=1, frequency_penalty, presence_penalty=0.

To better understand the model’s level of self-consistency, we randomly chose 10 species and posed the same prompt three times. A table of model outputs for all three runs is provided in **SI Table 16** and shows that the model presents a certain level of variability across multiple outputs from which we can identify high-confidence segments based on consensus across all three outputs. For example, here are the three outputs for *Acidaminococcus intestini*: Output 1: “Acidaminococcus intestini has been isolated from the gut of humans and pigs, as well as from soil samples and wastewater”. Output 2: “Acidaminococcus intestini has been isolated from the intestinal tracts of humans, pigs, chickens, and horses. It has also been found in soil, water, and sewage”. Output 3: “Acidaminococcus intestini has been isolated from the intestinal tracts of humans, pigs, and chickens. It has also been detected in the feces of horses and cows, as well as in the environment”.

Next to test the validity of the model’s outputs, we randomly selected 100 species from the GEM dataset, providing us a subset of habitats that a given species has been previously isolated from. By running three independent queries per species and combining the model’s outputs, we were able to estimate the model’s precision and recall. First, we created two categories, based on whether a given species was previously isolated from the human microbiome or not (i.e. species isolated from natural environments, human-built environments, or non-human hosts). Then we grouped Davinci’s responses based on the same two categories. Using GEM as the truth dataset, we found that the model was able to identify 59 out of 63 species previously found in the human microbiome. Out of 37 species not found in the human microbiome, the model correctly identified 25. The model outperformed the truth dataset in some cases. For example, among the model’s false positives, we noticed four species that had been found previously in the human body by other studies (*Serratia plymuthica*^111^, *Leifsonia sp.*^112^, *Rickettsia japonica*^113^, and *Lysobacter oculi*^114^). Correcting for these four species, we arrive at a precision of 89% and a recall of 94%. We provide a table of these 100 test species along with GEM habitat categories and model outputs (**SI Table 17)**. To increase the precision further, we only accepted species that had been identified as having been found in humans by all three Davinci outputs. Again, *Serratia plymuthica* and *Lysobacter oculi* were both found in humans by all three Davinci outputs and thus were counted as true positives. We arrived at a precision of 98% and a recall of 75% (see **SI Figure 1.e** for the confusion table). Details of programmatic query submission along with benchmarking can be found in a Jupyter notebook listed under **Data and Materials Availability**.

It is important to state some of the limitations of the current Davinci model. First, we found the model’s self-consistency depends on the prompt. Generally, the simpler and more open-ended prompts provided better, less biased outputs. Additionally, while the model performs well at classifying species into these two broad categories, we observed that it performs poorly if the task is identifying detailed habitat information. We noticed it can generate false results interspersed with true results which are difficult to distinguish. This is currently a common drawback of generative models in all areas of natural language processing, referred to as model “hallucination”. Thus, although we provide the model’s combined outputs under “davinci_combined” column for each species, we suggest readers perform independent search for obtaining more detailed habitat information.

Having benchmarked the Davinci model against this prompt, we then applied the model to the dataset of 1500+ species. To ensure high-confidence classifications, we set this filter to 0 (0=keep) if for a given species, all three Davinci outputs independently state that the species has been found previously in the human body. The filter is otherwise set to 1 (1=remove). We found that ∼100 species met this criterion. The full dataset with Davinci model outputs is provided as (**SI Table 18**). Imposing this filter, we arrived at ∼3K unique hits, ∼300 species, and ∼200 genera from the three biomes of interest. These hits originated from ∼500 cell-containing (∼300 of which have cell type annotation) and ∼2K empty droplets.

### Impact of filters on the Tabula Sapiens Microbiome dataset

The TSM dataset consisting of significant hits (E-values <10^-5) from viruses, bacteria and fungi is referred to as TSM F0. The TSM dataset that has gone through the first filter will be referred to as the TSM F1, and so on, such that TSM F7 is the version of the dataset that has passed through all seven filters. In addition to droplets with TSM F0 hits, we kept track of droplets (cell-containing or empty) that did not produce any significant hits. The combination of this group of cells and the TSM F0 group of cells is referred to as the TSM F00.

F00: The total number of cell-containing and empty droplets analyzed were ∼1.3M and ∼156M, respectively.

TSM F0: There were ∼724K significant unique hits (counted based on Unique Molecular Identifiers or UMIs) from three biomes spanning bacteria, fungi and viruses with E-values < 10^-5. These hits originated from ∼79K cell-containing droplets (∼6% of F00 cell-containing droplets) and ∼380K empty droplets (∼0.2% of F00 empty droplets).

TSM F1: When we excluded all species found in negative control samples, the number of unique hits was reduced to ∼63K, or roughly 10% of the F0 hits. Interestingly, the number of species dropped from ∼8200 to only ∼7100 and the number of genera went down from ∼2500 to ∼2400. Filter 1 reduced the numbers of cell-containing and empty droplets to ∼13K (∼16% of F0 cell-containing droplets) and ∼44K (∼12% of F0 empty droplets), respectively.

TSM F2: The removal of Salter et al. bacterial genera, reduced the number of unique hits, species, and genera to ∼52K, ∼6200, and ∼2300, respectively. It reduced the numbers of cell-containing and empty droplets to ∼12K and ∼35K.

TSM F3: The removal of putative fungal contaminants previously identified by Narunsky-Haziza *et al.*, reduced the number of species to ∼6100. Other numbers remained nearly the same.

TSM F4: The application of the GBC decontamination model, reduced the number of unique hits, species, and genera down to ∼41K, ∼4700, and ∼2200. It lowered the numbers of cell-containing and empty droplets with hits to ∼10K and ∼28K.

TSM F5: With the application of the gaussian mixture model on the alignments, we arrived at ∼13K unique hits, ∼2300 species, and ∼1100 genera from ∼1K cell-containing droplets, and ∼11K empty droplets.

TSM F6: Removing bacterial species and genera based on their habitat of origin in the HUE datasets further reduced the number of unique hits, species, and genera down to ∼10K, ∼1700, and ∼900, respectively, originating from 1K cell-containing droplets and 8K empty droplets.

TSM F7: Lastly, removing bacterial, viral and fungal species that were not previously isolated from the human microbiome based on three independent outputs of OpenAI’s Davinci model, we arrived at ∼3K unique hits, ∼300 species, and ∼200 genera from the three biomes of interest. These hits originated from ∼500 cell-containing (∼300 of which have cell type annotation) and ∼2K empty droplets.

The exact numbers and fractions of remaining unique hits, species, genera, cell-containing and empty droplets after each filter relative to TSM F0 dataset is shown in **SI Table 13** and **SI Figure 5**. With the imposition of all filters, we arrive at ∼0.4% of the unique hits found in the TSM F0 dataset, ∼3% of the species and ∼5% of the genera. Similar to the percentage of unique hits remaining, the percentage of cell-containing and empty droplets is ∼0.7% and ∼0.4%, respectively. If we consider the percentage of *total* cell-containing and empty droplets (i.e. compare TSM F7 dataset to the TSM F00 dataset), we see that ∼0.04% of cell-containing droplets and ∼0.001% of empty droplets remain. Based on these numbers, we can conclude cell-containing droplets are ∼40x more likely to harbor microbial reads than empty droplets.

Considering that the starting number of sequences in the TSM dataset was ∼70 billion (including human reads) and the final number of unique microbial hits passing through all filters was a few thousand, we conclude that finding high-confidence microbial sequences in human cellular transcriptomes is a search roughly for one in 25 million sequences. Considering the less stringent criteria imposed on the TSM F5 dataset, it is a search for one in 2.5 million sequences. These numbers are a function of various parameters and filters affecting the recall of our search. Thus, they could be a few orders of magnitude improved with experimental assays designed specifically for capturing microbial transcripts as well as other computational improvements.

### Exploring the inter-and-intra donor tissue microbiomes

To explore the inter-and-intra donor tissue microbiomes, we measure the species unique fractions. For example, for donor 2, when we sum the number of species that only appear in one tissue and divide by the total number of species identified in that donor, we arrive at the unique species fraction equal to 0.53. The average intra-donor unique fraction is 0.64 and the average inter-donor unique fraction, calculated for one tissue across multiple donors, is 0.76 (**SI Figure 17**). The personal nature of human tissue microbiomes, at least in part, likely reflects the highly individual-specific microbiomes that exist at external-facing microbiomes^115–117^. As we also demonstrate through the overlap between species from different isolation sites in the HMP database, external-facing microbiomes have limited overlap at the species level **(SI Figure 22)**. Moreover, as we reduce the taxonomic resolution from species to orders and phyla, the correlations between and within individuals increase considerably (**SI Figures 18-19**), suggesting that high-level tissue microbiome compositions are largely conserved.

### Single-droplet statistics, outlier cells and outlier cell types

In the population of hit-positive TSM F7 cells, cells with one unique hit dominate the distribution. The median number of uhits/cell is 1, the mean is ∼1.5 and the standard deviation is 8. See **SI Table 23** for statistics on cell-containing and empty droplets from TSM F7 dataset. We define outlier cells by those with uhits/cell > 1.5xIQR, where IQR is the interquartile range for the uhits/cell distribution. The outlier cells are shown in **SI Table 24**. The most prevalent species in outlier cells was EBV. It appeared in outlier cells almost exclusively from donor 2 (TSP2), with the exception of one cell from the Salivary gland of TSP7. TSP2 outlier cells positive for EBV hits consisted of bladder B cells, Lymph node Plasma cells and B cells, among others. In addition to EBV in TSP2, we found a bladder macrophage containing hits from *Ureaplasma urealyticum*^118^, which is a commensal organism of the urogenital tract and associated with premature birth and adverse pregnancy outcomes. We found hits to *Aggregatibacter aphrophilus*^119^, an oral bacteria associated with periodontal disease and endocarditis, in an adventitial cell of the salivary gland in TSP7.

Six out of eight cell types associated with outlier cells were immune cells, namely Plasma cell, B cell, Natural Killer cell, Macrophage, Memory B cell, and CD34+ CD38-Multipotent T cell. Interestingly, 72% of outlier cells had viral hits (exclusively EBV) compared to only 7% of empty droplets (**SI Table 25**). Nearly one quarter of outlier cells had bacterial hits compared to ∼83% of outlier empty droplets. Fungal hits appeared in only a small fraction of outlier cells and droplets. These numbers support the hypothesis that bacterial and fungal hits mostly come from extracellular spaces and are co-localized with cells or passively uptaken, whereas in cases of intracellular pathogens such as EBV, one might expect a higher number of positive cells and a higher number of unique hits per cell. We found five cells that harbored unique hits from more than one species, primarily bacteria from different orders and phyla in a single cell.

We grouped cells with TSM F7 hits by their cell type, and through dividing the number of F7 hit-positive cells by the total number of cells, we measured the fraction of hit-positive cells for each cell type. To have sufficient numbers for statistical analysis, we chose only cell types with a minimum of 10 cells that were sequenced and annotated. The mean and standard deviation of the fraction of cells in each cell type positive for F7 hits were ∼0.0006 and ∼0.0015%, respectively. We identified outlier cell types as those that have a higher fraction of F7 positive cells than 1.5xIQR. There were 195 cell types that were included in the analysis, from which 36 were identified as outlier cell types (**SI Table 26**). Among the outliers, ∼60% of cell types belonged to the immune compartment. Some examples of non-immune outlier cell types are lung alveolar epithelial cells, lung ciliated cells, intestinal transient amplifying cells, vascular and lymph endothelial cells.

## SI Figures

**SI Figure 1.**
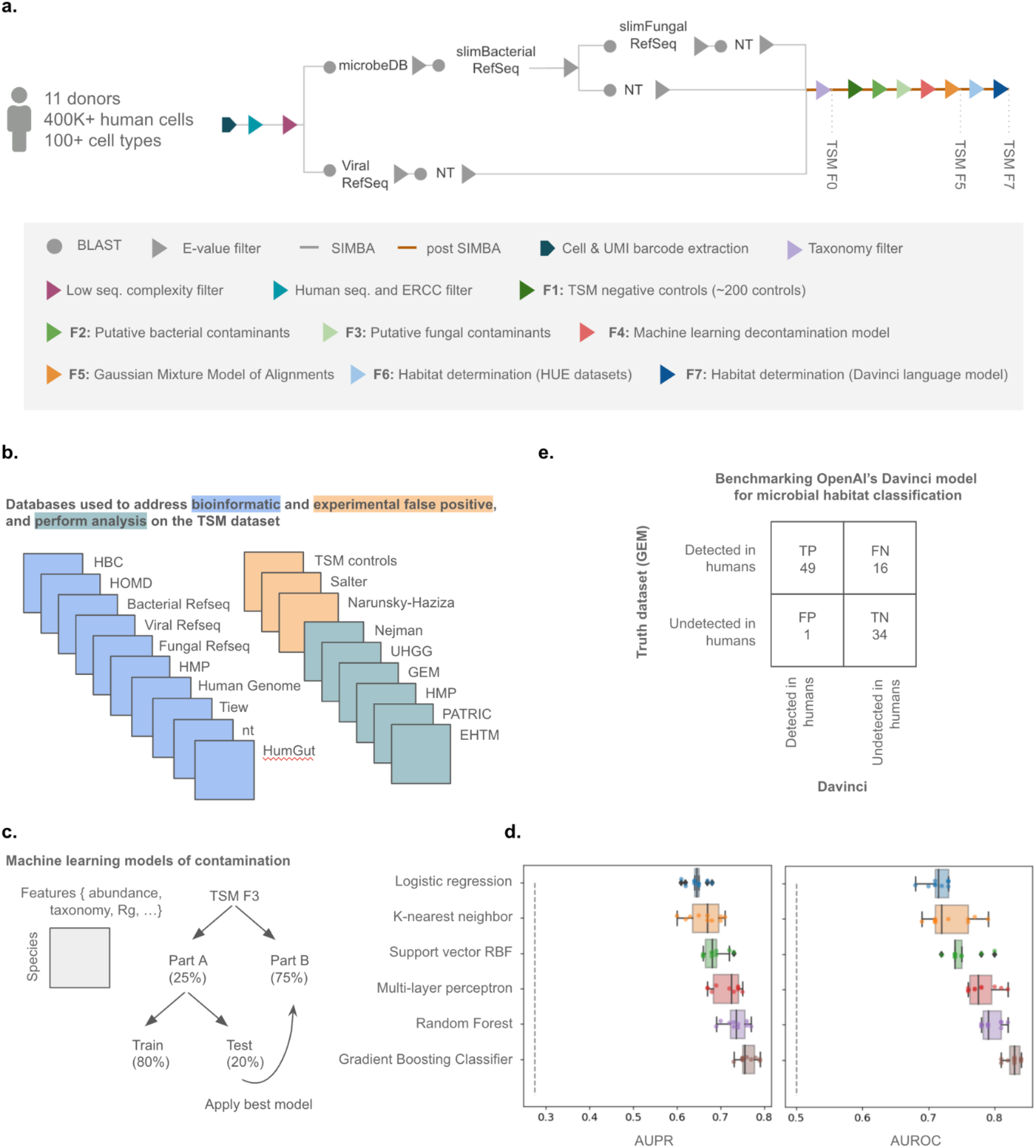
Computational pipeline and post-processing steps developed for the identification of high-confidence bacterial, viral and fungal sequences in human single-cell data. **a)** The computational steps comprising SIMBA and post SIMBA filtration of the Tabula Sapiens Microbiome (TSM) dataset. **b)** Databases used to mitigate and benchmark bioinformatic false positives (blue) and experimental false positives (orange), as well as those used for microbial flow and other analyses (green). HBC: Human Gastrointestinal Bacteria Culture Collection, HOMD: Human Oral Microbiome Dataset, Tiew *et al.* (gut mycobiome), HMP: Human Microbiome Project, nt, Bacterial, Fungal and Viral Refseq datasets (all publically available from NCBI), TSM controls (∼200 instrument and reagent negative controls), Salter *et al.* (bacterial genera previously found in prior sequencing studies), Narunsky-Haziza *et al.* (fungal contaminants seen in human tumor samples), Nejman *et al.* (bacterial taxa previously seen in human tumor samples), UHGG (Unified Human Gastrointestinal Genome collection), GEM (Genomes from Earth’s Microbiomes), PATRIC (the Pathosystems Resource Integration Center), and EHTM (Extracted Human Tissue Microbiome). **c)** The process by which we used known and putative contaminants to identify additional putative contaminants in the TSM dataset using 6 different types of machine learning models. **d)** Performance metrics of each machine learning model type on withheld test data. For each model type, 10 different models using different train/test splits were created and AUPR and AUROC scores were measured. Dashed lines correspond to null model performance (0.5 for AUROC and the prevalence of the positive class for AUPR). **e)** Confusion matrix for benchmarking the Davinci large language model for microbial habitat classification against the GEM dataset using 100 randomly selected species (Precision = 98%, Recall = 75%). Species are categorized into two groups: 1) found in the human body 2) found anywhere else but the human body. TP: True Positive, FP: False Positive, FN: False Negative, TN: True Negative.

**SI Figure 2.**
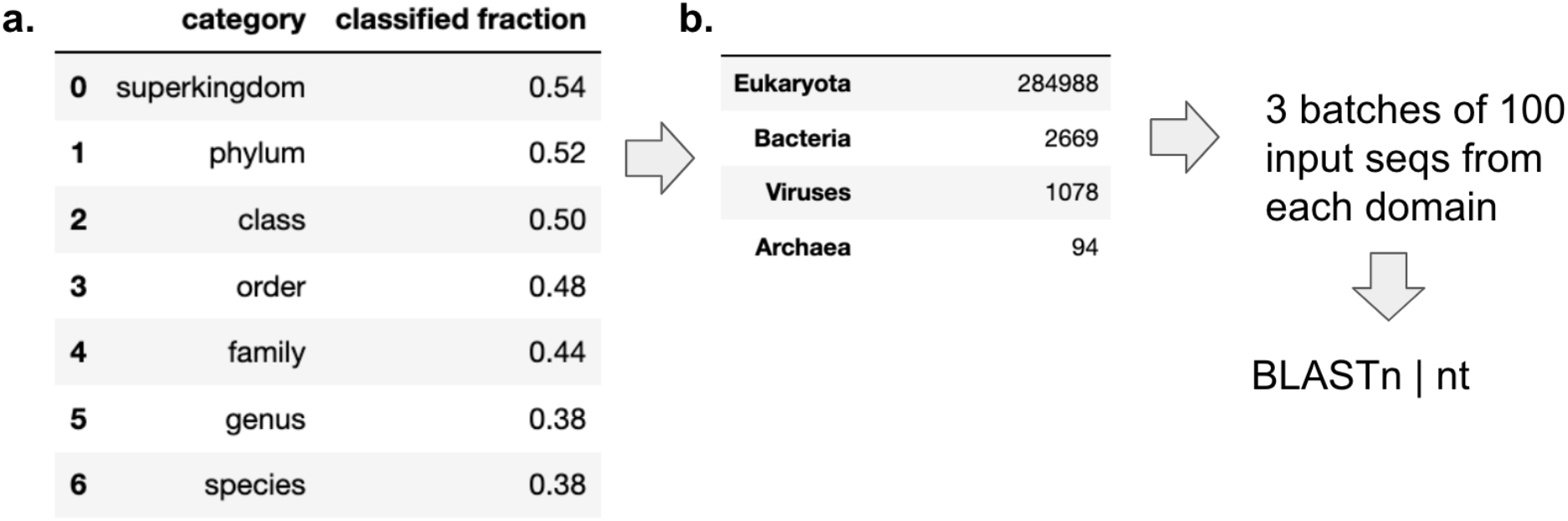
Kraken2 (against nt) classification assignments for a Tabula Sapiens Microbiome sample (Bladder, TSP2). **a)** Fraction of classified input sequences by Kraken2 at different taxonomic levels. **b)** Number of classified sequences that were assigned to each domain.

**SI Figure 3.**
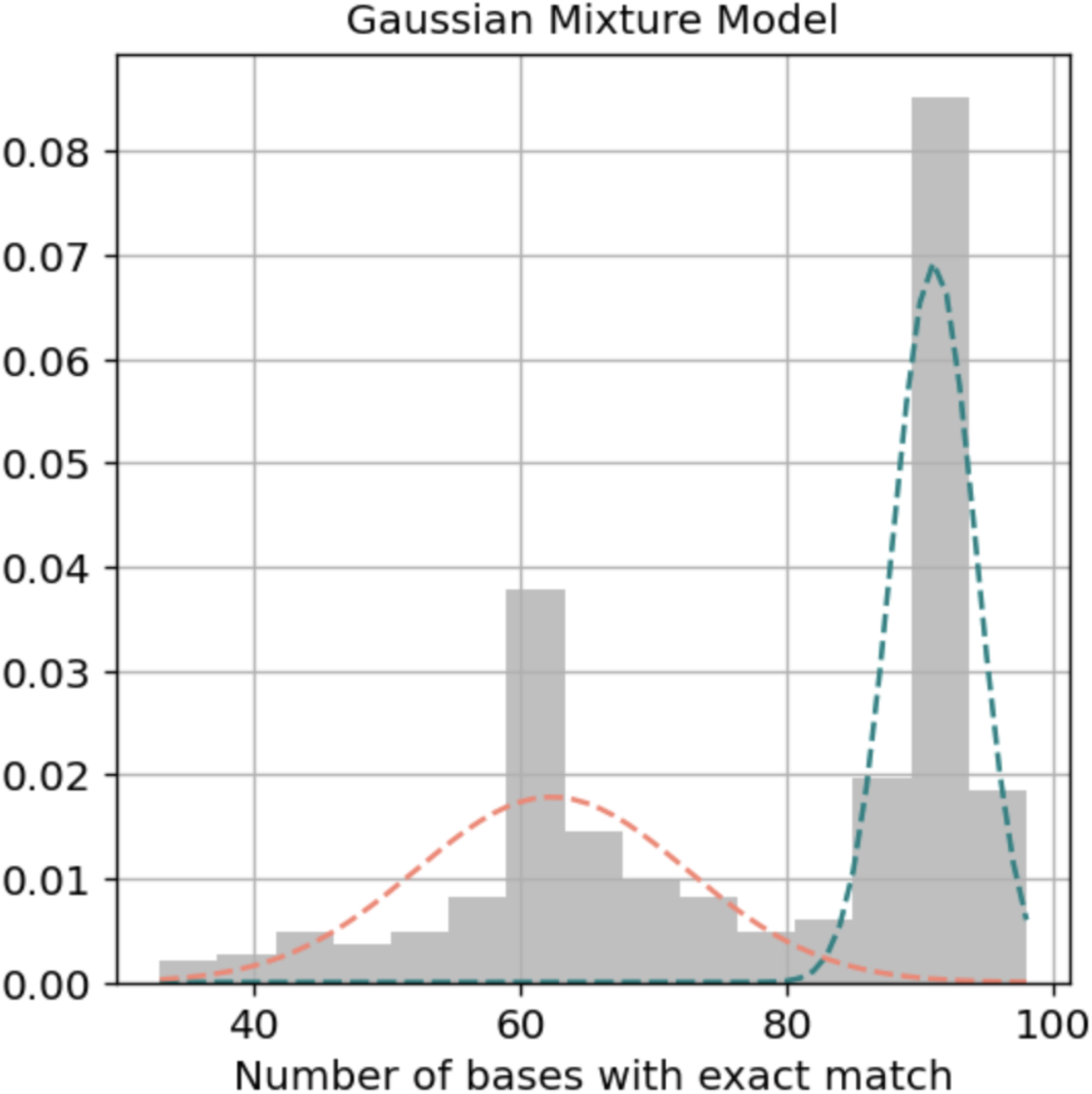
Number of bases with exact match to a reference sequence in the nt database (n_match). The Gaussian Mixture model is superimposed on the distribution of n_match (mean or μ_’_= 62 bases for first distribution, and μ_&_= 91 bases for the second distribution shown in green).

**SI Figure 4.**
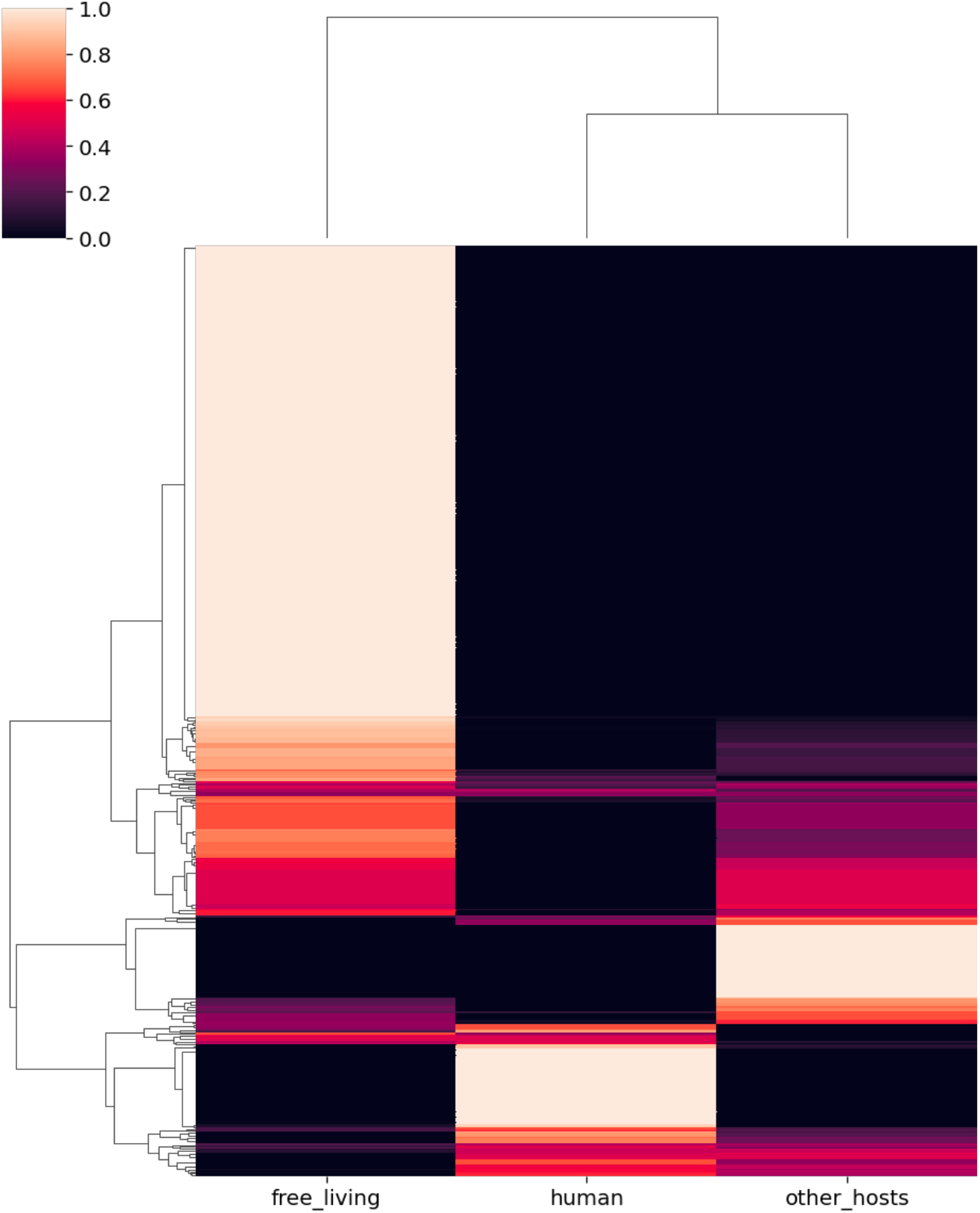
Clustering of bacterial genera based on the fraction of their species belonging to three course-grained labels: free-living (e.g. aquatic, terrestrial), human-dwelling (e.g. gi tract, skin), and other hosts (e.g. plants, insects). Each row represents a bacterial genus from the TSM dataset that also appears in the GEM dataset.

**SI Figure 5.**
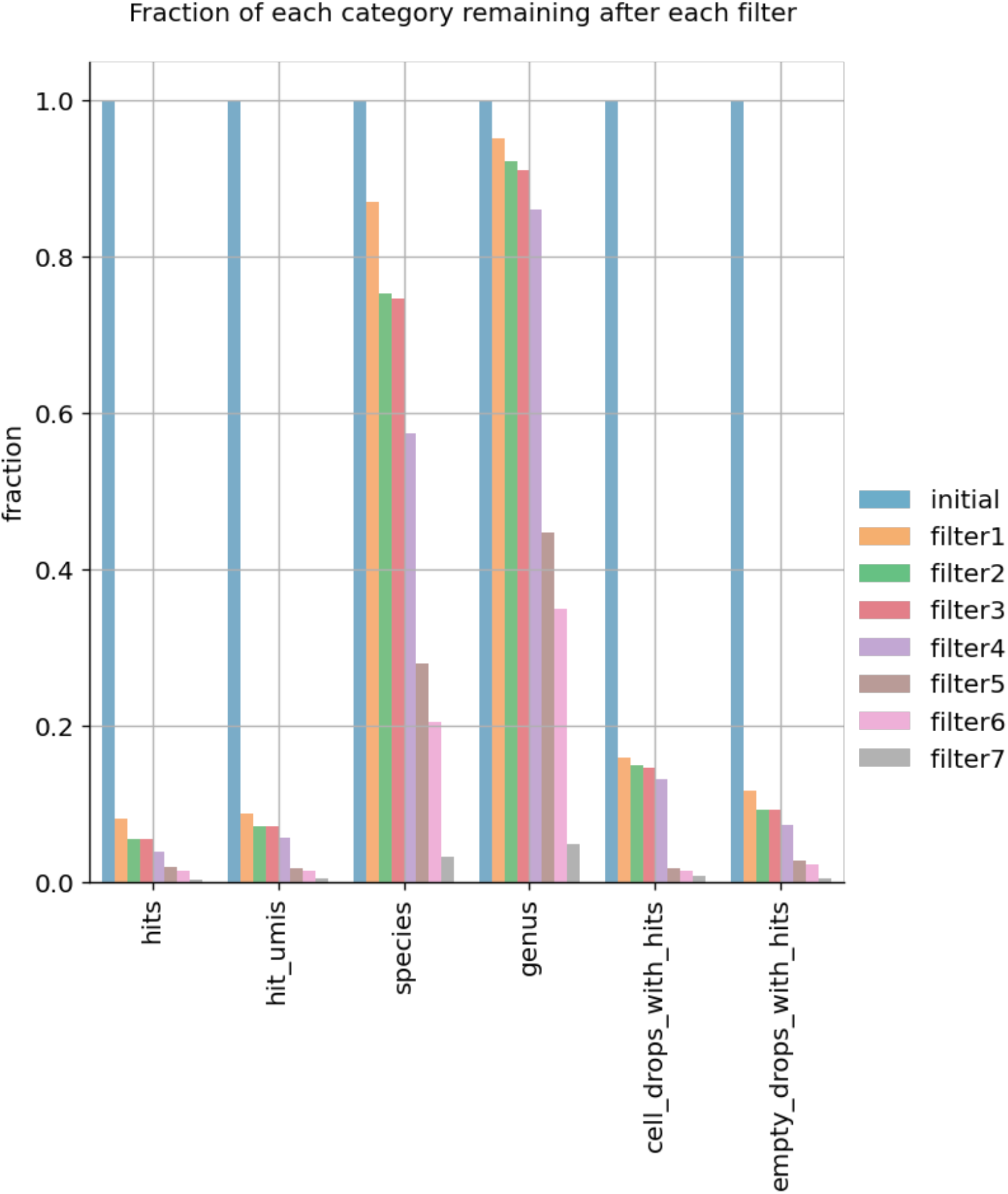
Fraction of total microbial hits, unique hits (based on UMIs), species, genera, cell-containing droplets with hits, and empty droplets with hits remaining after each filter is applied. Filter 1: ∼200 negative instrument and reagent controls, Filter 2: removal of Salter *et al.* genera, Filter 3: removal of putative fungal contaminants identified by Narunsky-Haziza *et al.*, Filter 4: applying the Gradient-boosting machine learning model for decontamination, Filter 5: applying a Gaussian Mixture Model to select species with higher quality alignments, Filter 6: putative habitat determination using HUE datasets and selecting those taxa more likely to be residents of the human microbiome, and Filter 7: putative habitat determination using the Davinci Large Language Model and selecting additional taxa that are likely residents of the human microbiome.

**SI Figure 6.**
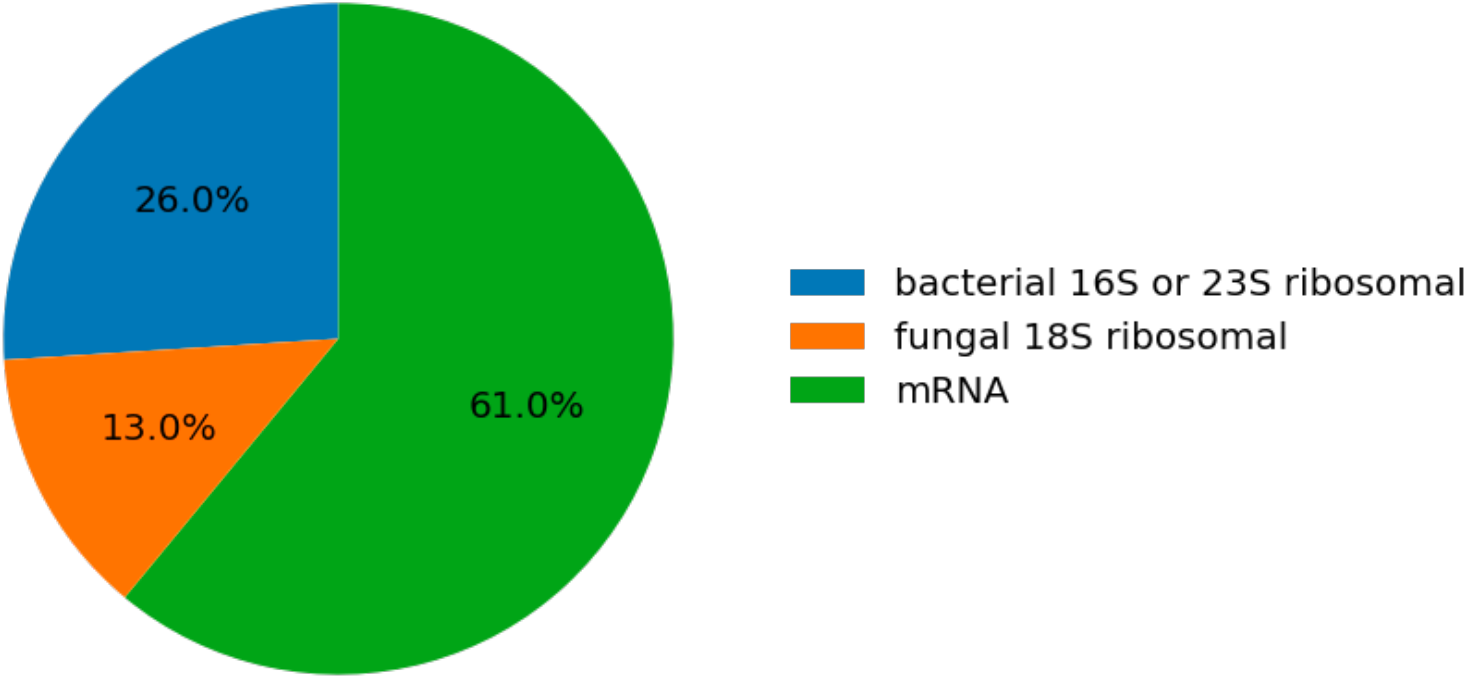
Percentage of different types of microbial RNA discovered in the TSM F7 dataset.

**SI Figure 7.**
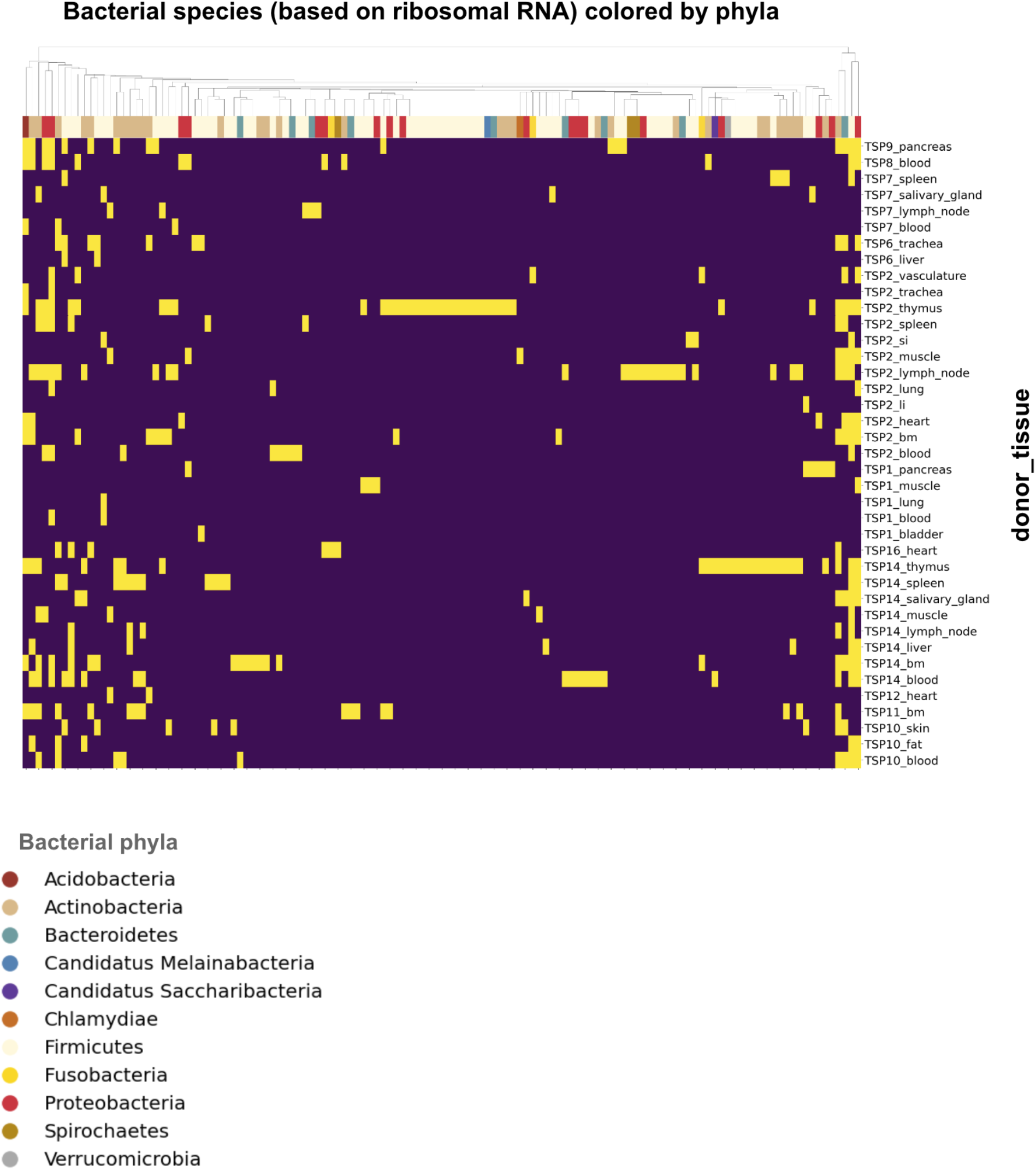
Ribosomal RNA from different bacterial phyla across various tissue types. Clustered heatmap of ribosomal hits (presence/yellow and absence/blue is shown) found in each donor’s tissue sample. Bacterial species are color-coded based on their phylum-level classification. bm: bone marrow, li: large intestine, si: small intestine.

**SI Figure 8.**
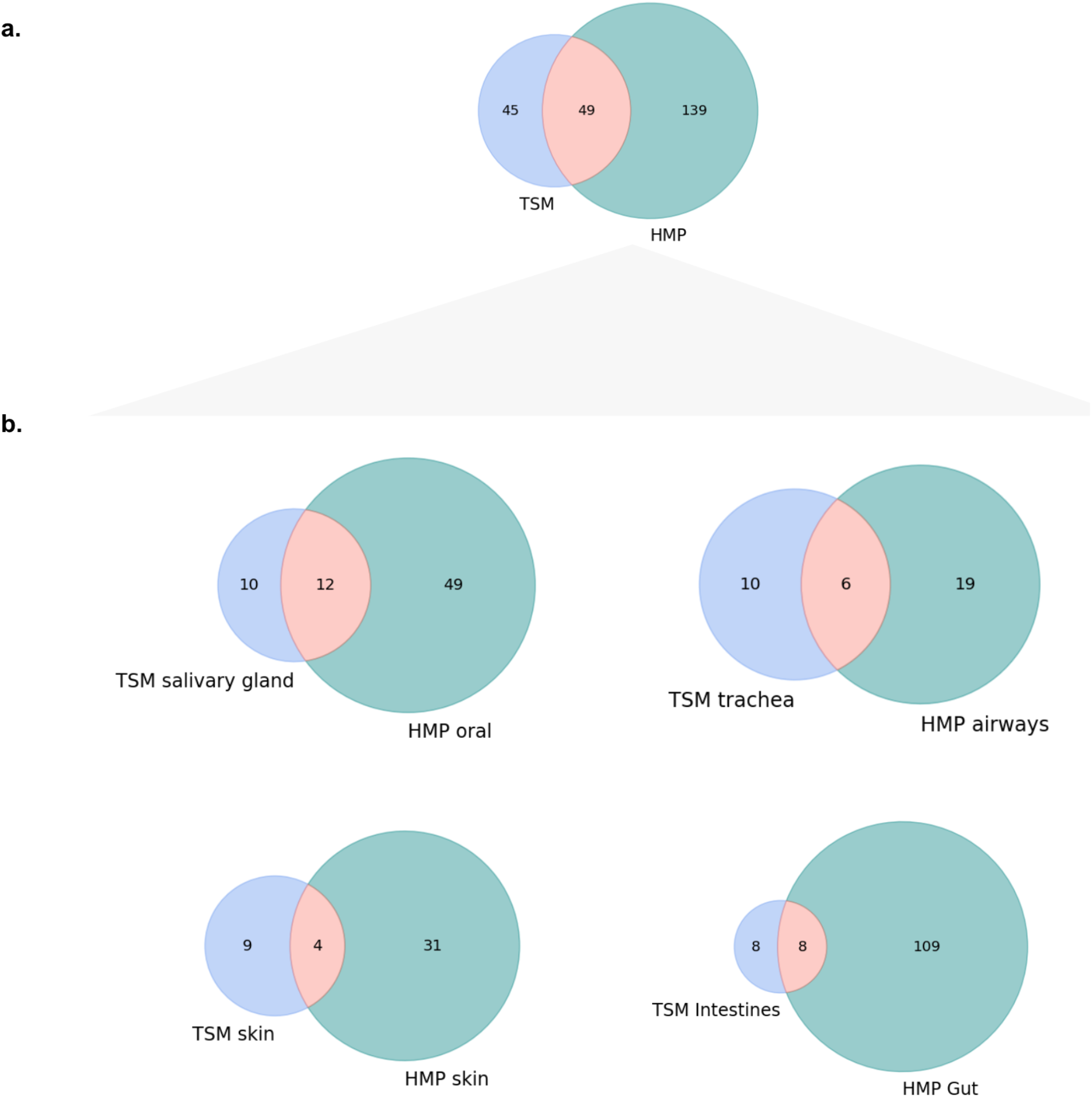
Comparisons between the Tabula Sapiens Microbiome (TSM F7) dataset and the Human Microbiome Project (HMP). a) Number of bacterial genera shared between TSM F7 and HMP, b) broken down by HMP isolation sites and their closest tissue counterpart in the TSM dataset. We exclude any members of either dataset that do not have a taxonomic assignment at the genus level in this comparison. TSM intestines refers to both the small and the large intestine samples.

**SI Figure 9.**
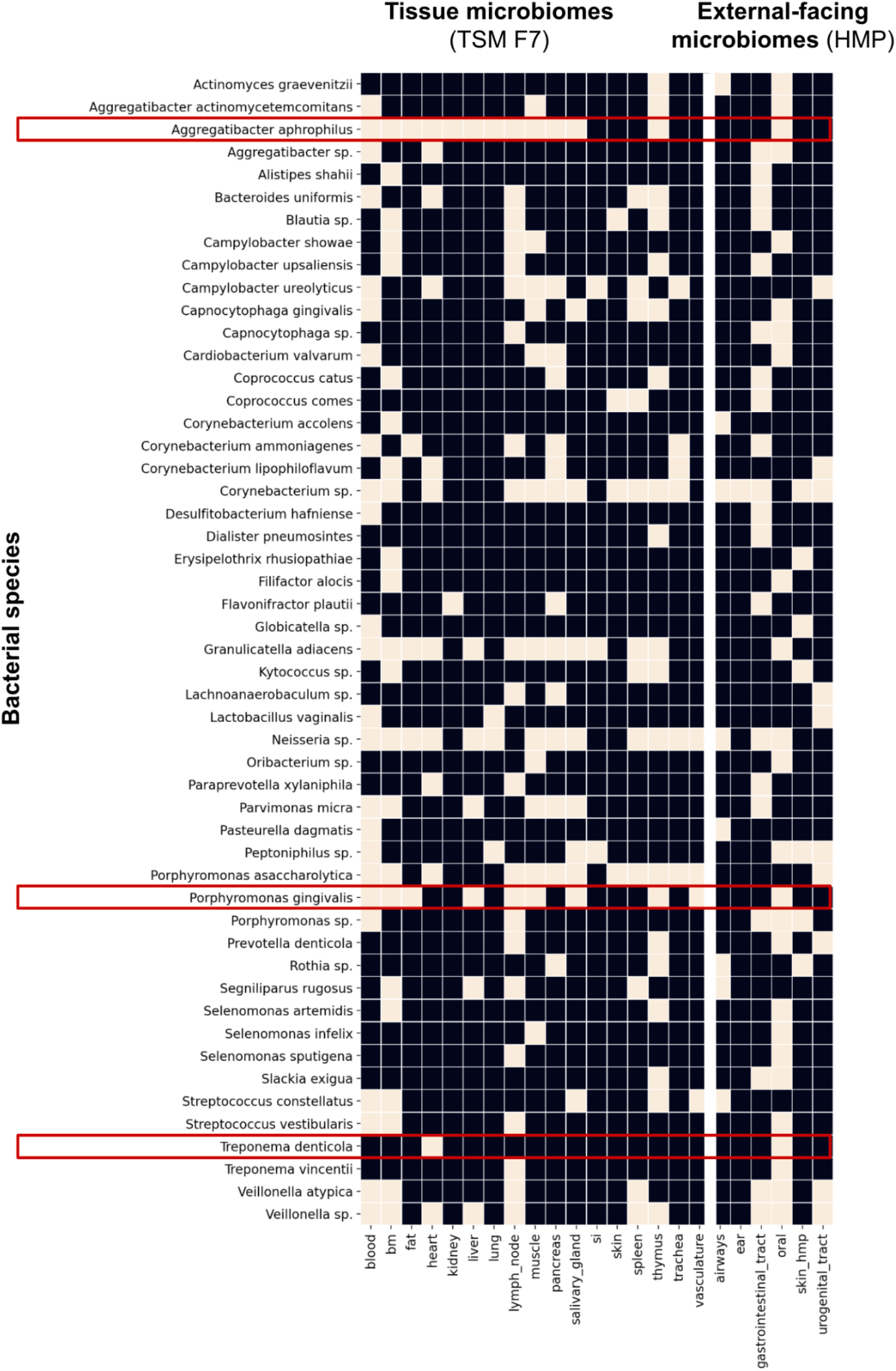
Bacterial species found in both the TSM F7 dataset and the Human Microbiome project (HMP). Presence is denoted by the light colored boxes. Boxed in red are species that were referred to in the main text. bm: bone marrow, li: large intestine, si: small intestine.

**SI Figure 10.**
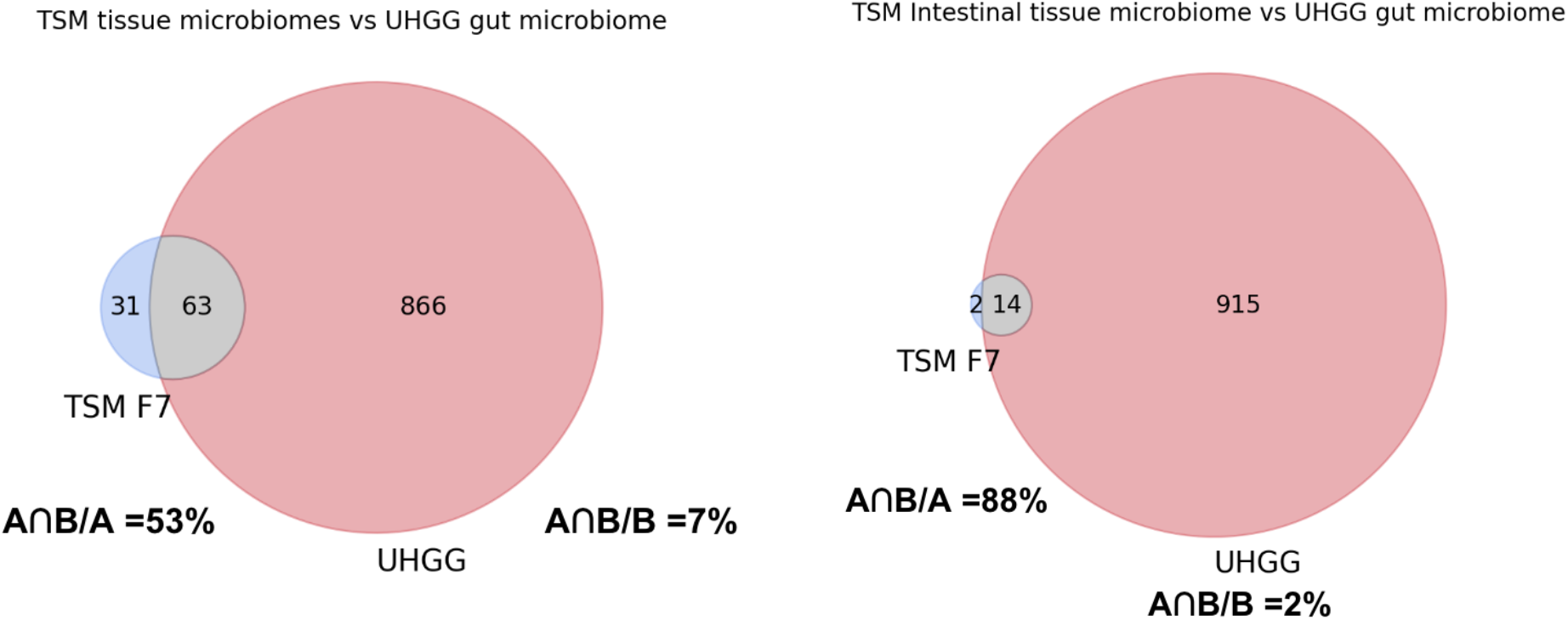
Comparisons between the Tabula Sapiens Microbiome (TSM f7) and the Unified Human Gastrointestinal Genome (UHGG) collection datasets. left) Number of bacterial genera shared between TSM F7 tissue microbiomes and UHGG. right) Number of bacterial genera shared between TSM F7 intestinal tissue microbiomes (both large and small intestines) and UHGG. We treat TSM datasets as set A and UHGG as set B.

**SI Figure 11.**
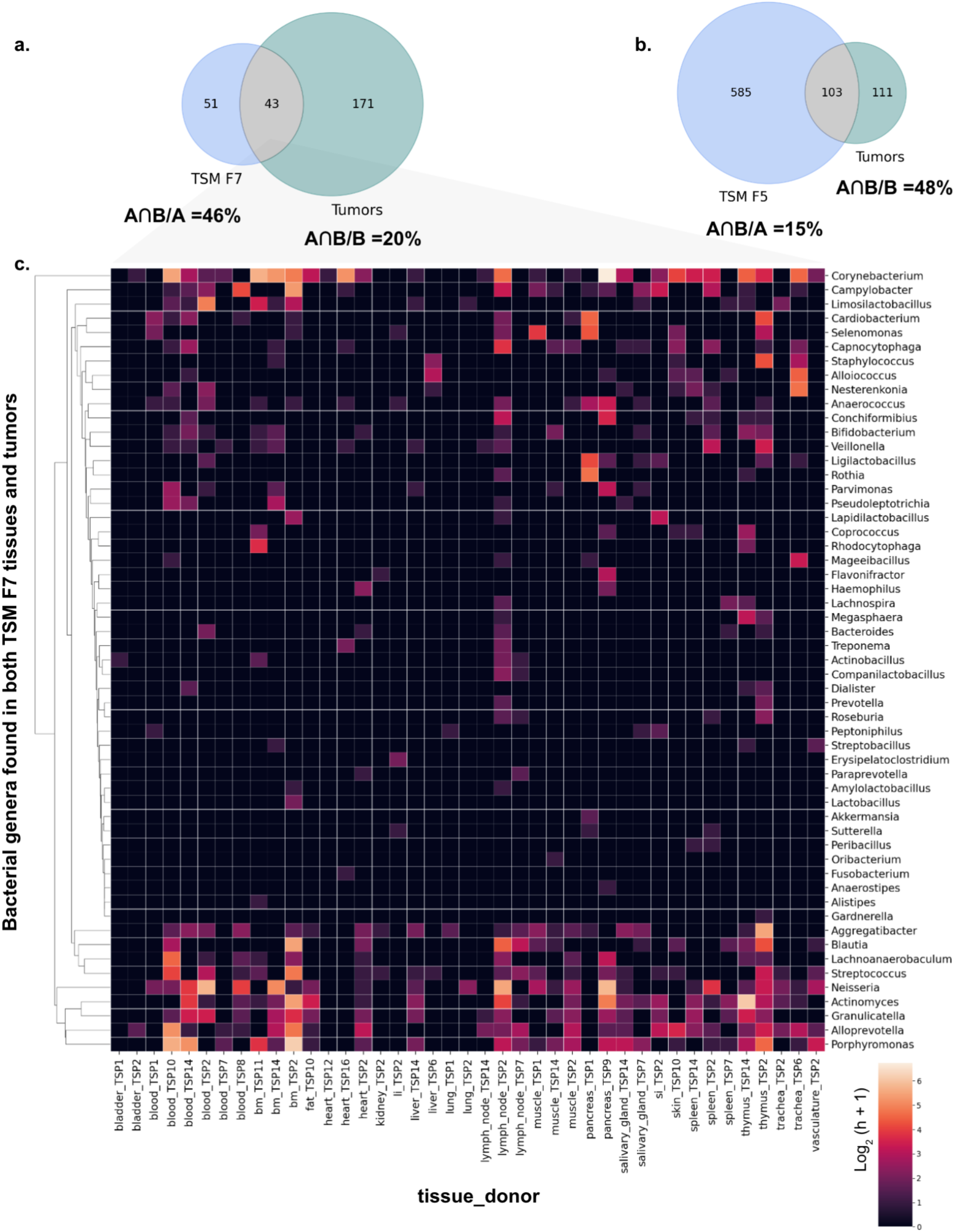
Exploring the intersection of bacterial genera found in the Tabula Sapiens Microbiome (TSM) a) F7 or b) F5 datasets and tumor microbiomes from Nejman *et al.* (only species and orders appearing in tumors that passed all of this study’s contamination filters were selected for analysis, refer to this study’s Table S4). The percentage of shared genera compared to total number of genera in each dataset are shown (set A corresponds to TSM datasets, and set B corresponds to the Nejman *et al.* dataset). c) abundance of bacterial genera shared between the TSM F7 stratified by tissue and donor of origin and Nejman *et al.* datasets. Abundance of genera is based on Log_2_(hits +1). Entries with unknown genera are excluded from comparisons shown in the venn diagrams. bm: bone marrow, li: large intestine, si: small intestine.

**SI Figure 12.**
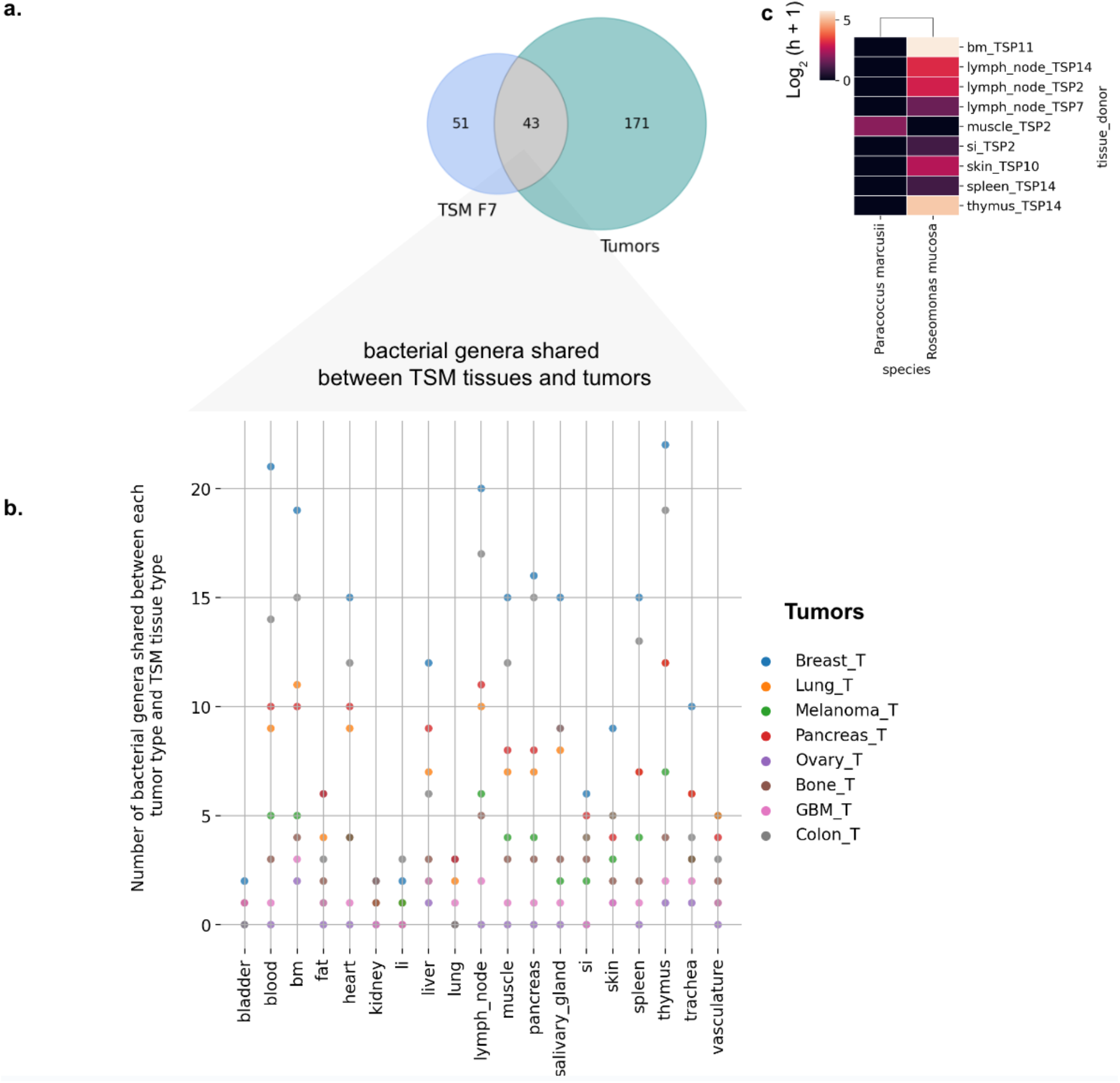
Comparison of bacterial genera in healthy tissues from TSM F7 and tumors from Nejman *et al.* (Only genera appearing in tumors that passed all of this study’s contamination filters were selected for analysis from Nejman *et al*. Table S4, and from their Figure 4D for tumor enriched species). a) venn-diagram representing the number of shared genera between the two datasets. b) Number of bacterial genera shared between each tumor type and each TSM F7 tissue type is shown. bm: bone marrow, li: large intestine, si: small intestine, GBM: glioblastoma multiforme. c) abundance of species that were highly enriched in tumors and also appeared in the TSM F5 dataset. Abundance of species is based on Log_2_ hits +1.

**SI Figure 13.**
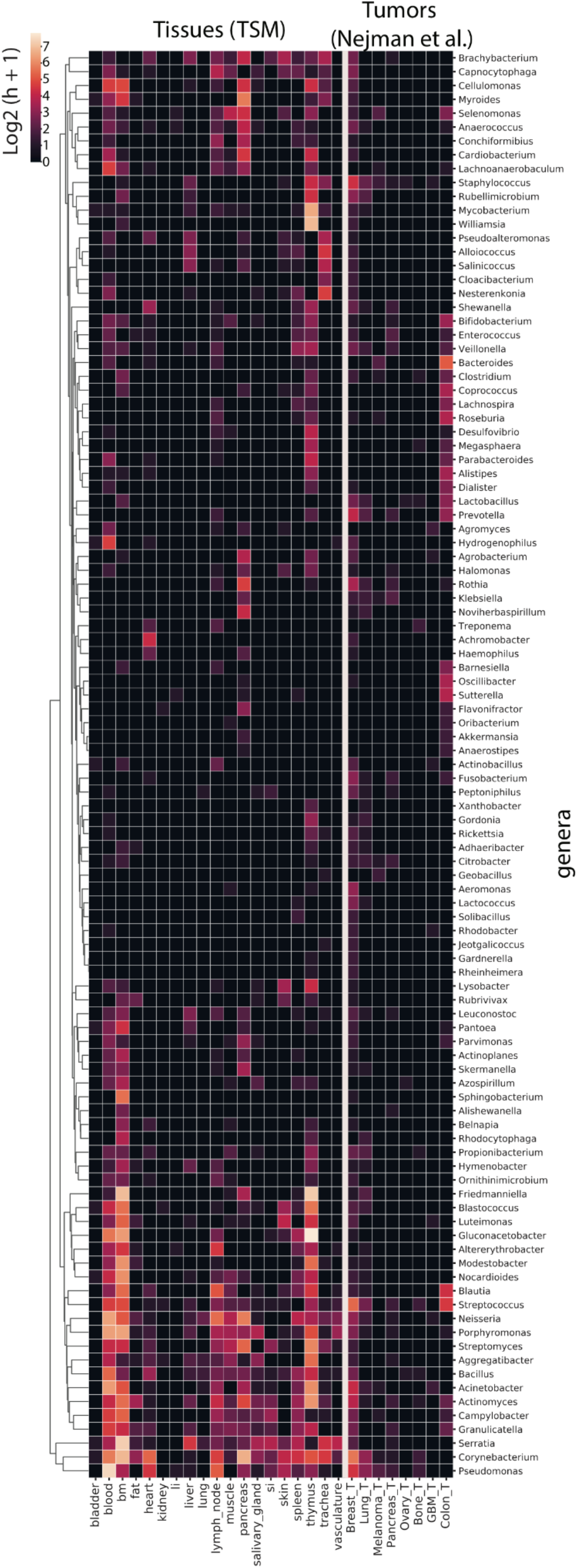
Abundance of bacterial genera previously found in tumor samples (Nejman *et. al*, 2020) and in the Tabula Sapiens Microbiome (TSM) F5 dataset stratified by tissue of origin. Abundance is based on log_2_(hits +1). Samples with “_T” appended correspond to tumors. bm: bone marrow, li: large intestine, si: small intestine, GBM: glioblastoma multiforme.

**SI Figure 14.**
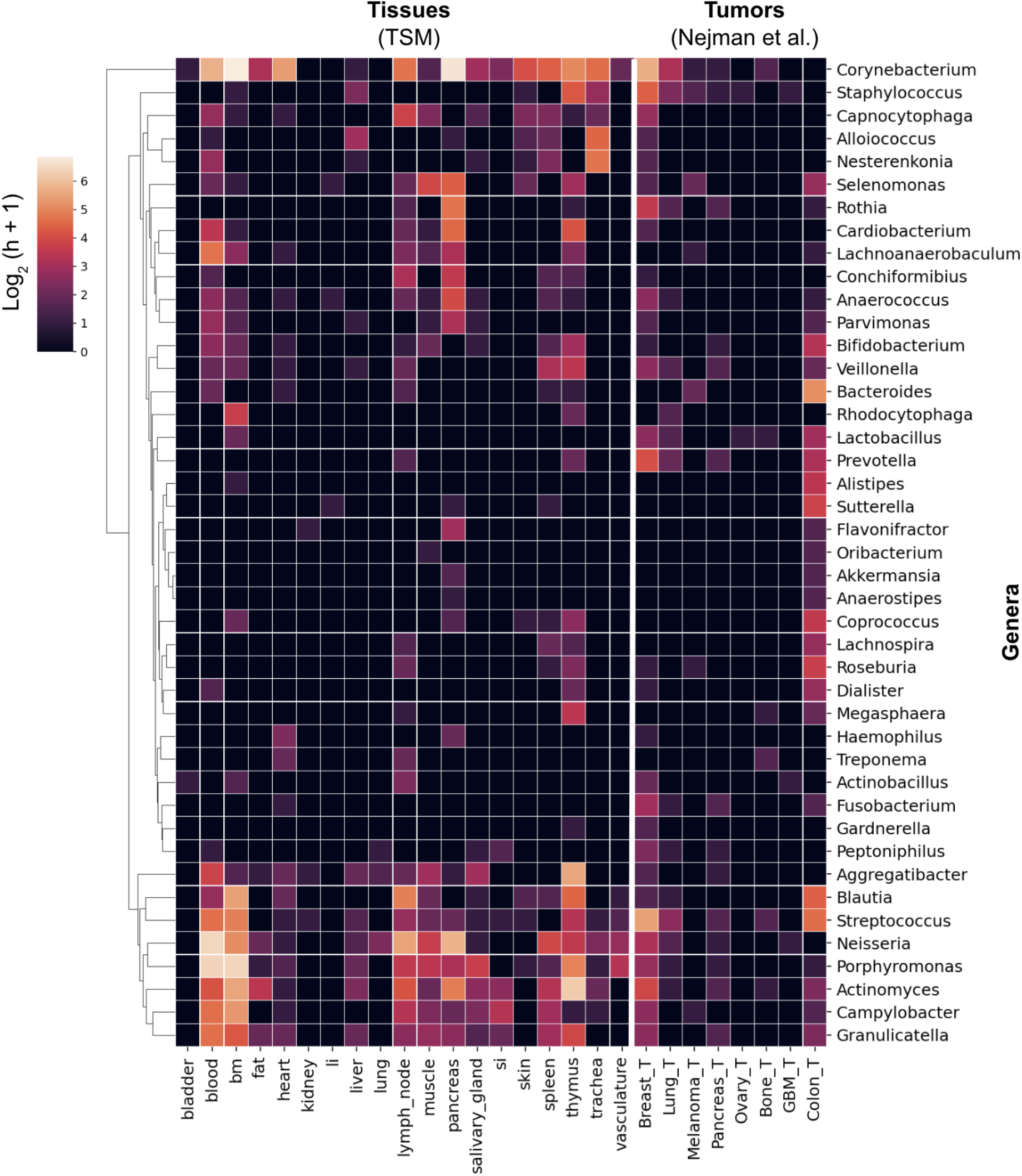
Abundance of bacterial genera previously found in tumor samples (Nejman *et. al*, 2020) and in the Tabula Sapiens Microbiome (TSM) F7 dataset stratified by tissue of origin. Abundance is based on log_2_ (hits +1). Samples with “_T” appended correspond to tumors. bm: bone marrow, li: large intestine, si: small intestine, GBM: glioblastoma multiforme.

**SI Figure 15.**
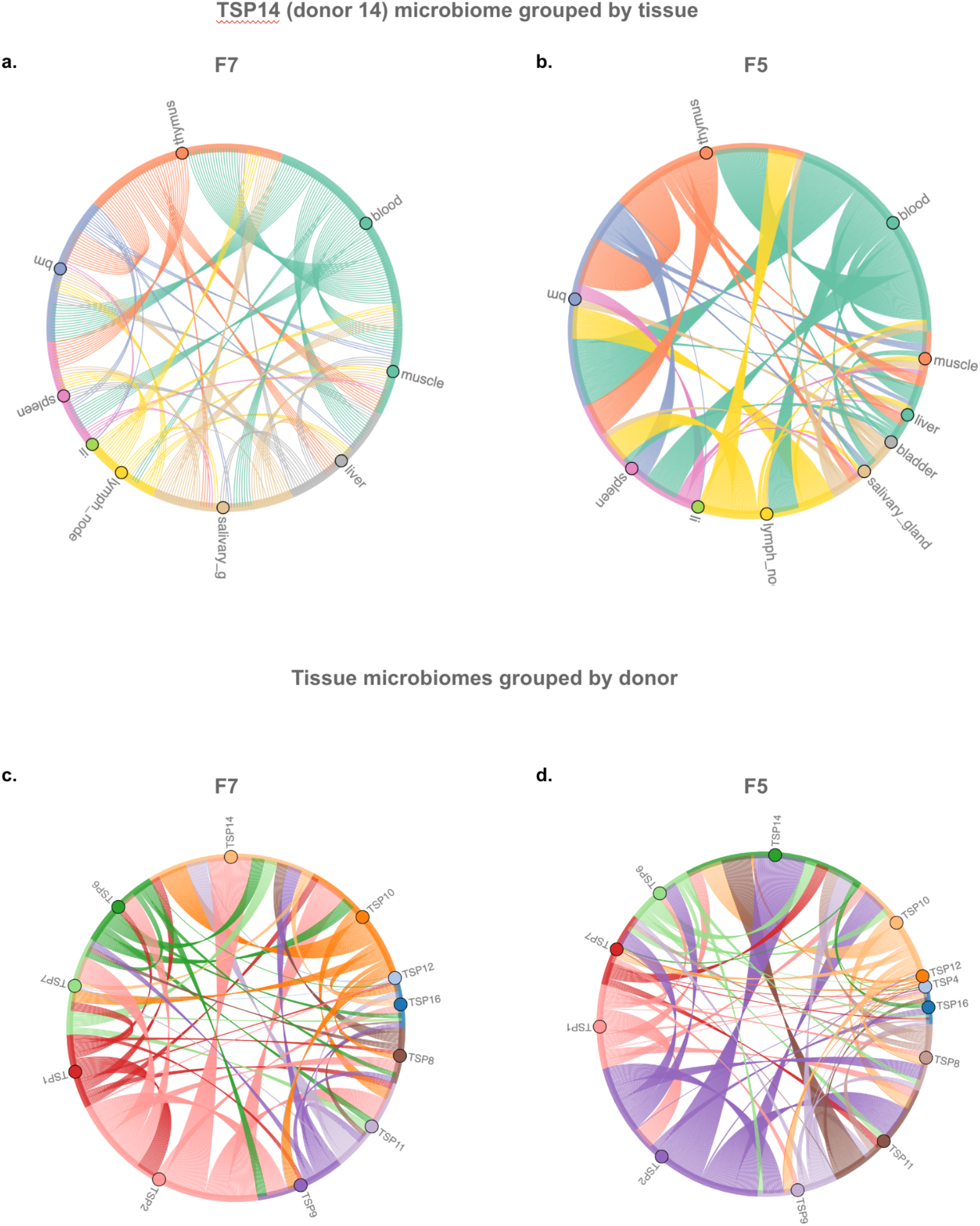
Circos plots of shared species within (a,b) and between donors (c,d). Each link represents a unique species. Species that are not shared are not shown. Left panels correspond to TSM F7 and the right panels correspond to TSM F5 datasets.

**SI Figure 16.**
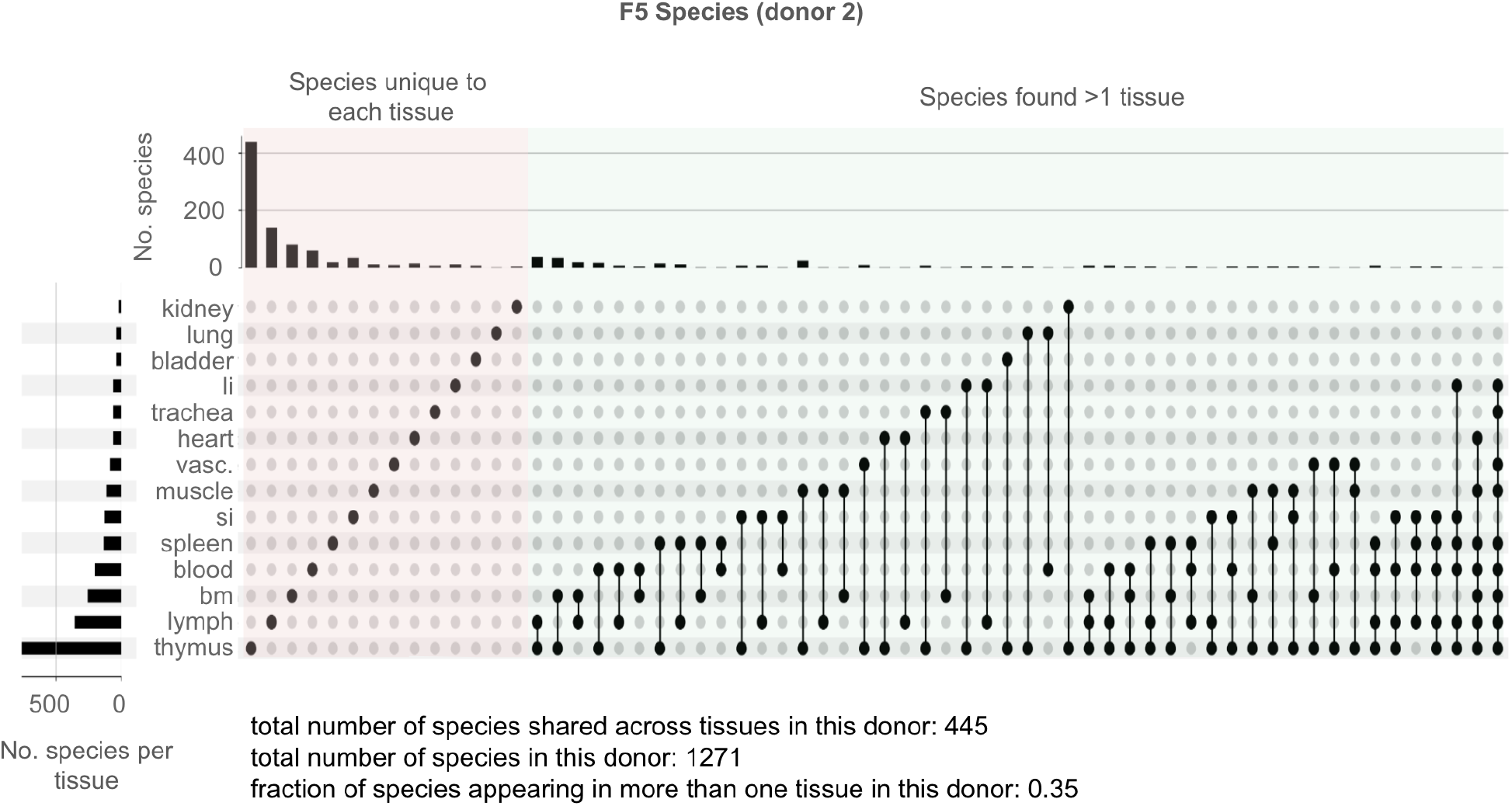
UpSet plots for TSP2 (donor 2) TSM F5 species demonstrating the number of shared taxa between tissues, as well as the number of taxa unique to each tissue. Unique sets of species are shown as filled circles. The intersection of two or more sets are shown via connected circles. Bars denote the size of each set or the intersection of sets. For visual clarity, the minimum subset size has been set to 2. bm: bone marrow, li: large intestine, si: small intestine, vasc: vasculature.

**SI Figure 17.**
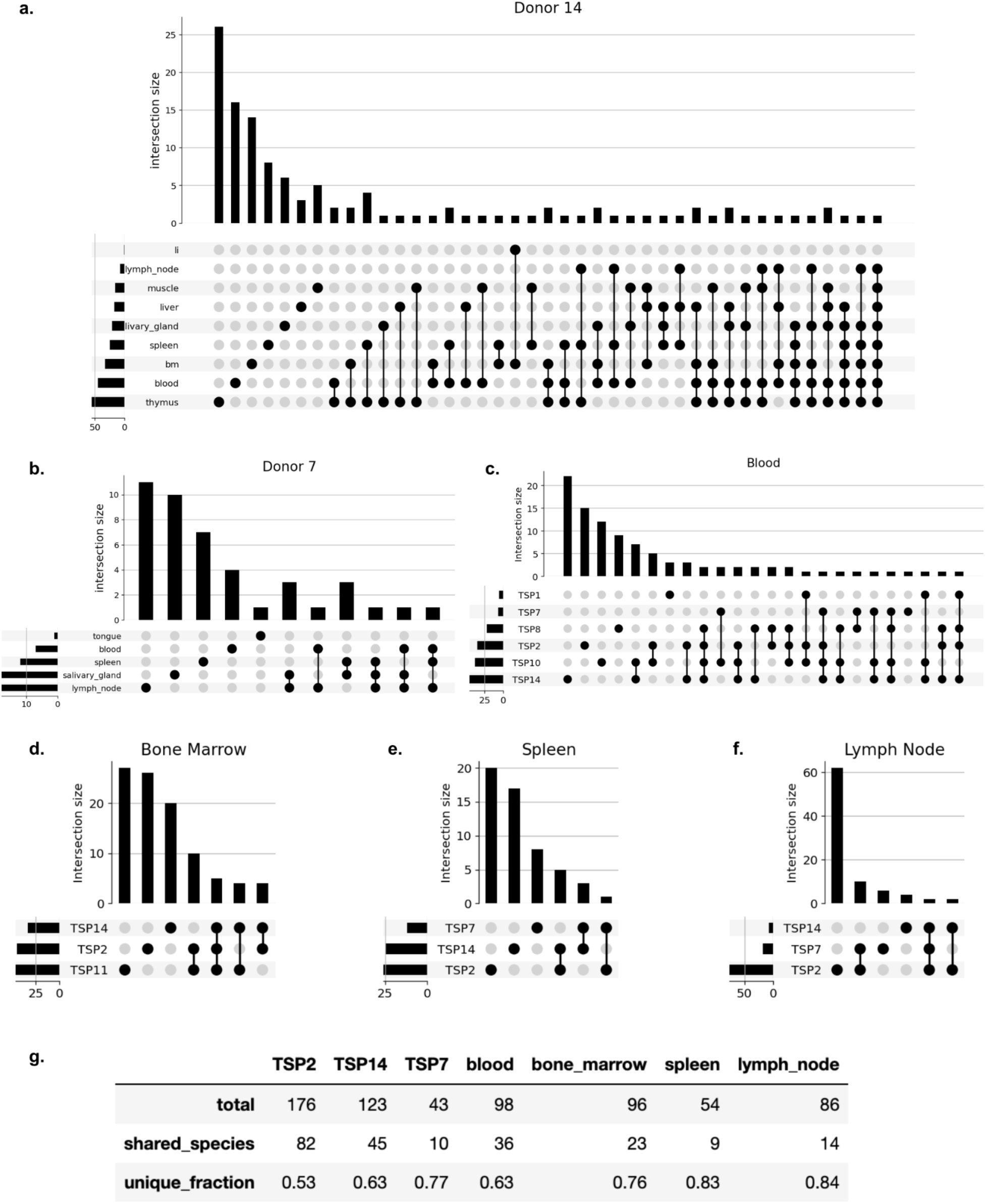
UpSet plots depicting the number of species shared and unique to each sample. Panels a and b depict intra-donor comparisons of tissues, while c-f depict inter-donor comparisons of the same tissue. Unique sets of species are shown as filled circles. The intersection of two or more sets are shown via connected circles. Bars denote the size of each set or the intersection of sets. Panel g summarizes the number and fraction of species that appear in more than one sample within each panel a-f.

**SI Figure 18.**
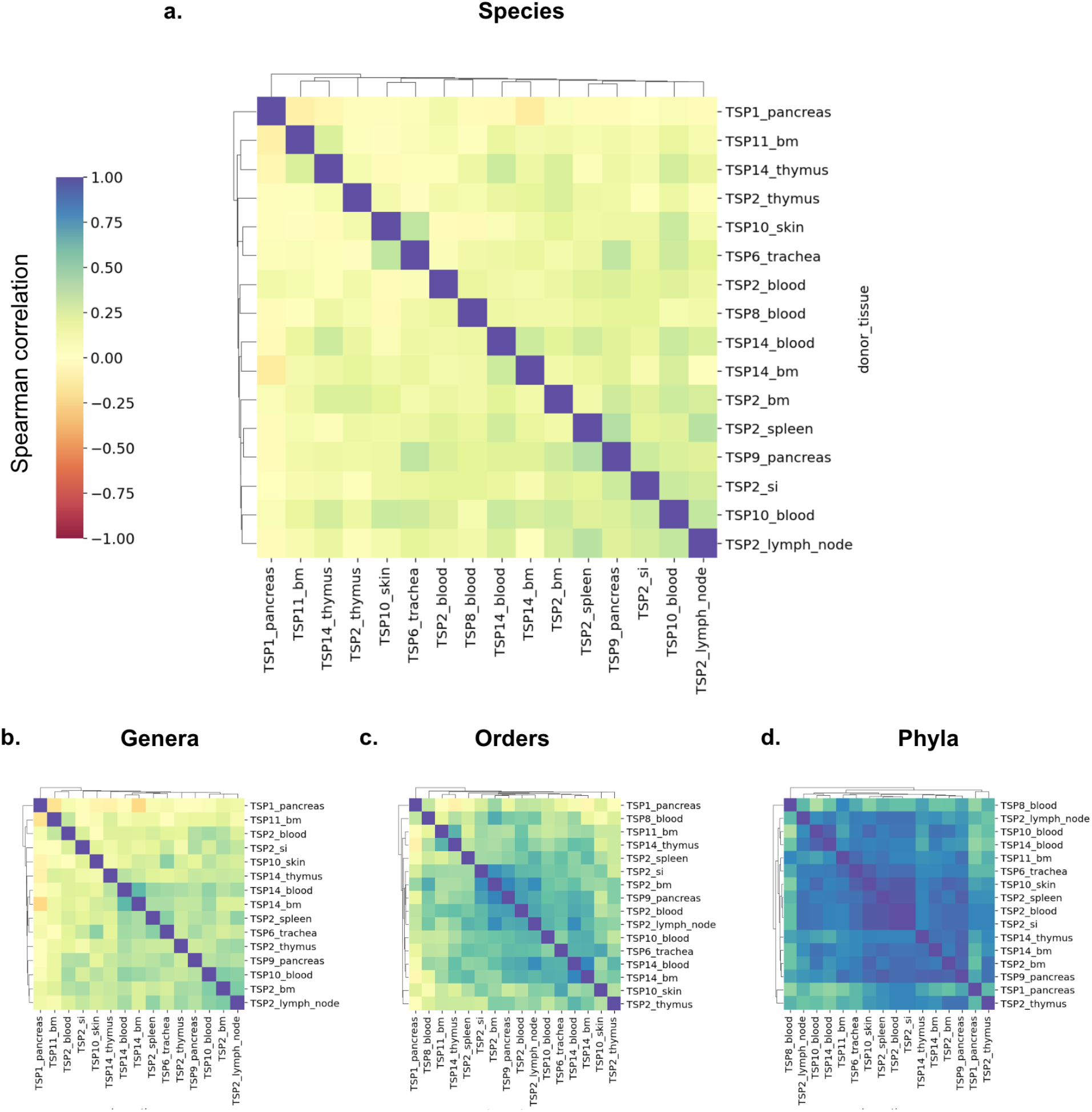
Tissue microbiomes across tissues and donors for TSM F7 bacterial dataset at the a) species, b) genera, c) orders and d) phyla-level classification. Pairwise Spearman correlation cluster maps across samples grouped by donor and tissue are shown. Only samples with more than 50 hits are shown.

**SI Figure 19.**
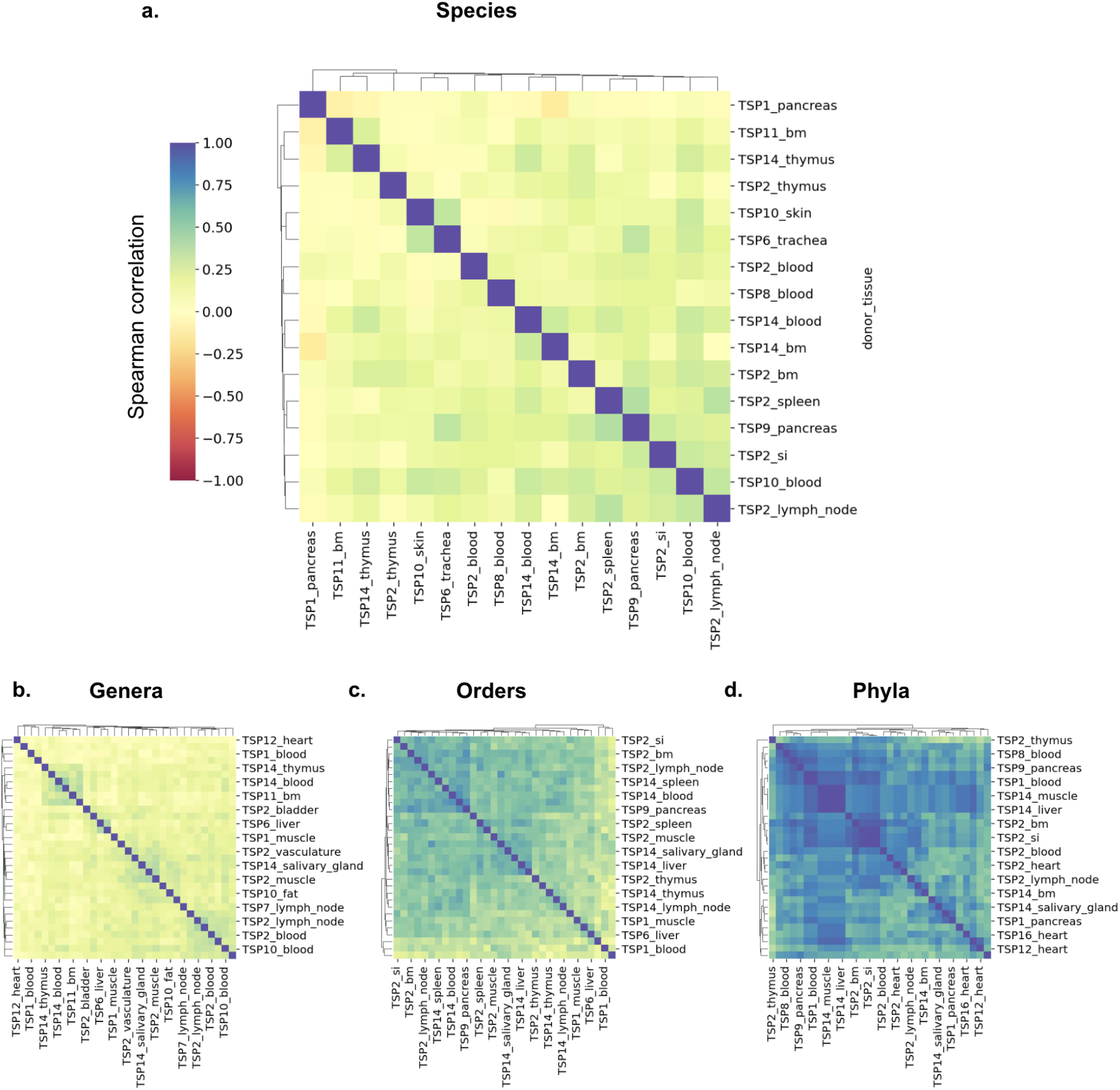
Tissue microbiomes across donor_tissue samples for TSM F5 bacterial dataset at the a) species, b) genera, c) orders and d) phyla-level classification. Pairwise Spearman correlation cluster maps across samples grouped by donor and tissue are shown. Only samples with more than 50 hits are shown.

**SI Figure 20.**
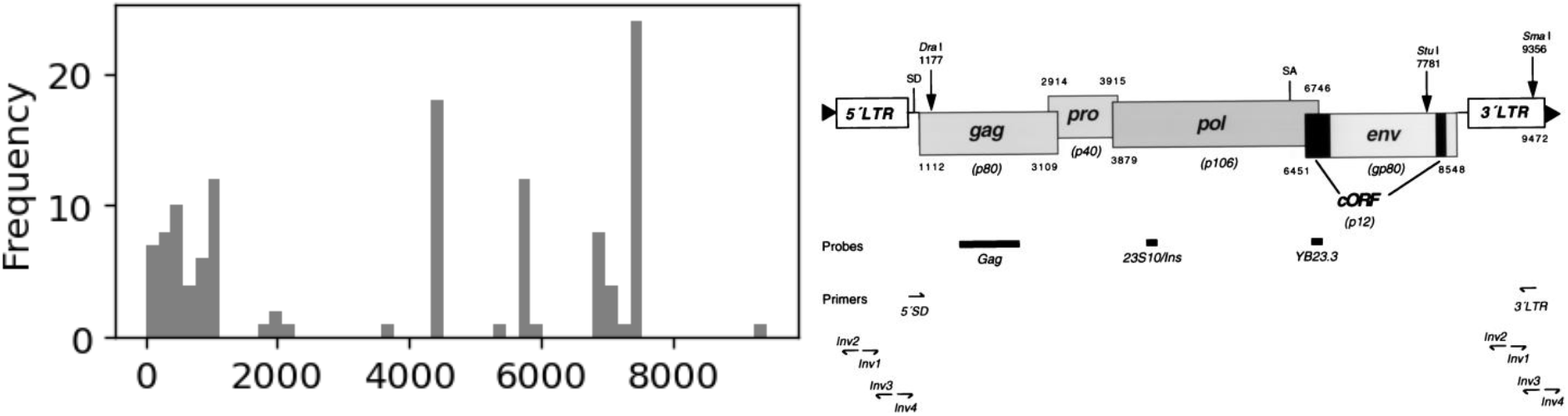
The genomic position of identified HERV K hits (left) with respect to the annotated HERV K genome (right) (Tӧnjes, 1999).

**SI Figure 21.**
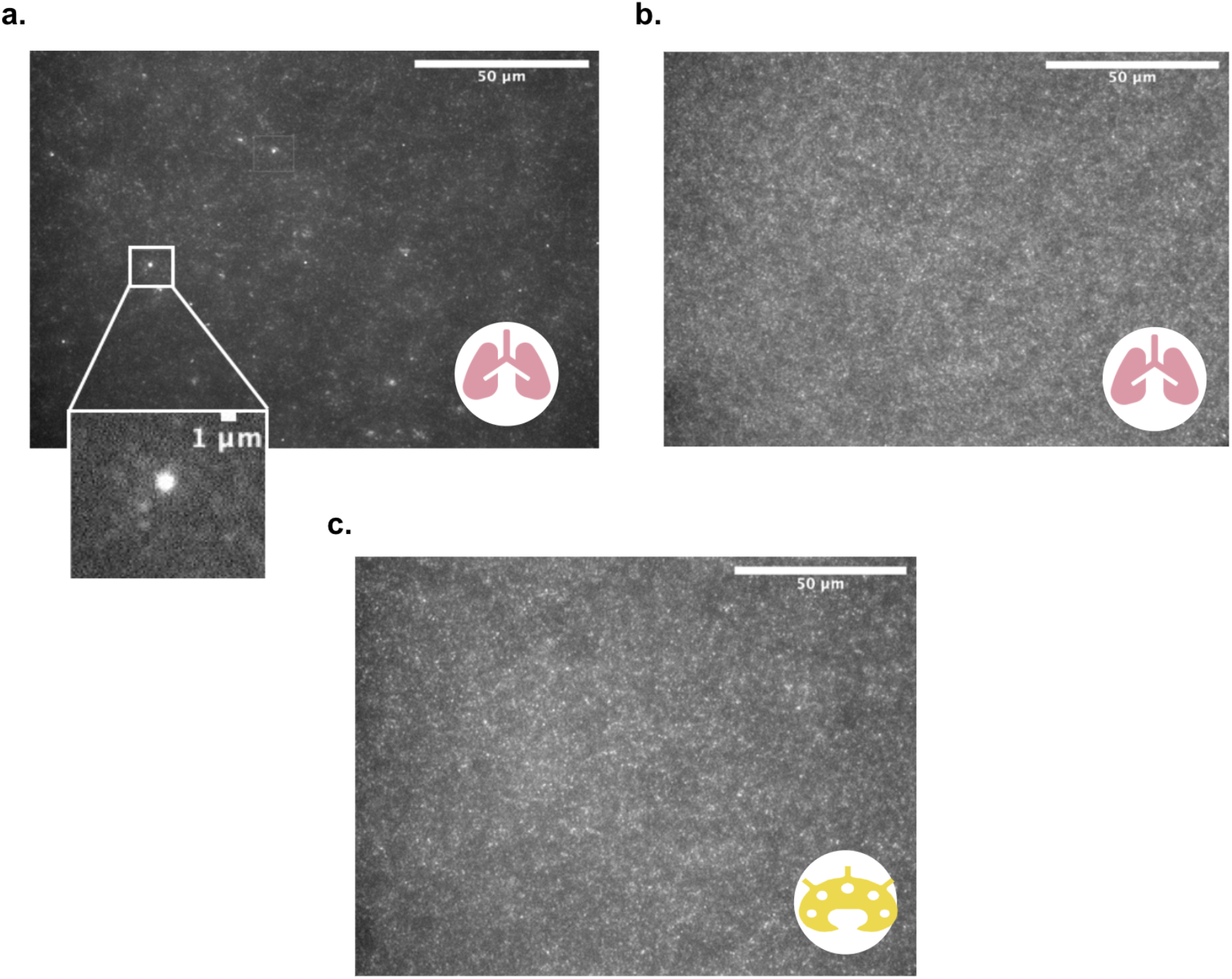
Fluorescence staining of donor-derived bacterial-like particles (BLPs) from the lung (a) and Virus-like particles (VLPs) from a lung and a lymph node sample (b,c). These images were used to estimate VLP and BLP concentrations in a cubic centimeter of tissue (∼10^8^ VLPs/cm^3^ and ∼10^7^ BLPs/cm^3^).

**SI Figure 22.**
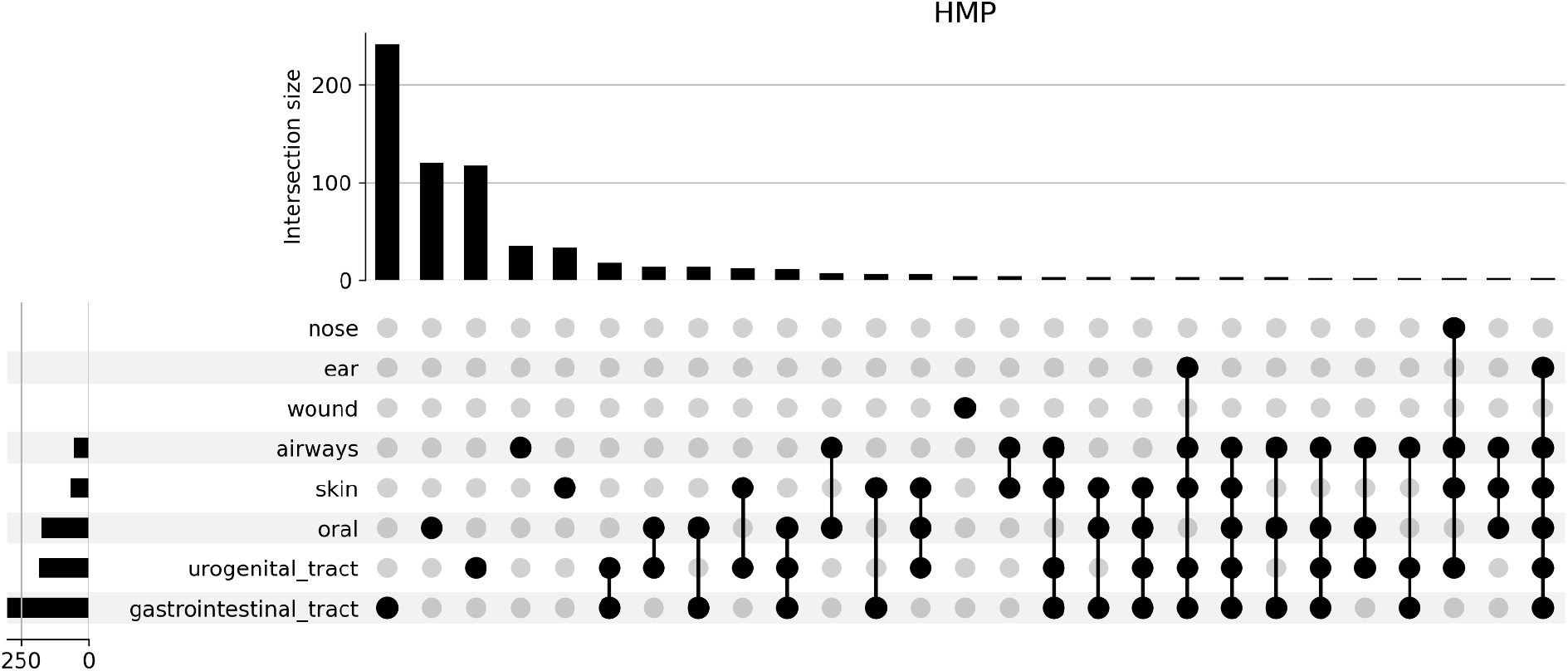
UpSet plot showing the taxa shared across different external-facing microbiomes in the Human Microbiome Project (HMP) dataset. Unique sets of taxa are shown as filled circles. The intersection of two or more sets are shown via connected circles. Bars denote the size of each set or the intersection of sets.

## SI Tables

**SI Table 1.** Precision and recall associated with SIMBA using a random selection of 100 viral genomes and 10 sequences per genome as a truth dataset.

|  | domain | phylum | class | order | family | genus | species |
| --- | --- | --- | --- | --- | --- | --- | --- |
| precision | 1.0 | 1.0 | 1.0 | 1.0 | 1.0 | 0.999 | 0.809 |
| recall | 1.0 | 1.0 | 1.0 | 1.0 | 1.0 | 1.000 | 1.000 |

**SI Table 2.** Examples of false positives associated with the viral truth dataset at the species level. “Species_truth” and “species_test” columns correspond to species found in the truth and test datasets, respectively.

|  | species_truth | species_test |
| --- | --- | --- |
| 0 | Acinetobacter virus AbP2 | Acinetobacter phage Bphi-R1888 |
| 1 | Acinetobacter virus D2 | Acinetobacter phage vB_AbaP_APK44 |
| 2 | Acinetobacter virus D2 | Acinetobacter phage vB_AbaP_D2M |
| 3 | Acinetobacter virus D2 | Acinetobacter phage vB_AbaP_PMK34 |
| 4 | Arthrobacter virus Piccoletto | Arthrobacter virus Beans |
| 5 | Campylobacter virus CP220 | Campylobacter phage CP20 |
| 6 | Campylobacter virus CP220 | Campylobacter virus CP21 |
| 7 | Campylobacter virus CP220 | Campylobacter virus IBB35 |
| 8 | Canna yellow mottle associated virus | Canna yellow mottle virus |
| 9 | Escherichia virus VR7 | Escherichia virus VR25 |
| 10 | Escherichia virus VR7 | Escherichia virus VR26 |
| 11 | Leuconostoc virus Lmd1 | Leuconostoc virus LN6B |
| 12 | Mycobacterium phage ArcherNM | Mycobacterium phage Changeling |
| 13 | Mycobacterium phage Phaedrus | Mycobacterium phage Akoma |
| 14 | Mycobacterium phage Phaedrus | Mycobacterium phage Compostia |
| 15 | Mycobacterium phage Tiffany | Mycobacterium phage Dieselweasel |
| 16 | Mycobacterium phage Tiffany | Mycobacterium phage P28Green |
| 17 | Mycobacterium phage Tiffany | Mycobacterium phage SoilDragon |
| 18 | Mycobacterium phage Tiffany | Mycobacterium virus Bxz2 |
| 19 | Mycobacterium virus Skinnyp | Mycobacterium phage ShrimpFriedEgg |
| 20 | Mycobacterium virus TA17a | Mycobacterium phage Holeinone |

**SI Table 3.**
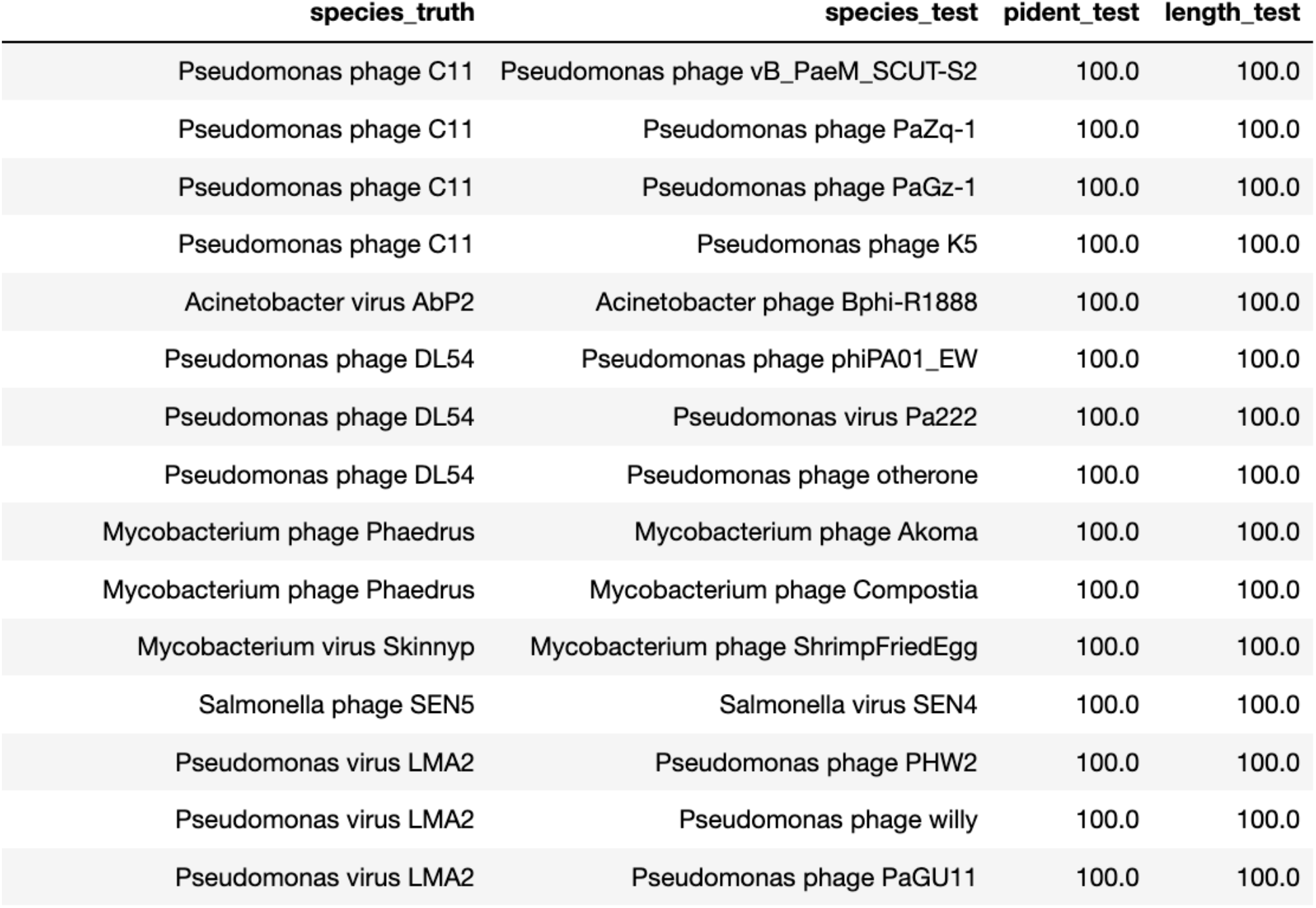
Examples of false positives associated with the viral truth dataset and the corresponding percent identity (“pident_test” column) and the length (“length_test” column) of the alignment between the input seq and output seq. “Species_truth” and “species_test” columns correspond to species found in the truth and test datasets, respectively.

**SI Table 4.** Precision and recall scores using a truth dataset containing 5000 sequences from 50 gut bacterial species (randomly selected sequences and species).

|  | domain | phylum | class | order | family | genus | species |
| --- | --- | --- | --- | --- | --- | --- | --- |
| **precision** | 1.000 | 0.999 | 0.997 | 0.996 | 0.973 | 0.947 | 0.699 |
| **recall** | 0.291 | 0.291 | 0.290 | 0.290 | 0.285 | 0.280 | 0.223 |

**SI Table 5.**
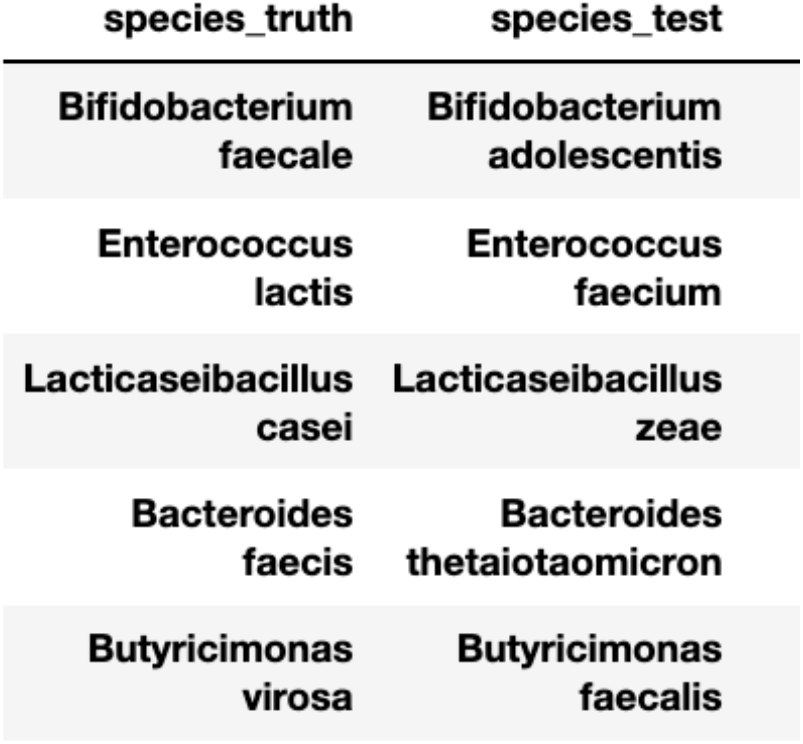
Species from the truth dataset and their corresponding false label at the level of species classification. First five false positives are shown. “Species_truth” and “species_test” columns correspond to species found in the truth and test datasets, respectively.

**SI Table 6.** Precision and recall scores using a truth dataset containing 5000 sequences from 50 oral bacterial species from the expanded Human Oral Microbiome Database (randomly selected sequences and species).

|  | domain | phylum | class | order | family | genus | species |
| --- | --- | --- | --- | --- | --- | --- | --- |
| **precision** | 1.000 | 1.000 | 0.996 | 0.990 | 0.987 | 0.974 | 0.854 |
| **recall** | 0.209 | 0.209 | 0.208 | 0.207 | 0.207 | 0.205 | 0.184 |

**SI Table 7.**
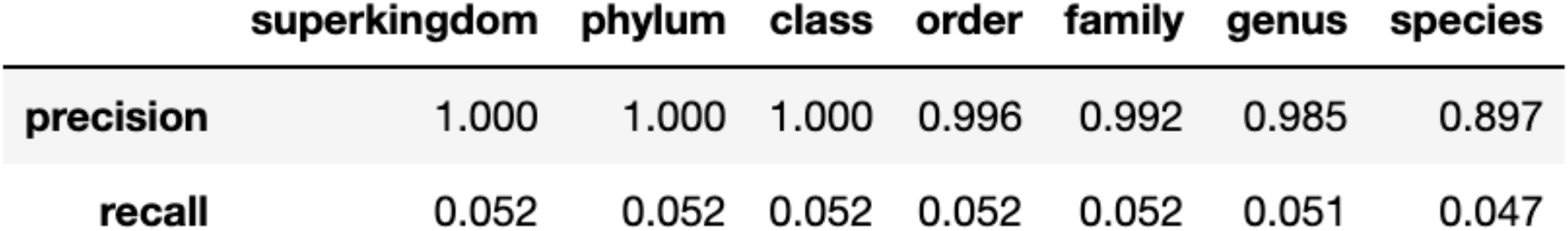
Precision and recall scores using a truth dataset containing 5000 sequences from the bacterial RefSeq database (based on 50 randomly selected species and 100 sequences from each).

|  | superkingdom | phylum | class | order | family | genus | species |
| --- | --- | --- | --- | --- | --- | --- | --- |
| **precision** | 1.000 | 1.000 | 1.000 | 0.996 | 0.992 | 0.985 | 0.897 |
| **recall** | 0.052 | 0.052 | 0.052 | 0.052 | 0.052 | 0.051 | 0.047 |

**SI Table 8.** Precision and recall scores using a truth dataset containing sequences from a fungal dataset^104^ (based on ∼5000 sequences).

|  | domain | phylum | class | order | family | genus | species |
| --- | --- | --- | --- | --- | --- | --- | --- |
| **precision** | 1.000 | 0.990 | 0.988 | 0.985 | 0.961 | 0.948 | 0.793 |
| **recall** | 0.207 | 0.205 | 0.205 | 0.204 | 0.200 | 0.198 | 0.171 |

**SI Table 9.**
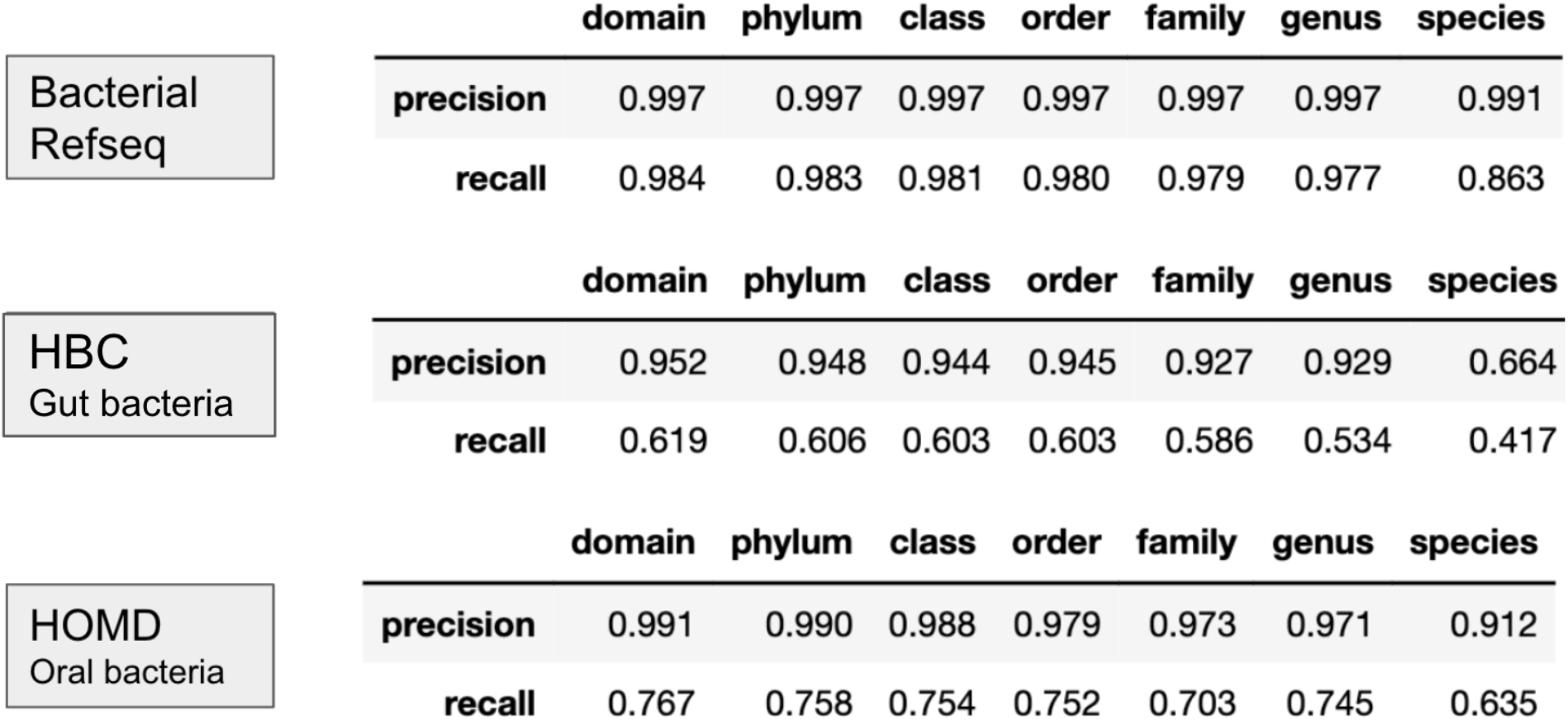
Kraken2 precision and recall scores using 3 truth datasets each containing 5000 sequences (50 randomly selected species and 100 sequences from each) from the bacterial RefSeq database, Human Gastrointestinal Bacteria Culture Collection (HBC) and the expanded Human Oral Microbiome Database (HOMD).

**SI Table 10.** Testing the precision of Kraken2 (against nt) classification assignments for a Tabula Sapiens Microbiome sample (Bladder, donor 2). Mean precision and standard deviation for three batches of input sequences selected from the classified reads (100 randomly selected sequences per batch from each domain, excluding Archaea).

| Mean Precision +/- SD |  |  |  |  |  |  |  |
| --- | --- | --- | --- | --- | --- | --- | --- |
|  | domain | phylum | class | order | family | genus | species |
| **Eukaryotic** | 0.77 ± 0.04 | 0.72 ± 0.07 | 0.61 ± 0.06 | 0.6 ± 0.06 | 0.57 ± 0.06 | 0.54 ± 0.07 | 0.54 ± 0.07 |
| **Bacterial** | 0.06 ± 0.01 | 0.06 ± 0.01 | 0.06 ± 0.01 | 0.06 ± 0.01 | 0.06 ± 0.01 | 0.06 ± 0.01 | 0.05 ± 0.01 |
| **Viral** | 0.03 ± 0.01 | 0.03 ± 0.01 | 0.03 ± 0.02 | 0.03 ± 0.02 | 0.03 ± 0.02 | 0.04 ± 0.03 | 0.03 ± 0.02 |

**SI Table 11.**
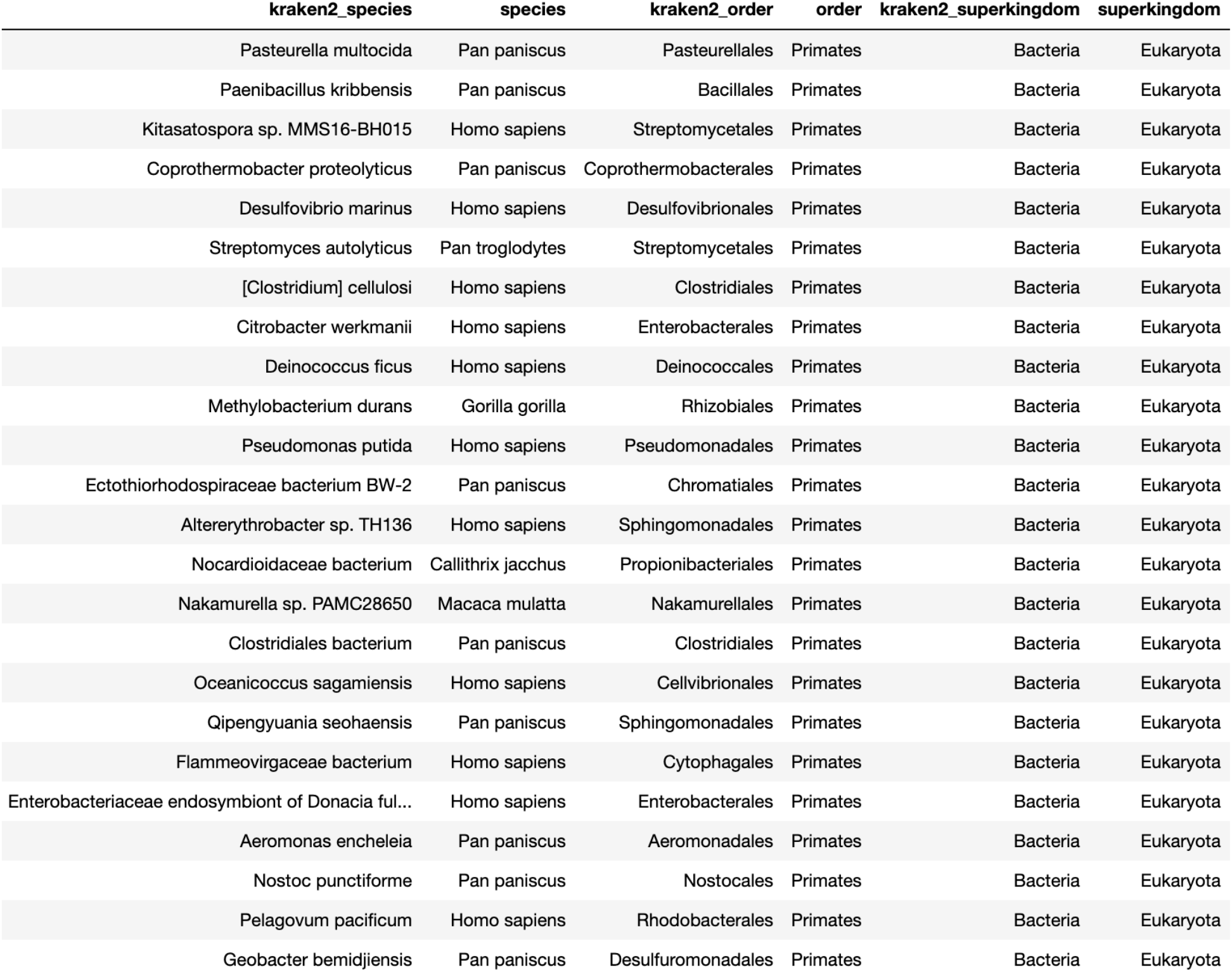
Examples of the type of reads that are misclassified as bacterial by Kraken2 (against nt) and their classification based on BLASTn (against nt).

**SI Table 12**. List of reagents and instruments per tissue used to generate negative control samples.

**SI Table 13**. Absolute counts and fraction of total microbial hits, unique hits (based on UMIs), species, genera, cell-containing droplets with hits, and empty droplets with hits remaining after each filter is applied.

**SI Table 14.** 95% AUROC confidence intervals for different machine learning models tested on unseen data. “auroc_test_95perc_ci_lower” and “auroc_test_95perc_ci_upper” columns correspond to the lower and upper bounds for the 95% confidence interval, respectively.

|  | **auroc_test_95perc_ci_lower** | **auroc_test_95perc_ci_upper** |
| --- | --- | --- |
| **gradient boosting classifier** | 0.82 | 0.836 |
| **random forest** | 0.785 | 0.807 |
| **multi-layer perceptron** | 0.765 | 0.797 |
| **support vector rbf** | 0.733 | 0.767 |
| **knn** | 0.71 | 0.758 |
| **logistic regression** | 0.703 | 0.727 |

**SI Table 15.** 95% AUPR confidence intervals for different machine learning models tested on unseen data. “aupr_test_95perc_ci_lower” and “aupr_test_95perc_ci_upper” columns correspond to the lower and upper bounds for the 95% confidence interval, respectively.

|  | **aupr_test_95perc_ci_lower** | **aupr_test_95perc_ci_upper** |
| --- | --- | --- |
| **gradient boosting classifier** | 0.748 | 0.776 |
| **random forest** | 0.715 | 0.753 |
| **multi-layer perceptron** | 0.69 | 0.736 |
| **support vector rbf** | 0.668 | 0.702 |
| **knn** | 0.636 | 0.69 |
| **logistic regression** | 0.63 | 0.66 |

**SI Table 16**. Evaluating Davinci model’s level of self-consistency to the same input prompt. Three independent outputs are listed for each species. The temperature parameter was set to default (0.5).

**SI Table 17**. Davinci model outputs for 100 test species using the GEM dataset as the truth dataset. “davinci_0”, “davinci_1”, “davinci_2” columns report the three independent outputs of the Davinci model for each species. “Davinci_human” is set to 0 if a given species is found in humans by at least one of the three Davinci outputs, and 1 otherwise. “Davinci_human_3x” is set to 0 if all three Davinci outputs report that a given species has been found in humans. The “gem_human” column is the truth dataset, set to 0 if a given species is found in humans in the GEM dataset, and 1 otherwise.

**SI Table 18**. Davinci model outputs for 1500+ species whose habitats were not identified by the HUE datasets. “davinci_0”, “davinci_1”, “davinci_2” columns report the three independent outputs of the Davinci model for each species. “habitat_assignment” column is set to 0 if all three outputs state the presence of a given species in the human body, and set to 1 otherwise.

**SI Table 19.** Fraction of microbial hits in the TSM F5 and F7 datasets that have taxonomic assignment at different levels of taxonomic categorization.

|  | index | f5 | f7 |
| --- | --- | --- | --- |
| 0 | species | 1.00 | 1.00 |
| 1 | genus | 0.88 | 0.90 |
| 2 | family | 0.92 | 0.94 |
| 3 | order | 0.96 | 0.95 |
| 4 | class | 0.97 | 0.97 |
| 5 | phylum | 0.98 | 0.99 |
| 6 | domain | 1.00 | 1.00 |

**SI Table 20**. The table underlying the flow paths from **Figure 2.a** (Sankey plot). Columns: source, dest (destination), values. Source and destination columns represent the nodes in the Sankey plot, while the values demonstrate the weight applied onto the connecting edges. The suffixes “_phy”, “_ord”, “_tis”, “_tum”, “_cou”, “_hmp”, “_hab” are denoting phyla, orders, tissues, Nejman *et al.* tumor types, UHGG countries, HMP external facing sites, and GEM habitats, respectively. bm: bone marrow, li: large intestine, si: small intestine.

**SI Table 21**. Genera found in tumor microbiomes that are not found in the TSM F7 or TSM F5 datasets.

**SI Table 22**. The table underlying the flow paths from **SI File 2** (Sankey plot). Columns: tumor_type (Nejman *et al.*), genus and tissue (TSM F7), hmp_site (HMP), Country (UHGG), habitat2 (GEM). The suffixes are: “_phy”, “_ord”, “_tis”, “_tum”, “_cou”, “_hmp”, “_hab” are denoting phyla, orders, tissues, Nejman *et al.* tumor types, UHGG countries, HMP external facing sites, and GEM habitats, respectively. bm: bone marrow, li: large intestine, si: small intestine.

**SI Table 23**. Statistics for the distribution of unique hits per cell (“cell_umi” column) and empty droplets (“empty_umi column) in the TSM F7 dataset.

**SI Table 24**. Outlier cells with their associated species and domains, as well as their tissue, donor and cell type of origin. BM: Bone Marrow.

**SI Table 25**. Relative fraction of outlier cells and empty droplets from TSM F7, stratified by the domains where the hits map to. All outlier viral hits from cells and empty droplets map to EBV.

**SI Table 26**. Outlier cell types in TSM F7 dataset. Fraction and total number of cells in each cell type that had F7 hits are shown. Only cell types with a minimum of 10 cells sequenced are included.

**SI Table 27**. HERV K (HML-2) transcripts across donors, spanning a diverse range of tissues and cell types.

## SI Files

**SI File 1.** The HTML file that corresponds to **Figure 2.a** (Sankey plot). This file can be opened using a browser for an interactive experience.

**SI File 2.** The HTML file of a Sankey plot showing the genera shared between several datasets: TSM F7, GEM, UHGG, Nejman et al.). This file can be opened using a browser. Columns from left to right are the same as those from the main Sankey plot (**Figure 2.a**). The suffixes are: “_phy”, “_ord”, “_tis”, “_tum”, “_cou”, “_hmp”, “_hab” are denoting phyla, orders, tissues, Nejman *et al.* tumor types, UHGG countries, HMP external facing sites, and GEM habitats, respectively. bm: bone marrow, li: large intestine, si: small intestine.

**SI File 3.** The HTML file for the network plot shown in **Figure 3.b**. Please see **Figure 3.b** caption for more information about node and edge weights.

**SI File 4.** The HTML file for EBV Sankey plot shown in **Figure 4.a**. The columns from left to right correspond to donors, tissues, cell types and compartments. The suffixes for these columns are “_don”, “_tis”, “_cel”, and “_com”. bm: bone marrow.

## The Tabula Sapiens Consortium Author List

### Overall Project Direction and Coordination

Robert C. Jones^1^, Jim Karkanias^2^, Mark Krasnow^3, 4^, Angela Oliveira Pisco^2^, Stephen R. Quake^1, 2, 5^, Julia Salzman^3, 6^, Nir Yosef^2, 7–9^

### Donor Recruitment

Bryan Bulthaup^10^, Phillip Brown^10^, William Harper^10^, Marisa Hemenez^10^, Ravikumar Ponnusamy^10^, Ahmad Salehi^10^, Bhavani A. Sanagavarapu^10^, Eileen Spallino^10^

### Surgeons

Ksenia A. Aaron^11^, Waldo Concepcion^10^, James M. Gardner^12, 13^, Burnett Kelly^10, 14^, Nikole Neidlinger^10^, Zifa Wang^10^

### Logistical coordination

Sheela Crasta^1, 2^, Saroja Kolluru^1, 2^, Maurizio Morri2, Angela Oliveira Pisco^2^, Serena Y. Tan^15^, Kyle J. Travaglini^3^, Chenling Xu^7^

### Organ Processing

Marcela Alcántara-Hernández^16^, Nicole Almanzar^17^, Jane Antony^18^, Benjamin Beyersdorf^19^, Deviana Burhan^20^, Kruti Calcuttawala^21^, Matthew M. Carter^16^, Charles K. F. Chan^18, 22^, Charles A. Chang^23^, Stephen Chang^3, 19^, Alex Colville^21, 24^, Sheela Crasta^1, 2^, Rebecca N. Culver^25^, Ivana Cvijović^1, 5^, Gaetano D’Amato^26^, Camille Ezran^3^, Francisco X. Galdos^18^, Astrid Gillich^3^, William R. Goodyer^27^, Yan Hang^23, 28^, Alyssa Hayashi^1^, Sahar Houshdaran^29^, Xianxi Huang^19, 30^, Juan C. Irwin^29^, SoRi Jang^3^, Julia Vallve Juanico^29^, Aaron M. Kershner^18^, Soochi Kim^21, 24^, Bernhard Kiss^18^, Saroja Kolluru^1, 2^, William Kong^18^, Maya E. Kumar^17^, Angera H. Kuo^18^, Rebecca Leylek^16^, Baoxiang Li^31^, Gabriel B. Loeb^32^, Wan-Jin Lu^18^, Sruthi Mantri^33^, Maxim Markovic^1^, Patrick L. McAlpine^11, 34^, Antoine de Morree^21, 24^, Maurizio Morri^2^, Karim Mrouj^18^, Shravani Mukherjee^31^, Tyler Muser^17^, Patrick Neuhöfer^3, 35, 36^, Thi D. Nguyen^37^, Kimberly Perez^16^, Ragini Phansalkar^26^, Angela Oliveira Pisco^2^, Nazan Puluca^18^, Zhen Qi18, Poorvi Rao^20^, Hayley Raquer-McKay^16^, Nicholas Schaum^18, 21^, Bronwyn Scott^31^, Bobak Seddighzadeh^38^, Joe Segal^20^, Sushmita Sen^29^, Shaheen Sikandar^18^, Sean P. Spencer^16^, Lea Steffes^17^, Varun R. Subramaniam^31^, Aditi Swarup^31^, Michael Swift^1^, Kyle J. Travaglini^3^, Will Van Treuren^16^, Emily Trimm^26^, Stefan Veizades^19, 39^, Sivakamasundari Vijayakumar^18^, Kim Chi Vo^29^, Sevahn K. Vorperian^1, 40^, Wanxin Wang^29^, Hannah N.W. Weinstein^38^, Juliane Winkler^41^, Timothy T.H. Wu^3^, Jamie Xie^38^, Andrea R.Yung^3^, Yue Zhang^3^

### Sequencing

Angela M. Detweiler^2^, Honey Mekonen^2^, Norma F. Neff^2^, Rene V. Sit^2^, Michelle Tan^2^, Jia Yan^2^

### Histology

Gregory R. Bean^15^, Vivek Charu^15^, Erna Forgó^15^, Brock A. Martin^15^, Michael G. Ozawa^15^, Oscar Silva^15^, Serena Y. Tan^15^, Angus Toland^15^, Venkata N.P. Vemuri^2^

### Data Analysis

Shaked Afik^7^, Kyle Awayan^2^, Rob Bierman^3, 6^, Olga Borisovna Botvinnik^2^, Ashley Byrne^2^, Michelle Chen^1^, Roozbeh Dehghannasiri^3, 6^, Angela M. Detweiler^2^, Adam Gayoso^7^, Alejandro A Granados^2^, Qiqing Li^2^, Gita Mahmoudabadi^1^, Aaron McGeever^2^, Antoine de Morree^21, 24^, Julia Eve Olivieri^3, 6, 42^, Madeline Park^2^, Angela Oliveira Pisco^2^, Neha Ravikumar^1^, Julia Salzman^3, 6^, Geoff Stanley^1^, Michael Swift^1^, Michelle Tan^2^, Weilun Tan^2^, Alexander J Tarashansky^2^, Rohan Vanheusden^2^, Sevahn K. Vorperian^1, 40^, Peter Wang^3, 6^, Sheng Wang^2^, Galen Xing^2^, Chenling Xu^6^, Nir Yosef^2, 6–8^

### Expert Cell Type Annotation

Marcela Alcántara-Hernández^16^, Jane Antony^18^, Charles K. F. Chan^18, 22^, Charles A. Chang^23^, Alex Colville^21, 24^, Sheela Crasta^1, 2^, Rebecca Culver^25^, Les Dethlefsen^43^, Camille Ezran^3^, Astrid Gillich^3^, Yan Hang^23, 28^, Po-Yi Ho^16^, Juan C. Irwin^29^, SoRi Jang^3^, Aaron M. Kershner^18^, William Kong^18^, Maya E Kumar^17^, Angera H. Kuo^18^, Rebecca Leylek^16^, Shixuan Liu^3, 44^, Gabriel B. Loeb^32^, Wan-Jin Lu^18^, Jonathan S Maltzman^45, 46^, Ross J. Metzger^27, 47^, Antoine de Morree^21, 24^, Patrick Neuhöfer^3, 35, 36^, Kimberly Perez^16^, Ragini Phansalkar^26^, Zhen Qi^18^, Poorvi Rao^20^, Hayley RaquerMcKay^16^, Koki Sasagawa^19^, Bronwyn Scott^31^, Rahul Sinha^15, 18, 35^, Hanbing Song^38^, Sean P. Spencer^16^, Aditi Swarup^31^, Michael Swift^1^, Kyle J. Travaglini^3^, Emily Trimm^26^, Stefan Veizades^19, 39^, Sivakamasundari Vijayakumar^18^, Bruce Wang^20^, Wanxin Wang^29^, Juliane Winkler^41^, Jamie Xie^38^, Andrea R.Yung^3^

### Tissue Expert Principal Investigators

Steven E. Artandi^3, 35, 36^, Philip A. Beachy^18, 23, 48^, Michael F. Clarke^18^, Linda C. Giudice^29^, Franklin W. Huang^38, 49^, Kerwyn Casey Huang^1, 16^, Juliana Idoyaga^16^, Seung K Kim^23, 28^, Mark Krasnow^3, 4^, Christin S. Kuo^17^, Patricia Nguyen^19, 39, 46^, Stephen R. Quake^1, 2, 5^, Thomas A. Rando^21, 24^, Kristy Red-Horse^26^, Jeremy Reiter^50^, David A. Relman^16, 43, 46^, Justin L. Sonnenburg^16^, Bruce Wang^20^, Albert Wu^31^, Sean M. Wu^19, 39^, Tony Wyss-Coray^21, 24^

### Affiliations

^1^Department of Bioengineering, Stanford University; Stanford, CA, USA.

^2^Chan Zuckerberg Biohub; San Francisco, CA, USA.

^3^Department of Biochemistry, Stanford University School of Medicine; Stanford, CA, USA.

^4^Howard Hughes Medical Institute; USA.

^5^Department of Applied Physics, Stanford University; Stanford, CA, USA.

^6^Department of Biomedical Data Science, Stanford University; Stanford, CA, USA.

^7^Center for Computational Biology, University of California Berkeley; Berkeley, CA, USA.

^8^Department of Electrical Engineering and Computer Sciences, University of California Berkeley; Berkeley, CA, USA.

^9^Ragon Institute of MGH, MIT and Harvard; Cambridge, MA, USA.

^10^Donor Network West; San Ramon, CA, USA.

^11^Department of Otolaryngology-Head and Neck Surgery, Stanford University School of Medicine; Stanford, California, USA.

^12^Department of Surgery, University of California San Francisco; San Francisco, CA, USA.

^13^Diabetes Center, University of California San Francisco; San Francisco, CA, USA.

^14^DCI Donor Services; Sacramento, CA, USA.

^15^Department of Pathology, Stanford University School of Medicine; Stanford, CA, USA.

^16^Department of Microbiology and Immunology, Stanford University School of Medicine; Stanford, CA, USA.

^17^Department of Pediatrics, Division of Pulmonary Medicine, Stanford University; Stanford, CA, USA.

^18^Institute for Stem Cell Biology and Regenerative Medicine, Stanford University School of Medicine; Stanford, CA, USA.

^19^Department of Medicine, Division of Cardiovascular Medicine, Stanford University; Stanford, CA, USA.

^20^Department of Medicine and Liver Center, University of California San Francisco; San Francisco, CA, USA.

^21^Department of Neurology and Neurological Sciences, Stanford University School of Medicine; Stanford, CA, USA.

^22^Department of Surgery - Plastic and Reconstructive Surgery, Stanford University School of Medicine; Stanford, CA, USA.

^23^Department of Developmental Biology, Stanford University School of Medicine, Stanford, CA, USA

^24^Paul F. Glenn Center for the Biology of Aging, Stanford University School of Medicine; Stanford, CA, USA.

^25^Department of Genetics, Stanford University School of Medicine; Stanford, CA, USA.

^26^Department of Biology, Stanford University; Stanford, CA, USA.

^27^Department of Pediatrics, Division of Cardiology, Stanford University School of Medicine; Stanford, CA, USA.

^28^Stanford Diabetes Research Center, Stanford University School of Medicine, Stanford, California

^29^Center for Gynecology and Reproductive Sciences, Department of Obstetrics, Gynecology and Reproductive Sciences, University of California San Francisco; San Francisco, CA, USA.

^30^Department of Critical Care Medicine, The First Affiliated Hospital of Shantou University Medical College; Shantou, China.

^31^Department of Ophthalmology, Stanford University School of Medicine; Stanford, CA, USA.

^32^Division of Nephrology, Department of Medicine, University of California San Francisco; San Francisco, CA, USA.

^33^Stanford University School of Medicine; Stanford, CA, USA.

^34^Mass Spectrometry Platform, Chan Zuckerberg Biohub; Stanford, CA, USA.

^35^Stanford Cancer Institute, Stanford University School of Medicine; Stanford, CA, USA.

^36^Department of Medicine, Division of Hematology, Stanford University School of Medicine, Stanford, CA, USA

^37^Department of Biochemistry and Biophysics, Cardiovascular Research Institute, University of California San Francisco; San Francisco, CA, USA.

^38^Division of Hematology and Oncology, Department of Medicine, Bakar Computational Health Sciences Institute, Institute for Human Genetics, University of California San Francisco; San Francisco, CA, USA.

^39^Stanford Cardiovascular Institute; Stanford CA, USA.

^40^Department of Chemical Engineering, Stanford University; Stanford, CA, USA.

^41^Department of Cell & Tissue Biology, University of California San Francisco; San Francisco, CA, USA.

^42^Institute for Computational and Mathematical Engineering, Stanford University; Stanford, CA, USA.

^43^Division of Infectious Diseases & Geographic Medicine, Department of Medicine, Stanford University School of Medicine; Stanford, CA, USA.

^44^Department of Chemical and Systems Biology, Stanford University School of Medicine, Stanford, CA, USA

^45^Division of Nephrology, Stanford University School of Medicine; Stanford, CA, USA.

^46^Veterans Affairs Palo Alto Health Care System; Palo Alto, CA, USA.

^47^Vera Moulton Wall Center for Pulmonary and Vascular Disease, Stanford University School of Medicine; Stanford, CA, USA.

^48^Department of Urology, Stanford University School of Medicine, Stanford, CA, USA

^49^Division of Hematology/Oncology, Department of Medicine, San Francisco Veterans Affairs Health Care System, San Francisco, CA, USA.

^50^Department of Biochemistry, University of California San Francisco; San Francisco, CA, USA.

## Notes

### Competing Interest Statement

The authors have declared no competing interest.

### Summary of Updates

New bioinformatic filters have been applied on the dataset. New external databases have been integrated. Additional benchmarking of the computational pipeline has been performed.

